# Structural Mechanisms of DNAJC13 Dimeric Assembly and InsP6 binding in Recycling Endosome Regulation

**DOI:** 10.64898/2026.07.28.740790

**Authors:** Tao Fu, Chan Lee, Frances V. Hundley, Joao A. Paulo, J. Wade Harper

## Abstract

The balance of plasma membrane protein degradation and recycling during endocytosis is regulated, in part, by the large J-domain-containing protein DNAJC13/RME-8 and WASH complexes, which function together with Retromer to support cargo trafficking into recycling endosomes. Despite extensive cellular and biochemical studies, the structural basis and proteomic landscape of DNAJC13 function remain elusive. Here, we find that DNAJC13 forms an unexpected antiparallel homodimeric architecture involving distinctive symmetrical interactions between composite IWN1 and α-solenoid ARM2 domains in each protomer. Additionally, two PH-like domains of unknown function (PHL2 and PHL3), adjacent to the PI(3)P-binding PHL1 domain, form an unanticipated composite, positively charged pocket occupied by InsP_6_, as visualized structurally and verified by mass spectrometry. Proteomic profiling of DNAJC13-associated endosomes revealed enrichment of recycling endosomal components and WASH complexes. Mutations disrupting the dimer interface disable recruitment of WASH complexes to endosomes and result in elongated endosomal tubulation. Disruption of InsP_6_ binding impairs DNAJC13 binding to PI(3)P-containing vesicles in vitro and Transferrin-positive endosomes in cells. We further demonstrate that DNAJC13 dimerization and InsP_6_ binding promote melanin production during melanosome maturation, a process known to require recycling endosomes. This work provides a structural and mechanistic framework for understanding DNAJC13 function in recycling endosome control.

## INTRODUCTION

Plasma membrane (PM) protein dynamics are regulated in part by the endolysosomal system, a network of membrane-bound organelles.^1–3^ Endocytic vesicles formed at the PM transition into RAB5-positive compartments known as early or sorting endosomes—referred to here simply as “early endosomes.” Early endosomes act as hubs for sorting and recycling of PM proteins and also acquire membrane and lumenal endolysosomal factors via fusion with transport vesicles originating at the Golgi.^4,5^ PM proteins are sorted vectorially within the early endosome. Proteins destined for lysosomal degradation are sorted into membrane domains that undergo ESCRT-driven invagination to form intralumenal vesicles (ILVs), thereby facilitating PM protein degradation during maturation to lysosomes.^2,6^ In contrast, proteins that are destined for recycling to either the PM or to the trans-Golgi network are sorted into more specialized recycling endosomes, marked by RAB11 and often localized to the perinuclear endocytic recycling compartment (ERC), in a process that involves cargo-sorting machinery including Retromer, Retriever, CCC complex, the Wiskott-Aldrich syndrome protein and SCAR homolog (WASH) complex, and sorting nexins (SNXs). These proteins function together to promote fission of recycling tubules and cargo sorting from the early endosome.^1,7,8^ Disruption of endosomal recycling machinery has been linked to a broad spectrum of neurodegenerative disorders, including Alzheimer’s disease, Parkinson’s disease, and Ritscher-Schinzel syndrome. The highly dynamic state of organelles within the endolysosomal system may represent a continuum of states with nanodomains on individual vesicles simultaneously undergoing distinct trafficking events as the vesicle matures toward the fully degradative lysosomal state.^9^

Among the factors that participate in recycling endosome biogenesis is DNAJC13 (also called RME-8 in *C. elegans* and *D. melanogaster*).^10,11^ *DNAJC13*, which is found across Eukarya with the exception of fungi,^10,11^ encodes a 250 kDa multi-domain protein containing a central ∼70 residue J-domain, a conserved feature of the HSP40 class of chaperones that promotes substrate recruitment and ATP hydrolysis by HSP70 chaperones.^12,13^ In addition, DNAJC13 contains an N-terminal PH domain that preferentially binds phosphatidylinositol-3 phosphate (PI(3)P)^14^, which is enriched on early endosomes,^15^ and extended α-solenoid domains interspersed by four ∼90 residue IWN repeats, named for the conserved Ile-Trp-Asn tripeptide.^10^ In *C. elegans*, RME-8 is thought to function together with SNX-1 to facilitate removal of clathrin from recently budded endosomes, and RME-8 mutants result in accumulation of clathrin on endosomes accompanied by re-routing of PM cargo to degradation rather than to retrograde recycling.^16^ In this context, the J-domain is proposed to promote HSP70-dependent disassembly of clathrin from the endosome.^17^ Beyond roles in clathrin removal, DNAJC13/RME-8 also regulates the balance between recycling and degradative trafficking by facilitating the formation of distinct degradative and recycling microdomains on endosomes.^18^ First, DNAJC13 has been proposed to remove ESCRT-0 complexes from endosomal sub-compartments to limit degradative domains.^11,16^ Second, DNAJC13 functions together with Retromer complexes to establish recycling microdomains that allow sorting of PM cargo into a recycling vesicle.^11,16^ This latter function is accomplished, in part, via association with SNX-1 and WASH complexes. In cells lacking DNAJC13/RME-8, endosomal tubules accumulate and are enriched in SNX-1, Retromer and its cargos, including CI-MPR and GLUT1.^19^ In contrast, another study indicates that loss of DNAJC13 reduced the extent of lysosomal turnover of specific GPCRs, suggesting a positive role in trafficking of some cell surface proteins for degradation.^20^ Interestingly, a rare missense variant N855S in DNAJC13 was reported to segregate with an autosomal dominant, late-onset PD with Lewy body pathology in a large family,^21,22^ but studies in diverse ethnic groups indicate that this variant is rare across patient cohorts.^23–25^ Nevertheless, DNAJC13/RME-8 mutations in model systems promotes α-synuclein pathology and alterations in endolysosomal system components, ^20,26,21,27^ both pathways known to be associated with PD.^28,29^ Our understanding of DNAJC13/RME-8 cellular function and its interactions with endosomal components such as the WASH complex has been based primarily on co-immunoprecipitation using truncation mutants and imaging-based co-localization studies with marker proteins in mammalian cells or *C.elegans*.^11,19^ As such, structural mechanisms of DNAJC13 assembly and membrane recruitment, as well as a proteomic description of DNAJC13-associated endosomes, remain lacking.

Here, we use single-particle cryo-electron microscopy to determine the structure of DNAJC13 (1-1486), leading to two unanticipated discoveries. First, we find that DNAJC13 uses unique structural elements within the IWN1-ARM2 region to form a symmetrical arch-shaped antiparallel homodimer. DNAJC13 dimer interface mutants form monomers in vitro and in cells and reduce the ability of DNAJC13 to rescue endosomal tubulation observed in DNAJC13 knockout cells. Second, we discovered that two tandem N-terminal PH-like domains (PHL2 and PHL3) of unknown function form an unexpected composite, positively charged pocket that binds inositol hexakisphosphate (InsP_6_), as visualized structurally and confirmed by mass spectrometry. Disruption of InsP_6_ binding impairs recruitment of DNAJC13 to PI(3)P-containing membranes via its PI(3)P-binding PHL1 domain in vitro and reduces DNAJC13 localization to endosomes in cells. Additionally, to directly profile DNAJC13-associated endocytic compartments, we developed DNAJC13 as an endosome purification handle. Proteomic analysis of DNAJC13-associated endosomal populations revealed a bias for components of recycling endosomes and WASH complexes when directly compared with analogous EEA1-associated endosomal populations. Importantly, the endosomal population associated with dimerization defective DNAJC13 displays selective loss of components of the WASH complex, indicating that DNAJC13 dimerization promotes WASH recruitment. Finally, we confirmed that DNAJC13 is required for melanin production during melanosome maturation, consistent with a previously established role for recycling endosomes,^30^ and demonstrate that this activity depends on both InsP_6_ binding and homodimerization. Together, these studies provide a structural and proteomic framework that uncovers and validates two novel aspects of DNAJC13 function in recycling endosome regulation.

## RESULTS

### Cryo-EM structure of a DNAJC13 Dimer

To investigate structural mechanisms underlying DNAJC13 function, we expressed an N-terminally FLAG-tagged DNAJC13(1-2191) construct in Expi293F cells and purified by affinity capture using anti-FLAG resin followed by size-exclusion chromatography. This construct, which lacks a predicted unstructured C-terminal region (residues 2192-2243), eluted as a homogeneous peak by size-exclusion chromatography together with associated HSP70, as assessed by SDS-PAGE and mass spectrometry (**Figure S1A**). Mass photometry indicated a major peak with a mass matching the expected size of a DNAJC13 homodimer, with an additional peak matching the expected size of a dimer plus HSP70 (**Figure S1B**). The identification of a dimeric species was unanticipated. This preparation was used for single-particle cryo-EM (**Data S1.1 and Table S1**). Extensive 2D and 3D classification and refinement yielded a consensus reconstruction at 2.4 Å global resolution (**Data S1.1-S1.2**). To probe conformational heterogeneity, we performed 3D variability analysis followed by focused 3D classification and local refinement around the N- and C-terminal regions of both protomers, resulting in four locally refined maps at 2.5-2.7 Å resolution (**Data S1.1**). These four maps together with the consensus map were combined to generate a composite reconstruction used for model building and refinement (**Data S1.1**) and validation statistics were obtained against the consensus map (**Table S1**).

Well-resolved density extends from the N-terminal PH-like (PHL1) domain to approximately residue 1486, with two DNAJC13 molecules in the complex. Densities for the C-terminal IWN3 to ARM5 region and the poorly conserved residues 671-722 are not seen in the cryo-EM map, reflecting apparent flexibility in these regions (**Figures 1A-C**). The overall structure of the DNAJC13 dimer when viewed perpendicular to its C2 symmetry axis resembles an “arch” with the N-termini of both protomers forming the base (**Figures 1C-F**). When viewed from the C2 symmetry axis, two distinct dimer interfaces are observed, which due to symmetry, reflect three interaction interfaces (**Figures 1D and 1F**), as described in detail below. Comparison of the cryo-EM model with AlphaFold2-predicted^31,32^ full-length monomer DNAJC13, aligned based on the N-terminal 400 residues, revealed that the orientation of the C-terminal region is essentially reversed relative to the experimental structure (**Figure S1C**). Similarly, AlphaFold3^33^ predictions of both monomeric and dimeric DNAJC13 produced C-terminal arrangements that differ markedly from the cryo-EM structure (**Figures S1D and S1E**), further supporting substantial conformational variability in the C-terminal region (**Movie S1**). While the folds of individual sub-domains in the AlphaFold-predicted structures agree well with the experimentally determined structure (**Data S1.3**), the AlphaFold3-predicted structure of dimeric DNAJC13(1-1486) deviates from the cryo-EM structure (RMSD = 2.472 Å), reflecting, in part, the flexibility of the N-terminal PHL1-3 domains, consistent with our cryo-EM analysis (**Figure S1F, Movie S1**). We therefore used our cryo-EM map to generate a composite full-length DNAJC13 homodimer model. First, an AlphaFold3-predicted DNAJC13(1-1486) dimer was used as the initial model for refinement against the cryo-EM map, retaining regions with weak or absent density. Second, an AlphaFold2-predicted model of the C-terminal region (residues 1292-2243) was aligned to the refined structure, and the overlapping residues (1292-1486) were removed, yielding a seamless composite model spanning residues 1-2243. Finally, the coordinates of PI3P or PI(3,5)P_2_ were determined by aligning the AlphaFold3-predicted PI3P- or PI(3,5)P_2_-DNAJC13 complexes to the composite model. This hybrid model illustrates how the extended C-terminal IWN/ARM repeats may project from the central arch and also provides a plausible mechanism for dual interactions with PI(3)P in the membrane through the base of the arch (**Figures 1G and 1H, Figure S1G**). Importantly, representative 2D class averages provide support for extension of the C-terminal arms well beyond the regions of each promoter that were resolved with atomic resolution (**Figure 1I**).

**Figure 1.**
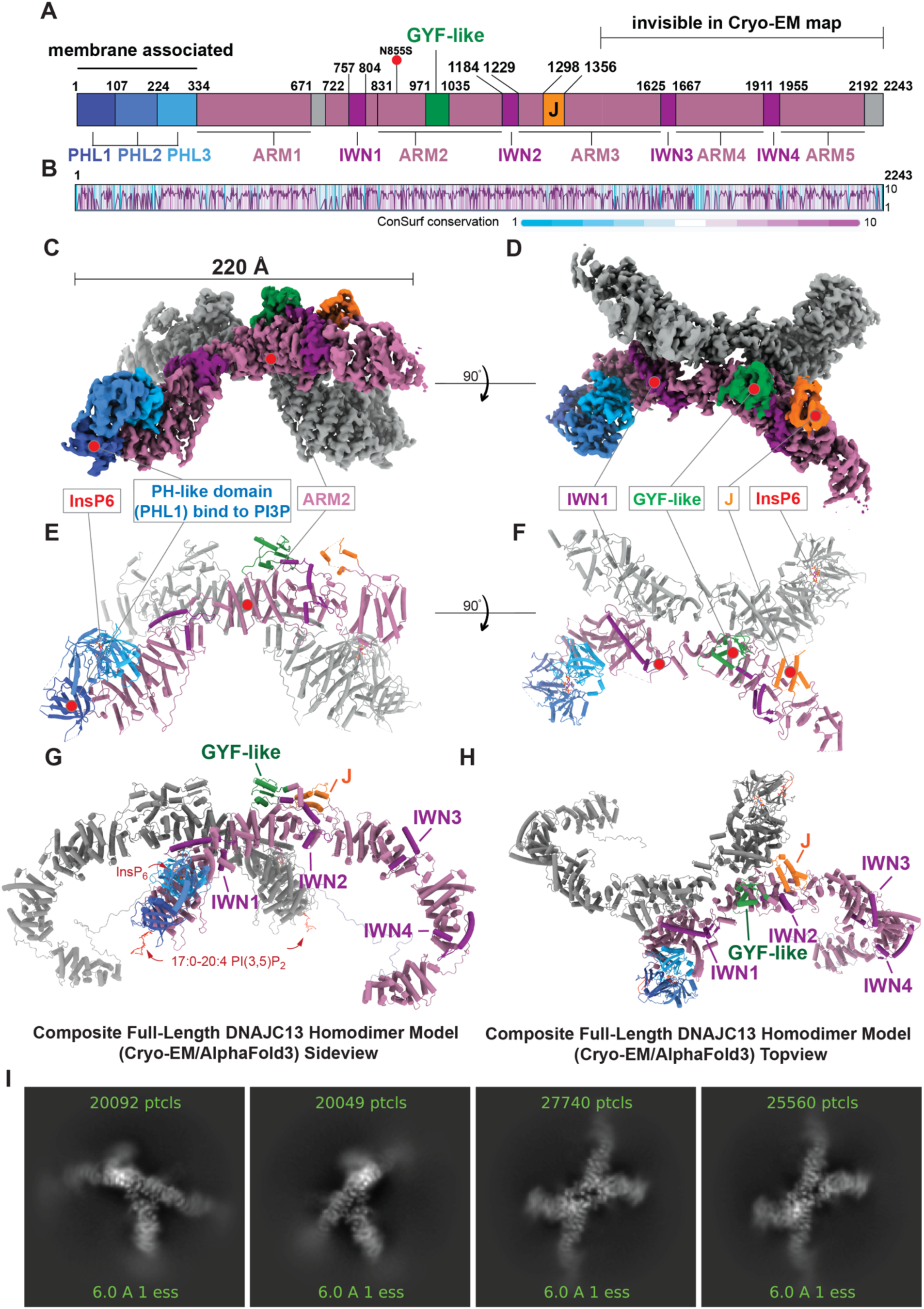
Structural architecture of DNAJC13 by cryo-EM. (A) Schematic of human DNAJC13 domain organization. The N-terminal PH-like domains (PHL1-3; deep/medium/light blue), five ARM domains (ARM1-5; mauve) and four IWN repeats (IWN1-4; purple) are shown together with the GYF-like domain (green) and J domain (orange). The position of Parkinson’s disease-associated mutation is indicated above the schematic. Regions not resolved in the cryo-EM map (residues 671-722) and the predicted disordered region (residues 2192-2243) are colored grey. Residues 1487-2191, encompassing the C-terminal IWN3-ARM5 region not visible in cryo-EM map, are indicated at the top. (B) ConSurf conservation analysis mapped onto the DNAJC13 primary sequence, with highly conserved residues shown in darker magenta and less conserved residues in darker blue. (C-D) Cryo-EM density map of DNAJC13(1-1486) shown in two orthogonal views, with the density of one protomer segmented and colored according to the domain scheme in (A). The overall length of the elongated assembly is about 220 Å. (E-F) Cryo-EM model of DNAJC13(1-1486) dimer shown in cartoon representation with the same color scheme and orientations as in (C) and (D). Positions of InsP_6_, the PHL1 domain, IWN1, ARM2, the GYF-like domain, and the J domain are indicated. (G-I) Composite full-length DNAJC13 homodimer model incorporating PI(3,5)P_2_ and InsP_6_. Side (Panel G) and top (Panel H) views are shown. This hybrid model illustrates how the extended C-terminal IWN/ARM repeat array may project from the dimer center, providing a plausible architecture for full-length dimeric DNAJC13/InsP6/PI(3,5)P_2_ at membranes. The structural model is colored according to the domain schematic in Figure 1A. 17:0-20:4 PI(3,5)P_2_ is shown in red. (Panel I) Representative 2D class average supporting the presence of the flexible C-terminal region observed in the composite model.

### Unique Structural Features within DNAJC13

Beyond the N-terminal PHL-domains, the domain organization of DNAJC13 features extended α-solenoid-like repeats interspersed with IWN, J, and GYF-like domains (**Figure 1A; Data S1.3-S1.4**). Our cryo-EM map together with AlphaFold predictions allowed us to establish a detailed understanding of DNAJC13 domain structure, including the identification of features that appear to be unique among structures currently represented in the PDB. First, each IWN repeat is comprised of two helices linked by two short anti-parallel β-sheets, which adopt a unique fold found in DNAJC13 orthologs, as assessed by FoldSeek^34^ (**Data S1.3**). The conserved IWN motif lies at the transition from a short β-strand into a ∼20-residue helix, with the tryptophan side chain buried between two helices within the IWN repeat. This 20-residue helix within the IWN domain packs against the helices from the preceding α-solenoid module and is connected by a short unstructured loop with the next α-solenoid domain (**Figure S1H**). Second, the α-solenoid repeats contain 12-15 helical segments, and in some cases display conformations similar to repeat units in the armadillo-repeat protein β-catenin (**Data S1.3**). We therefore refer to these segments as ARM domains 1-5 (**Figure 1A**). Third, the three N-terminal PH-like domains (PHL1-3) form a compact tri-domain module wherein the pockets of PHL2 and PHL3 face each other and are not solvent exposed, in contrast with PHL1 (**Figures 1C, 1E and 1G**). Fourth, FoldSeek identified a GYF-like fold within the ARM2 domain and confirmed the presence of a canonical J domain within ARM3 (**Figure 1E-H, Data S1.4**). The GYF domain of GIGYF2 contains a hydrophobic groove that binds a PPGL motif^35^; the DNAJC13 GYF-like domain similarly presents a hydrophobic patch that may also engage hydrophobic peptide motifs (**Data S1.4**). Although the J-domain is resolved in the structure, we did not detect particles containing HSP70 through the particle classification approach described above (see **Data S1.1**). Nevertheless, immobilized DNAJC13 J-domain (residues 1300-1355) recombinantly expressed and isolated from *E. coli* selectively captured multiple HSP70 family members as binding partners from Expi293F lysates (**Data S1.4**), consistent with its known interaction in cells.^14^ The structure of dimeric DNAJC13 provides a framework for dissecting functional components involved in endosomal trafficking.

### Composite InsP_6_ Binding Site Formed by PHL2-PHL3

PHL1 is reported to bind preferentially to PI(3)P (and to a lesser extent PI(3,5)P_2_) and thereby recruit DNAJC13 to endosomal membranes^14^, a finding we validated with both full-length DNAJC13 and DNAJC13(1-1486) (**Figure S2A-D**). AlphaFold3 predicts binding of PI(3)P- or PI(3,5)P_2_-phospholipids using a conserved basic pocket in PHL1, with key interactions for the 3-phosphate group of the inositide provided by highly conserved residues K17 and R26, and W20 (**Figure S2E, Data S2.1**).^14^ Given the similar sequence and fold of PHL2 and PHL3 relative to PHL1 (**Data S2.1**), as also observed in recent AlphaFold models^11,36^, we initially hypothesized that PHL2 and PHL3 might also bind one or two PI(3)P molecules. In initial experiments, we found that deletion of each PHL domain abolished DNAJC13 co-localization with Transferrin-positive endosomes, to an extent similar to that seen with the PHL1 point mutant W20A (**Data S2.2-S2.3**). However, during DNAJC13 model building, we unexpectedly observed strong density within the presumptive phospho-inositide-binding pockets of PHL2 and PHL3 matching closely the size and shape expected for InsP_6_ (**Figure 2A, Figure S2F-I**). InsP_6_ is an abundant soluble metabolite^37,38^ that can act as a structural cofactor in proteins.^39^ In our structure, K118, H120, R178, and H180 in PHL2, together with K242, R246, H247, K276, and R304 in PHL3, form a positively charged composite pocket that makes hydrogen bonds with each phosphate group of InsP_6_ (**Figure 2B, Figure S2F; Data S2.1**). This extensive interaction network suggests that InsP_6_ functions as a “molecular glue” that holds PHL2 and PHL3 in close apposition.

**Figure 2.**
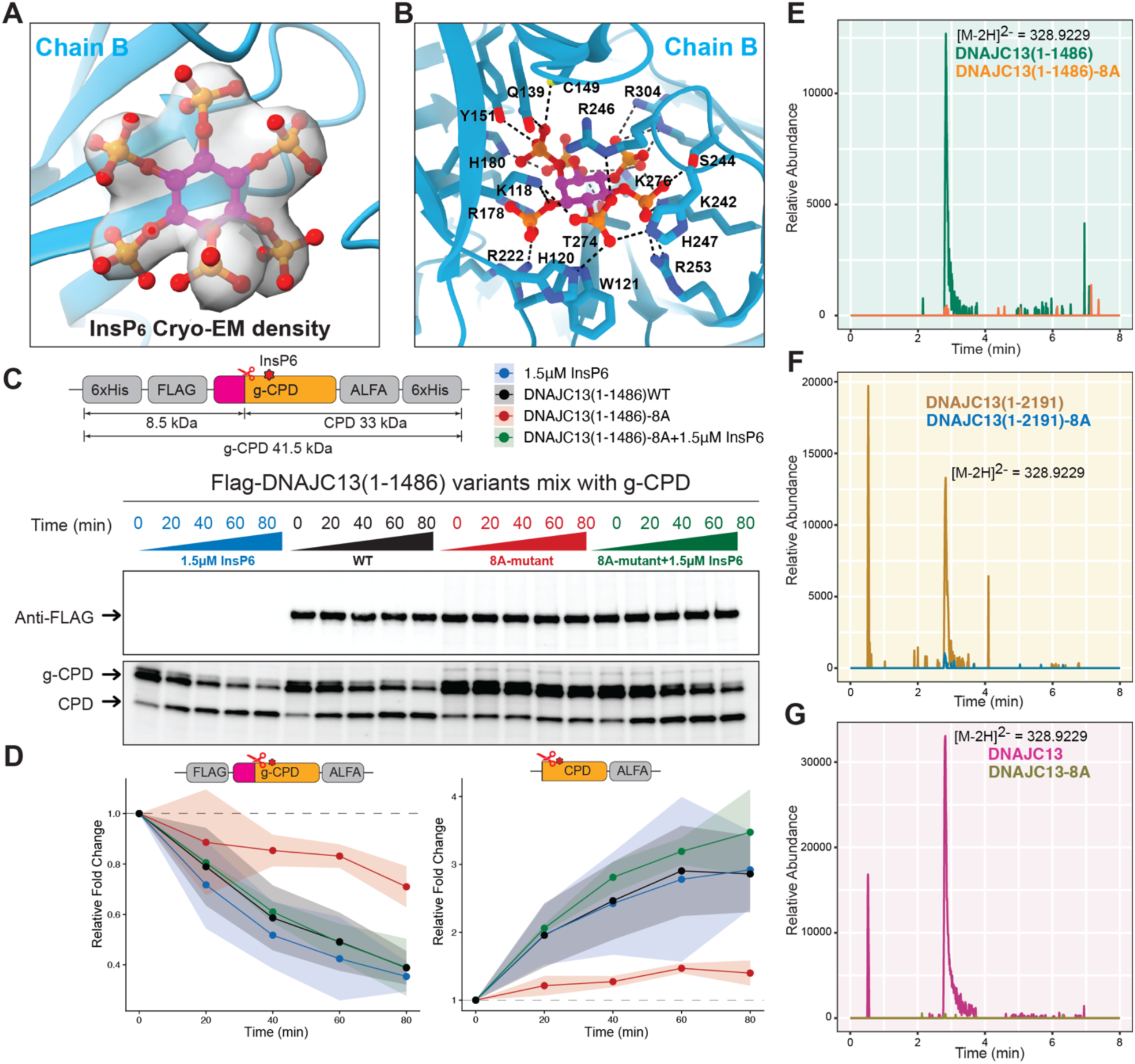
DNAJC13 binds InsP6 via tandem PHL2-PHL3 domains. (A) Cryo-EM density corresponding to InsP_6_ within the pocket formed by PHL2 and PHL3 in Chain B. (B) Binding interface of InsP_6_ by basic and polar residues from PHL2 and PHL3. Side chains contacting InsP_6_ are shown as sticks, and hydrogen-bonding interactions are indicated by dashed lines. (C) Schematic and implementation of an autocatalytic g-CPD cleavage assay reporting on InsP_6_ released from denatured DNAJC13. Samples from heat-denatured Flag-tagged DNAJC13(1-1486) variants were mixed with g-CPD and incubated for the indicated times. Top, anti-FLAG immunoblot showing equal loading of InsP_6_ relative to the input DNAJC13 variant proteins. Bottom, anti-FLAG immunoblot detecting uncleaved g-CPD and cleaved CPD in reactions containing InsP_6_ (1.5 µM), WT DNAJC13, the InsP_6_-binding-deficient 8A mutant, or the 8A mutant supplement with InsP_6_. (D) Quantification of g-CPD decrease (left) and CPD accumulation (right) over time from three technical replicates of the assay in (C). Data are normalized to time zero and plotted as mean ± SD. (E-G) LC-MS analysis of InsP_6_ associated with DNAJC13 variants. Panel E: Extracted ion chromatograms (EICs) for InsP_6_ from DNAJC13(1-1486) WT (green) and 8A mutant (orange). Panel F: EICs for InsP_6_ from DNAJC13(1-2191) WT (gold) and 8A mutant (blue). Panel G: EICs for InsP_6_ from purified full-length DNAJC13 (magenta) and DNAJC13-8A (brown). Peaks corresponding to InsP_6_ (eluting at ∼3 min) are readily detected in WT samples but are absent in 8A mutants, indicating loss of InsP_6_ binding in the mutant.

We next sought to directly examine InsP_6_ binding to DNAJC13 using biochemical analysis and mass spectrometry in the context of WT and mutant PHL2/3 proteins. To disrupt InsP_6_ binding with minimal structural perturbation while also considering the highly positive charged pocket, we generated an alanine-substitution mutant (DNAJC13-8A mutant) in which eight residues implicated in InsP_6_ coordination were replaced by Ala (H120A, W121A, R178A, H180A, R222A, R246A, H247A, and R304A) (**Figures 2B and S3A**). To qualitatively assess endogenous InsP_6_ bound to DNAJC13, we first devised an autocatalytic cleavage assay using the *Clostridium difficile* Toxin A glucosyltransferase-cysteine protease domain (g-CPD) as an InsP_6_-dependent molecular biosensor (**Figure 2C and 2D**). InsP_6_ binding activates g-CPD, triggering its autocatalytic cleavage.^40^ Using this assay, we established a biochemical readout to compare InsP_6_ levels released from purified DNAJC13(1-1486) WT and DNAJC13(1-1486)-8A mutant proteins by thermal denaturation (see **STAR METHODS**). By immunoblotting, we observed that samples released from DNAJC13(1-1486) WT caused a time-dependent decrease in g-CPD protein abundance and a corresponding increase in the abundance of cleaved CPD, similar to the positive control in which 1.5 µM of InsP_6_ standard was added, indicative of InsP_6_-dependent auto-cleavage (**Figure 2C**). In contrast, with the DNAJC13(1-1486)-8A mutant samples, the abundance of g-CPD and CPD remained essentially unchanged over 1 h, but cleavage was activated upon addition of 1.5 µM InsP_6_ to the 8A mutant sample as an additional positive control (**Figure 2C**). Quantification of protein intensities across three replicate assays confirmed the presence of InsP_6_ in WT but not 8A DNAJC13 (**Figure 2D and Data S2.4**). Next, to directly and quantitatively demonstrate InsP_6_ in association with DNAJC13, we analyzed three different truncations of DNAJC13-WT and -8A mutant proteins purified from Expi293F cells by mass spectrometry. For the WT protein samples, we detected a 328.9229 m/z species matching pure InsP_6_ as a molecular standard ([M-2H]^2-^) (**Figure 2E-G, Data S2.5**). In contrast, InsP_6_ was not detected in the DNAJC13-8A mutants, indicating that InsP_6_ binding was abolished by these substitutions (**Figure 2E-G**). These data confirm DNAJC13 as an InsP_6_-binding protein and provide a molecular tool for examining its potential functional role.

### InsP_6_ Binding Supports DNAJC13 Association with Membranes

PHL2 and PHL3 form a composite InsP_6_-binding domain that is sandwiched between PHL1 and the first ARM domain respectively (**Figure 1C and 2B**). We examined whether InsP_6_ binding might contribute to PHL1’s interaction with PI(3)P within membranes. We first established an imaging-based in vitro Giant Unilamellar Vesicle (GUV)-recruitment assay and tested PI(3)P dependence using 80 nM DNAJC13(1-1486)-mSG2(monomericStayGold2)-Flag or Flag-DNAJC13(1-1486)-acGFP, approximating the endogenous DNAJC13 concentration in HEK293T cells (81 nM).^41^ Both proteins displayed a time-dependent accumulation on GUVs containing 5% PI(3)P but failed to associate with GUVs lacking PI(3)P (**Figures S3B-S3E**). When WT and 8A mutant Flag-DNAJC13-mSG2(1-1486) proteins (**Figure S3D**) were assayed in parallel at 80 nM, WT protein displayed a time-dependent accumulation on PI3P-containing GUVs while the 8A mutant remained largely diffuse (**Figures 3A**). Quantification of membrane-associated mSG2 fluorescence confirmed these observations: signal at the GUV rim displayed a time dependent increased for WT DNAJC13 but was substantially reduced with the 8A mutant (**Figure 3B**). These data demonstrate that an intact InsP_6_-stabilized triple PHL module promotes PI(3)P binding in vitro.

**Figure 3.**
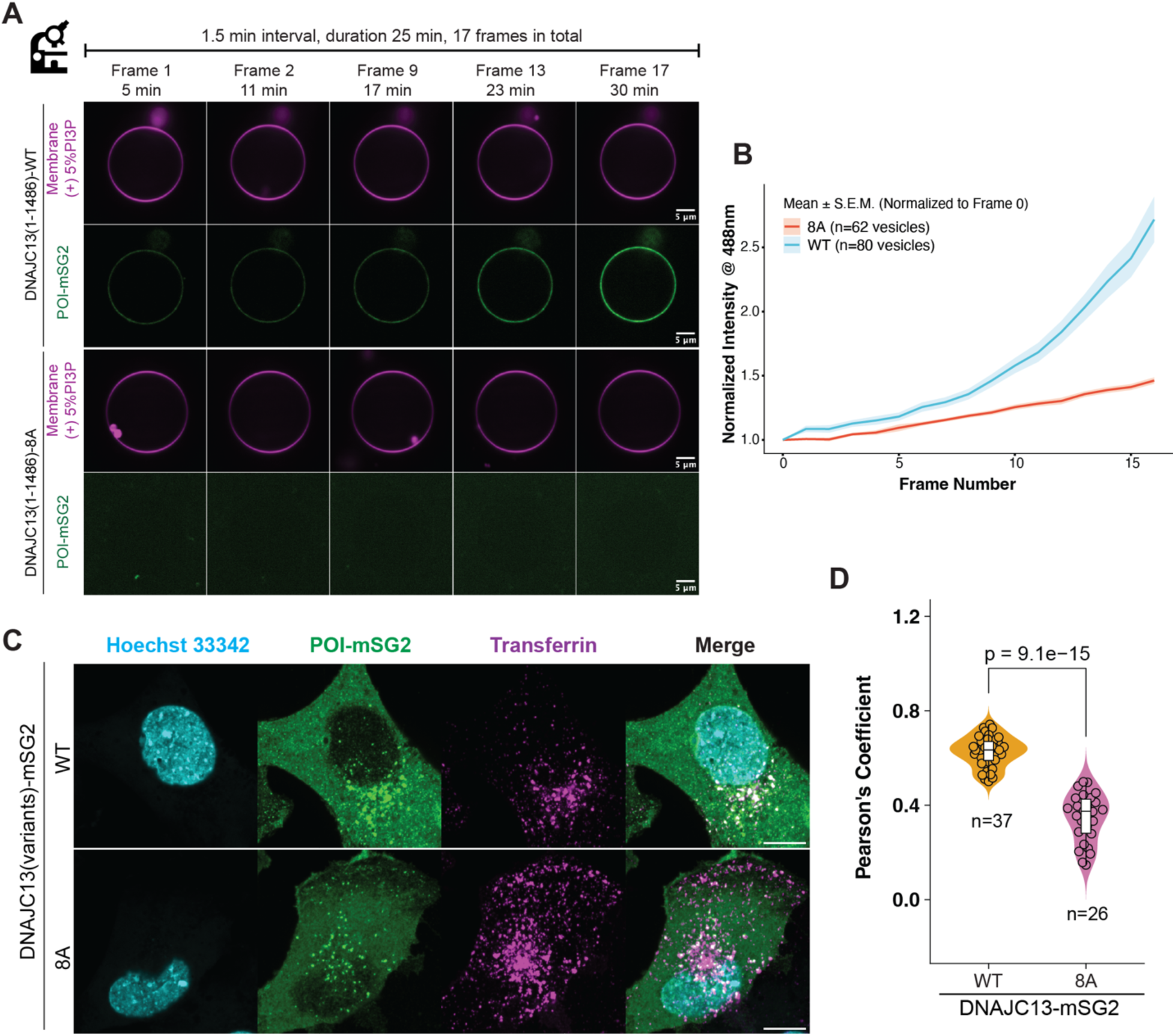
InsP_6_ binding promotes recruitment of DNAJC13 to PI(3)P-containing membranes. (A) Time-lapse confocal imaging of Flag-DNAJC13(1-1486)-mSG2 WT and 8A proteins incubated with GUVs containing 5% PI3P. Magenta, fluorescent lipid membrane dye; green, DNAJC13-mSG2. Representative frames from the 25-min imaging series (1.5-min interval, 17 frames total) are shown. WT DNAJC13 accumulates progressively on the GUV membrane, whereas the InsP_6_-binding-deficient 8A mutant fails to bind detectably. Scale bars, 5 µm. (B) Quantification of DNAJC13-mSG2 fluorescence at the GUV rim over time for WT and 8A variant. Curves represent mean ± S.E.M. of normalized membrane intensity (488-nm channel) from the indicated number of vesicles. (C) Confocal images of SUM159 DNAJC13-KO cells reconstituted with WT or 8A DNAJC13-mSG2 (green, see **Data S2.3**) 15 min post treatment with fluorescent Transferrin (magenta). Nuclei are labeled with Hoechst 33342 (cyan). Scale bar, 20 µm. (D) Pearson’s correlation analysis of DNAJC13-Transferrin colocalization for WT and 8A DNAJC13 15 min post-Transferrin treatment. Violin plots show the distribution of Pearson’s coefficients, with median and interquartile ranges indicated; n denotes the number of cells analyzed and p values (two-tailed t-test) are shown above the corresponding comparisons.

Previous studies have demonstrated that DNAJC13 concentrates on Transferrin-positive endosomal structures in response to Transferrin endocytosis.^42^ We employed this assay to assess the effect of PHL2 and PHL3-domain mutations in cells. As expected, DNAJC13 displayed extensive co-incident localization with fluorescent Transferrin-positive endosomes in DNAJC13-KO SUM159 cells reconstituted with WT DNAJC13-mSG2, as assessed by confocal microscopy and co-incidence detection (**Figures 3C and 3D, Data S3**). In contrast, the 8A mutant displayed reduced co-localization with Transferrin-positive endosomes, indicating impaired endosomal targeting, although the mutant protein also displayed reduced levels (**Figures 3C and 3D, Data S3**). Interestingly, structural analysis revealed that E264 does not contact InsP_6_ directly but instead forms hydrogen bonds with the side chains of K276 and K242 in PHL3, which directly engage InsP_6_ phosphates (**Figure S3F**). As such, we hypothesized that E264 acts to stabilize the local geometry of the InsP_6_-binding pocket. Consistent with this hypothesis, WT DNAJC13-mSG2 displayed robust punctate localization and strong overlap with Transferrin after 5-25 min of uptake, while the E264A mutant remained more diffuse and displayed reduced co-localization with Transferrin-positive endosomes, despite being expressed at near WT rescue levels (**Figure S3G and Data S3**). Quantification of Pearson’s coefficients for DNAJC13-Transferrin co-localization confirmed a significant decrease for E264A at all time points examined (**Figure S3H**). Introduction of a positively charged lysine at this position (E264K), expected to further perturb the pocket via electrostatic repulsion, resulted in further loss on co-localization with Transferrin (**Data S2.3**). Together with in vitro GUV assays, this data implicates InsP_6_ binding via PHL2/3 in the ability of DNAJC13 to associate with membranes containing PI(3)P.

### DNAJC13 dimerization via two discrete interfaces

DNAJC13(1-1486) forms an arch-shaped dimeric structure where the two protomers interact at the top of the arch (**Figures 1C**). Inspection of the model from the perspective of the C2 symmetry axis shows that dimerization is mediated by two interfaces (Site 1 and Site 2) near the ARM2 α-solenoid, with Site 1 being present in two copies based on symmetry (**Figures 4A and 4B**). At the Site 1 interface, a β-sheet and loop segment between IWN1 and ARM2 of chain A docks against a corresponding β-sheet and loop insertion in ARM2 from chain B to form an extended polar interface (**Figure 4C**). A dense network of largely conserved salt bridges and hydrogen bonds is observed between residues on chain A (N805, V806, S808, N810, E813, E815, K817) and residues on chain B (R1061, D1062, R1069, R1076, K1073, D932), together with several backbone-backbone contacts (**Figures 4C and 4D, S2H,I**). This highly complementary electrostatic surface suggests that Site 1 provides the principal driving force for dimer formation, and is present in the symmetrical protomer B to protomer A interface (**Figure 4A**). In contrast, Site 2 lies near the apex of the arch, also within the ARM2 region, and is dominated by hydrophobic packing with a more limited set of polar interactions (**Figures 4E and 4F, S2G-I**). Here, helices from chain A (Y858, H859, L862, L863, R860, M900) interdigitate with helices from chain B (Y858, H859, L862, L863, D893, Y896, M900), creating a small but tightly packed interface. Aromatic and aliphatic side chains form a contiguous hydrophobic patch (L862, L863, Y858, Y896, M900), while a few ionic interactions, such as R860:Y896 and H859:D893, appear to provide additional specificity (**Figures 4E and 4F**). Together, these structural observations indicate that DNAJC13 forms a stable homodimer involving two discrete conserved interfaces (**Figure S2I**).

**Figure 4.**
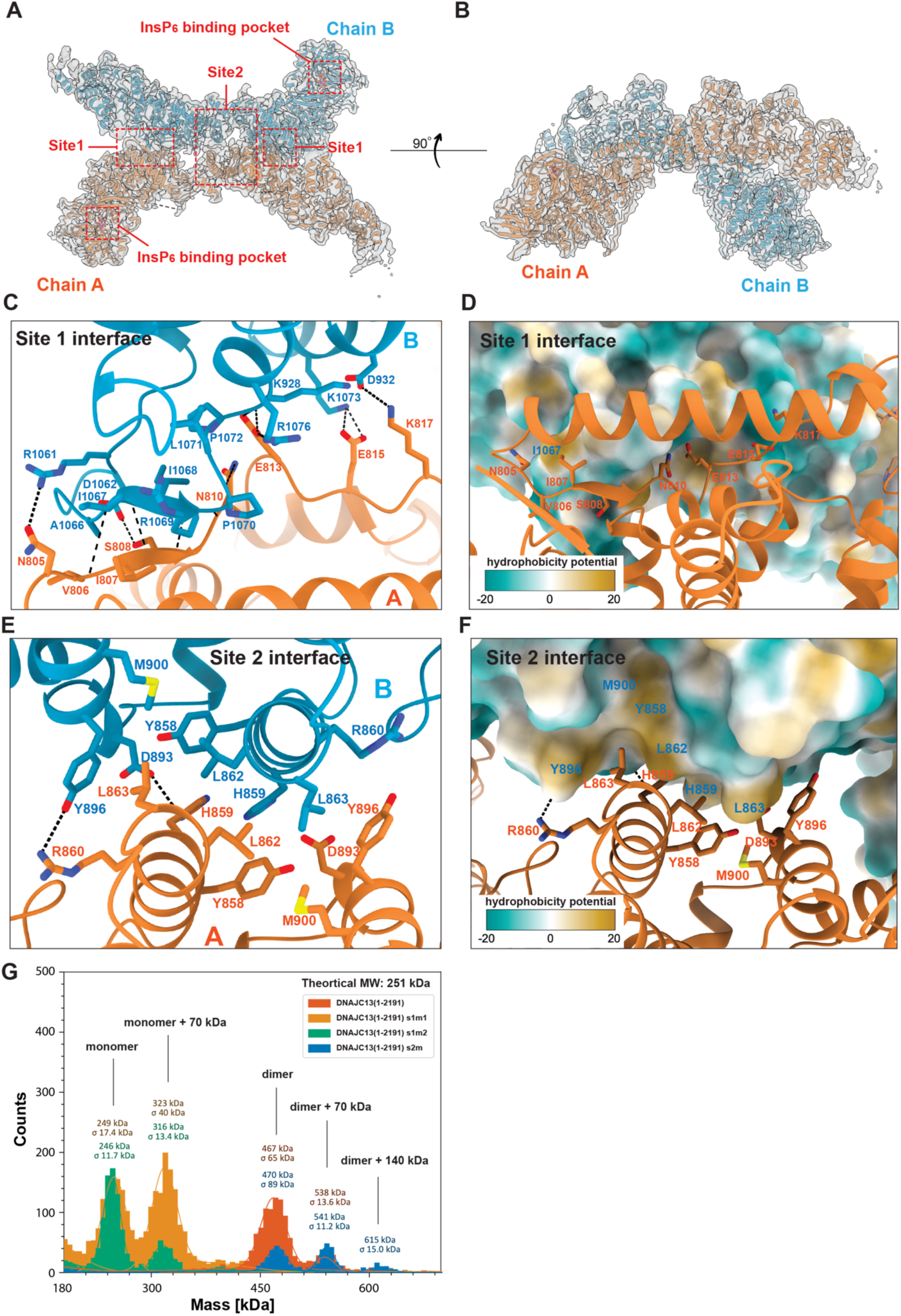
DNAJC13 homodimerization via two discrete interfaces. (A-B) Superposition of the cryo-EM density and atomic model of the DNAJC13 dimer viewed along its long axis (Panel A) and rotated 90 °C (Panel B). The two protomers are colored orange (chain A) and blue (chain B), with the cryo-EM map shown as a semi-transparent gray surface. Red dashed boxes highlight the PHL2-PHL3 InsP_6_-binding pocket and the two interfaces (Site 1 and Site 2) that mediate dimerization. (C-D) Site-1 interface. (Panel C) Close-up of Site-1 showing chain A (orange) and chain B (blue) residues involved in intermolecular contacts. Key salt bridges and hydrogen bonds are indicated by black dashed lines; selected side chains are labeled. (Panel D) Surface representation of chain B colored by hydrophobicity potential (scale shown), with the interacting helix from chain A shown as an orange ribbon. (E-F) Site-2 interface. (Panel E) Close-up of Site-2 showing chain A (orange) and chain B (blue) residues involved in intermolecular contacts. Key hydrogen bonds are indicated by black dashed lines; selected side chains are labeled. (Panel F) Surface representation of chain B colored by hydrophobicity potential, with the contacting segment of chain A in orange ribbon, illustrating the predominantly hydrophobic pocket formed at Site-2. (G) Mass photometry of DNAJC13(1-2191) and dimer-interface mutants. Site-1 mutants (s1m1 and s1m2) yield peaks corresponding to monomer and monomer bound to a 70-kDa complex. Wild-type DNAJC13(1-2191) and the Site-2 mutant (s2m) show predominant peaks corresponding to dimer and dimer bound to one or two 70-kDa complexes.

To test the contributions of these interfaces, we designed three mutants: two targeting Site 1 — s1m1 (V806G, S808G, N810G, E813A, E815A) and s1m2 (V806R, S808R, N810G, E813A, E815A) — and one targeting Site 2 — s2m (D893A, Y896A). Mass photometry of DNAJC13(1-2191) and DNAJC13(1-1486) purified from Expi293F or insect cells showed a predominant species corresponding to the homodimer, together with minor peaks assignable to dimer plus ∼70 kDa (**Figure 4G; Figures S4A and and S4B**), confirming that WT DNAJC13 dimerizes in solution with a mass consistent with the structural model. In contrast, mass photometry of the Site 1 mutants (s1m1 and s1m2) in either DNAJC13(1-2191) or (1-1486) revealed only monomer and monomer plus ∼70 kDa peaks, indicating that neutralizing or reversing charges at Site 1 is sufficient to disrupt dimerization (**Figures 4G and S4A**). By comparison, the Site 2 s2m mutation displayed peaks matching the dimer mass as well as the dimer plus ∼70 kDa (**Figure 4G**), suggesting that symmetrical Site 1 interactions are sufficient to support dimer formation in this context. We further assessed dimerization in cells using ALFA magnetic-bead co-immunoprecipitation of ALFA-tagged DNAJC13-mSG2 variants co-expressed with FLAG-tagged DNAJC13 variants or FLAG-GFP as a negative control in HEK293T cells (**Figure S4C and Data S2.2**). Robust co-precipitation was observed for WT and the Site 2 mutant, whereas both Site 1 mutants failed to co-precipitate with their FLAG-tagged partners (**Figure S4C**). Transferrin uptake assays showed that dimerization-defective mutants (s1m1 and s1m2) form puncta and colocalize to Transferrin-labeled endosomes at levels comparable to WT (**Data S2.3**), consistent with the expectation that membrane recruitment is mediated by the N-terminal PHL domains. Together, these biophysical and co-immunoprecipitation experiments demonstrate that DNAJC13 dimerization is primarily driven by an extensive, electrostatically complementary interface at Site 1, with a second, more hydrophobic interface at Site 2 that locks the two ARM2 solenoids at the apex of the arch (**Figures 4A and 4B**).

### Proteomic landscape of endosomes using DNAJC13 as an Endo-IP handle

Having identified mutants that result in disruption of DNAJC13 dimerization without impairing its recruitment to endosome membranes (**Data S2.3**), we next sought to understand the role of dimerization in its ability to coordinate protein interactions at endosomes. We envisioned that, as with the PI(3)P binding FYVE-domain of EEA1,^43,44^ the interaction of the PHL1 domain with PI(3)P could serve as a molecular handle for isolation of DNAJC13-associated endosomes using detergent-free extracts. To initially test the utility of DNAJC13 as an endosomal isolation handle (DNAJC13 Endo-IP), we stably expressed ALFA-mSG2-EEA1 and ALFA-DNAJC13-mSG2 in EEA1-KO or DNAJC13-KO HEK293T cells, respectively, at comparative levels (**Data S4** and **STAR METHODS**). This approach allowed us to quantitatively compare the proteomes of endosomal populations purified in biological quadruplicate with a single ALFA-tag affinity resin in the same Tandem Mass Tagging pro (TMTpro) 16-plex experiment, using the corresponding deletion cells as negative controls for enrichment (**Figure 5A**).

**Figure 5.**
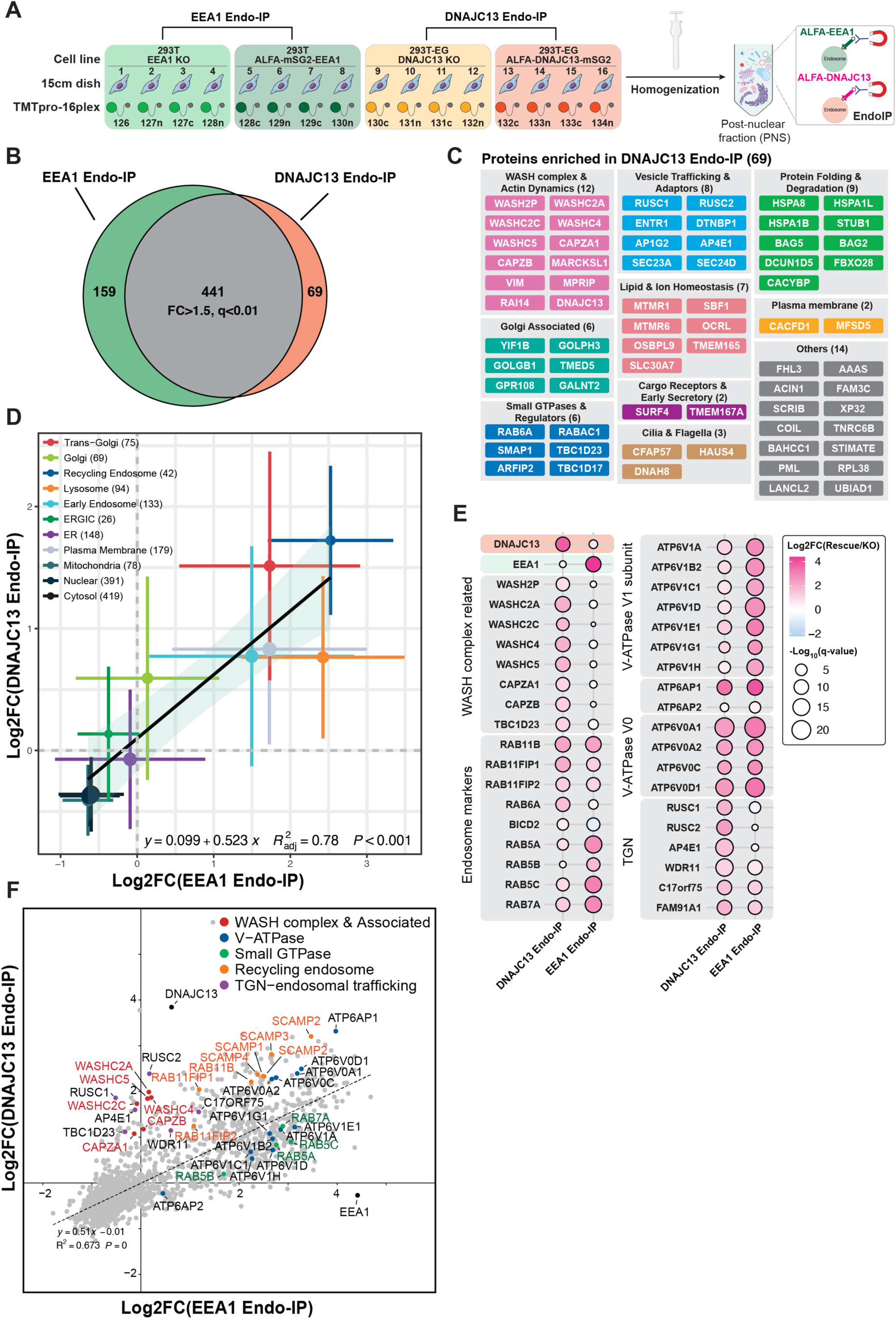
DNAJC13 Endo-IP enrich TGN and recycling endosome components. (A) Schematic of the endosomal immunoprecipitation (Endo-IP) workflow. For EEA1 Endo-IP, HEK293T EEA1-KO cells and HEK293T EEA1-KO cells rescued with ALFA-mSG2-EEA1 were processed in parallel. For DNAJC13 Endo-IP, HEK293T^EG^ DNAJC13-KO cells and HEK293T^EG^ cells rescued with ALFA-DNAJC13-mSG2 were used (see **STAR METHODS**). Cells from 15-cm dishes are harvested and homogenized to generate post-nuclear supernatants (PNS). ALFA-tagged EEA1- or DNAJC13-positive endosomes are then captured from the PNS using ALFA-nanobody magnetic beads for subsequent mass-spectrometry analysis of endosome-associated proteins. Four biological replicates were included for each condition with the indicated TMTpro reporters employed. (B) Venn diagram summarizing the overlapping and differential proteins enriched in EEA1 and DNAJC13-based Endo-IPs (log_2_ FC>1.5, q<0.01). The number of common and unique proteins are indicated. (C) Functional categorization of the 69 high-confidence DNAJC13-specific endosome proteins. Proteins are grouped into vesicle trafficking and fusion, vesicle coat and structural scaffolds, surface receptors/cargo/signaling, endosomal trafficking and sorting, ion channels/transporters and pH regulation, lipid transport and homeostasis, lysosomal proteins, Golgi/trans-Golgi associated factors, small GTPases and regulators, and others. Colors indicate major functional classes as indicated in the panel header. (D) Organelle correlation plot representing log_2_ FC enrichment for proteins from the indicated organelle designations, as performed previously.^45^ The number of proteins in each category are indicated in parenthesis. Dashed line indicates the linear regression fit. (E) Bubble heatmap comparing the relative enrichment of selected endosomal markers and complexes in DNAJC13 Endo-IP versus EEA1 Endo-IP. Circle color indicates log_2_ fold change (rescue/KO) and circle size represents -log_10_(q-value). Proteins include WASH complex subunits and partners, V-ATPase subunits, recycling-endosome markers (RAB11B, RAB11FIP1/2), TGN-endosomal trafficking factors (RUSC1/2, AP-4 subunits, C17orf75), and early/late-endosome RABs (RAB5, RAB7 family members). (F) Correlation scatter plot of log_2_ FC (rescue/KO) for proteins detected in EEA1 Endo-IP (x-axis) versus DNAJC13 Endo-IP (y-axis). Selected proteins are colored and annotated by functional class (WASH complex and associated proteins, V-ATPase, small GTPases, recycling-endosome markers, and TGN-endosomal trafficking factors). Dashed line indicates the linear regression fit.

Initial comparison of proteins enriched in ALFA-tag pulldowns relative to the corresponding deletion cells revealed that DNAJC13 could be used as an affinity handle for endosome enrichment. Indeed, DNAJC13 and EEA1 Endo-IPs share 441 proteins (FC > 1.5, q < 0.01), representing core endosomal machinery^43^ (**Figure 5B, Table S2**). This included multiple V-ATPase subunits, coat components, vesicle-fusion factors, RAB GTPases and their regulators, lumenal endo-lysosomal hydrolases and their receptors (e.g., M6PR, IGF2R), and cargo such as TFRC, SORT1, SORL1, and LDLR. Beyond this shared set of proteins, each bait displayed a characteristic enrichment profile (**Figures 5B and 5C, and S5A**; **Table S2**). DNAJC13 Endo-IPs were exclusively enriched for 69 proteins comprising all WASH complex subunits, HSP70 chaperones (HSPA1B, HSPA1L, HSPA8), the phosphatidylinositol phosphatases MTMR1, MTMR6, and OCRL, and components of the AP-4 Golgi-derived vesicle transport machinery (**Figure 5C**). In contrast, EEA1 Endo-IPs were selectively enriched for 159 proteins, including ESCRT components, a subset of luminal endo-lysosomal proteins, and subunits of the HOPS/CORVET tethering complexes, consistent with a broad distribution across the endo-lysosomal pathway (**Figure S5A**).

Correlation analysis of log_2_ FC for enriched proteins grouped by subcellular compartment^45^ revealed that EEA1-positive endosomes were comparatively enriched for more lysosomal components, whereas DNAJC13-positive endosomes displayed a bias towards recycling-endosome and trans-Golgi/Golgi proteins (**Figures 5D and S5B**). To further explore common and distinct molecular landscapes of DNAJC13 and EEA1-enriched organelles, we compared individual proteins quantified in the same TMTpro plex based on both the fold-enrichment within the immunoprecipitation (**Figure 5E**) and based on correlation plots, where deviation from the overall regression (y ≈ 0.51x) profile allows detection of proteins displaying biased enrichment (**Figure 5F**). First, DNAJC13 Endo-IP selectively enriched all WASH complex subunits and associated proteins (**Figures 5E and 5F**), indicating that WASH complex components are preferentially recruited to DNAJC13-containing endosomes. This result is consistent with the proposed direct interaction between the WASH subunit WASHC2A (also called FAM21A) and DNAJC13.^19^ WASH complex activity promotes Arp2/3-dependent branched actin assembly on sorting and recycling endosome domains.^46^ Second, compare to EEA1-positive endosomes, DNAJC13-positive endosomes were de-enriched for RAB7A (canonical late-endosome marker) and RAB5A/B/C (canonical early-endosome marker), consistent with immunoblotting of RAB5A (**Data S4**), while being similarly or more enriched for RAB6A, RAB11B, RAB11FIP1, and RAB11FIP2, which are involved in trafficking through the TGN or ERC (**Figures 5E and 5F**). Third, DNAJC13-positive endosomes were strongly enriched for AP-4, RUSC1, and RUSC2 (**Figure 5E**), factors implicated in vesicular trafficking from the TGN to the early endosome.^47^ Fourth, DNAJC13 and EEA1 Endo-IPs displayed comparable enrichment of V0-ATPase, but DNAJC13 Endo-IPs were less enriched in V1-ATPase subunits (**Figure 5E and 5F; Figure S5D**). Finally, the recycling-endosome carrier SCAMP1/2/3/4 proteins^48,49^ as well as the WDR11-TBC1D23-C17ORF75 Golgi-directed recycling complex^50^ were also similarly enriched in DNAJC13- and EEA1-positive endosomes (**Figure 5E and 5F**). Notably, these enrichment trends were consistent between two independent sets of EEA1 Endo-IP and DNAJC13 Endo-IP experiments in separate TMTpro multiplexes (**Data S4**). In addition, these enrichment patterns were reproducible across multiple DNAJC13 Endo-IP experiments performed in two independent DNAJC13-KO clonal lines, using either non-tagged rescue or KO cells as controls (**Figure S5C and S5D, Data S2.2**). Taken together, these data indicate that whereas EEA1 predominantly marks a broad endo-lysosomal population, DNAJC13 is more closely associated with a subpopulation of endosomes enriched in trans-Golgi network and recycling-endosome proteins, WASH complex components, and AP-4/RUSC-dependent trafficking machinery, and thus can be used as an affinity reagent to enrich a population of vesicles biased towards recycling endosomes. These data are consistent with prior studies demonstrating a requirement for DNAJC13/RME-8 in recycling of multiple cargoes, including CI-MPR ^42,51^, Wntless^17^, Notch receptor^52^, and GPCRs ^20^.

### DNAJC13 dimerization promotes WASH complex recruitment to endosomes

We next used the DNAJC13 Endo-IP approach to examine the contribution of DNAJC13 dimerization to endosomal recruitment. DNAJC13 Endo-IP proteomes were obtained using dimerization competent (WT and s2m) and dimerization-defective (s1m1 and s1m2) DNAJC13 proteins, with DNAJC13-KO cells as a negative control for enrichment (**Data S2.2 and S2.3**). Endo-IPs from dimeric DNAJC13 robustly enriched all WASH complex subunits (WASHC1/2A/2C/3/4/5), their actin-capping partners (CAPZA1/A2, CAPZB) and FKBP15, whereas these proteins were essentially absent with Endo-IPs from dimerization-defective DNAJC13 proteins, as assessed by quantitative proteomics (**Figure 6A and 6B, S6A-C**). Notably, monomeric forms of DNAJC13 maintained an ability to interact with Transferrin-positive endosomes based on confocal microscopy (**Data S2.3**) and captured the vast majority of endosomal proteins detected with the dimer, including strong retention of several cargo receptors (e.g. TFRC), Retromer subunits (VPS26/29/35) and V-ATPase subunits (**Figure 6A and 6B, S6A-C**). Together, this data suggests that DNAJC13 dimerization is particularly critical for recruitment of WASH machinery to endosomes but is not obligatory for association of DNAJC13 with endosomes via its PHL1 domain.

**Figure 6.**
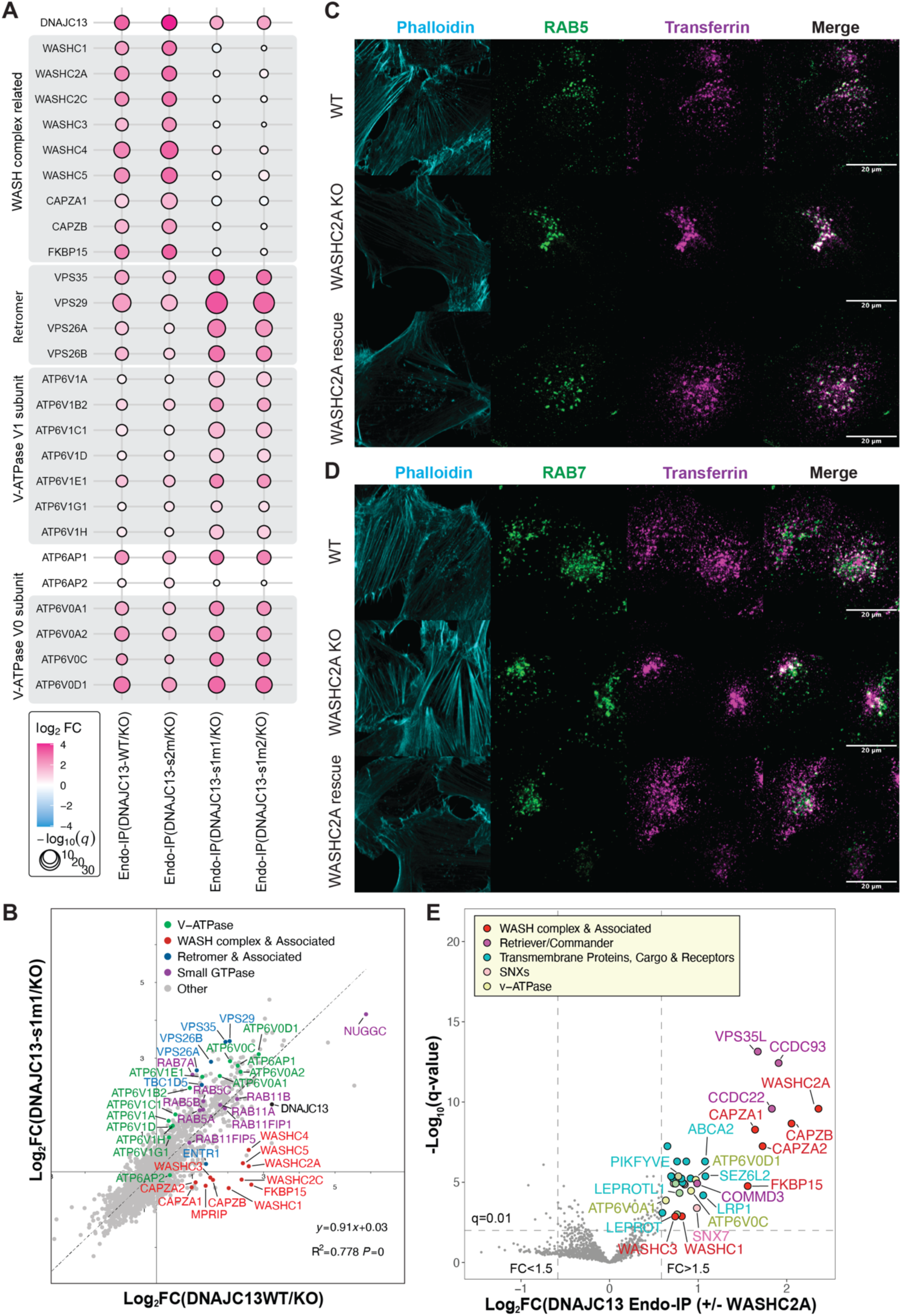
DNAJC13 dimerization is required for WASH complex assembly at recycling endosomes. (A) Bubble heatmap showing log_2_ FC (color scale) and statistical significance (circle size, -log_10_(q-value)) for WASH-complex subunits, Retromer components, and V-ATPase V0 and V1 subunits in Endo-IP proteomes from WT DNAJC13, the dimerization-competent mutant (s2m), and dimerization-defective mutants (s1m1 and s1m2), as indicated. (B) Scatter plot of protein enrichment in Endo-IP experiments comparing log_2_ FC (DNAJC13 WT/KO) with log_2_ FC (DNAJC13-s1m1/KO). WASH-complex and associated proteins (red), retromer and associated proteins (blue), V-ATPase subunits (green), and small GTPases (purple) are highlighted. Dashed line indicates the linear regression fit. (C-D) Confocal images of WT, WASHC2A^-/-^ (KO), and KO cells rescued with WASHC2A after 10-min uptake of fluorescent Transferrin (magenta), co-stained for RAB5 (Panel C, green) or RAB7 (Panel D, green), with F-actin visualized by phalloidin (cyan). Scale bars, 20 µm. (E) Volcano plot of DNAJC13 Endo-IP comparing WASHC2A-rescued versus WASHC2A-knockout (KO) cells. The x-axis shows log_2_ FC (+/−WASHC2A), and the y-axis shows −log_10_(q-value). Labeled proteins include WASH complex and associated proteins (red), Retriever/Commander subunits (purple), transmembrane cargo proteins and receptors (cyan), Sorting Nexins (pink), V-ATPase subunits (lime).

To define the contribution of the WASH complex to the DNAJC13 endosomal population, we performed DNAJC13 Endo-IP in WASHC2A-KO cells and in these cells rescued with WASHC2A (**Data S2.2**). Previous studies indicate that WASHC2A directly associates with DNAJC13.^19^ Volcano plot analysis revealed marked loss of WASH complex subunits, actin-capping partners (CAPZA1/A2, CAPZB), FKBP15, endocytic cargo (LRP1, SEZ6L2), and Retriever/Commander subunits from DNAJC13-positive endosomes upon WASHC2A deletion, underscoring this WASH complex subunit as a key organizer of the DNAJC13 compartment (**Figure 6E**). Consistently, WASHC2A deficiency had pronounced effects on cargo/early-endosome trafficking. First, WASHC2A deletion resulted in clustering of Transferrin-positive vesicles away from the cell cortex, as observed in fixed confocal images (**Figure 6C and 6D**) and as quantified using Ripley’s K function spatial analysis (**Figure S6D**). Second, after a 10-min Transferrin pulse, Transferrin accumulated in clustered RAB5- or EEA1-positive early endosomes with increased co-localization (**Figure 6C and S6E, Data S5**), but not in RAB7-positive late endosomes (**Figure 6D and Figure S6F**), indicating defective early-endosome sorting and recycling in WASHC2A mutant cells. Re-expression of WASHC2A restored the Transferrin and early-endosome distribution and Pearson’s coefficients for co-localization to near WT levels (**Figure 6C and S6E, Data S5**), demonstrating that WASH complex assembly via WASHC2A-DNAJC13 is necessary for proper endosomal sorting and recycling of Transferrin. Together with our proteomic data, these findings support a model in which dimeric DNAJC13 recruits and organizes the WASH complex likely in recycling-endosome/TGN-directed populations of the endosomal network.

### DNAJC13 Dimerization Controls Endosomal Tubular Fission

The dependence of DNAJC13 dimerization on WASH recruitment to endosomes led us to examine the link between endosomal tubule formation during recycling endosome formation in the context of Transferrin trafficking.^19,42^ In SUM159 WT cells, internalized Transferrin accumulated in numerous puncta distributed throughout the cytoplasm, with a subset in a perinuclear compartment (**Figure 7A**). In contrast, DNAJC13-KO cells displayed strikingly elongated, branched tubular endosomal structures in fixed cells, indicative of abnormal tubulation likely resulting from impaired endosome fission, as reported previously^19^ (**Figure 7A, Data S6**). Live-cell imaging after Transferrin treatment confirmed the presence of highly dynamic tubules in DNAJC13-KO cells (**Figure 7B and S7; Movie S2**). Expression of dimeric DNAJC13-mSG2 (WT or s2m) largely normalized endosomal morphology, whereas cells expressing monomeric DNAJC13-mSG2 (s1m1 and s1m2) retained prominent DNAJC13- and Transferrin-positive tubules that emanated from endosomes (**Figure 7B and S7A and S7B**). In order to quantify the extent of tubulation and vesicle deformation, we employed Feret analysis in DNAJC13-KO SUM159 cells expressing either WT or the s1m2 mutant. At 4 min post Transferrin addition, tubule Feret values increased from a mean of 1.8 μm in WT rescued cells to 2.45 μm in monomer rescued cells, representing an increase in tubulation when DNAJC13 cannot dimerize (**Figure S7C**). Similarly, puncta Feret values increased from 0.98 μm in the context of WT DNAJC13 to 1.1 μm in the context of the monomer mutant (**Figure S7D**). Together, these data suggest that dimerization defects do not impair DNAJC13 recruitment to endosomes; however, DNAJC13 dimerization is required for efficient endosomal tubule fission and cargo sorting/recycling.

**Figure 7.**
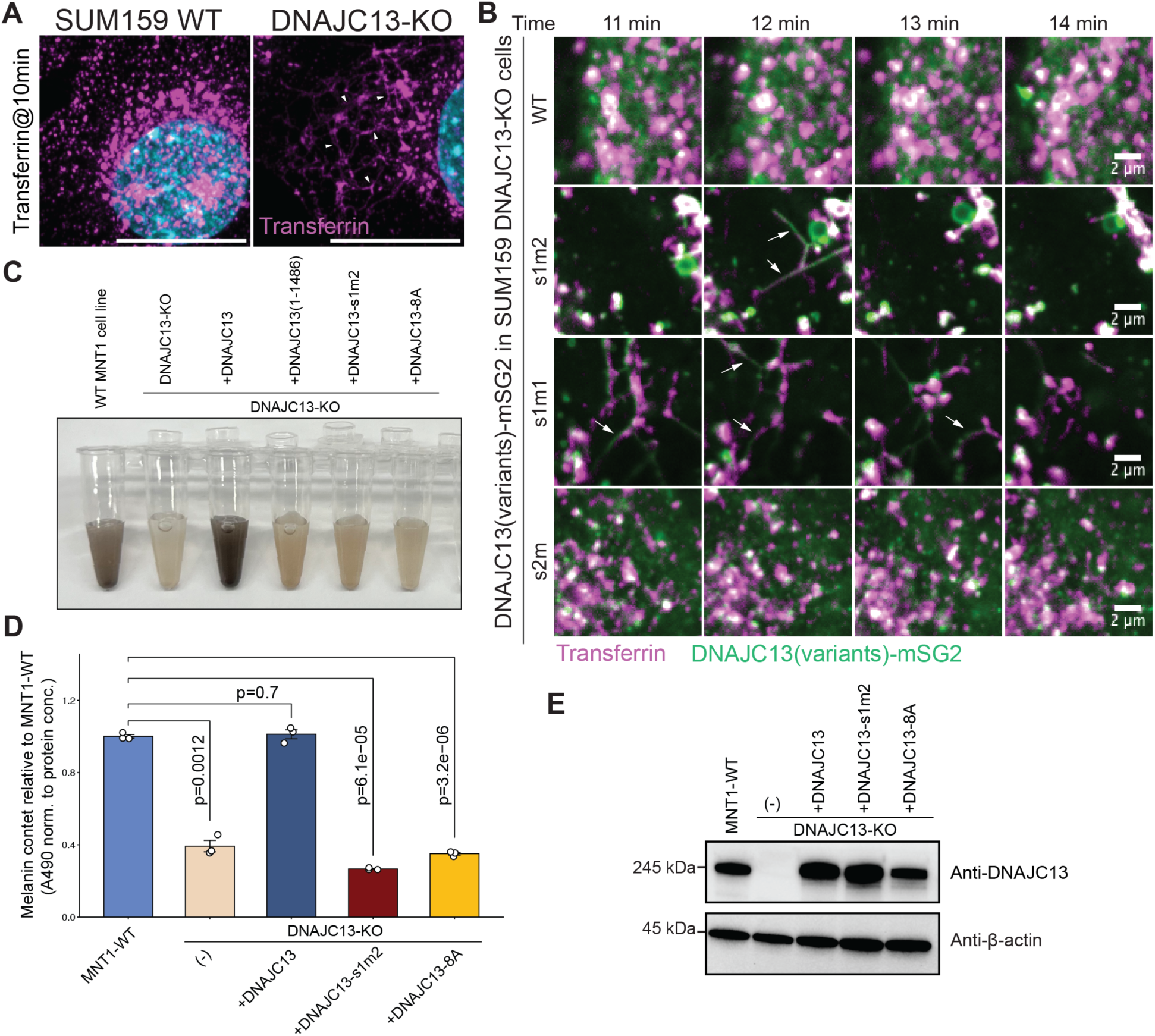
DNAJC13 dimerization controls endosomal tubule fission and melanosome maturation. (A) Confocal images of SUM159 WT and DNAJC13-KO cells after a 10-min uptake of fluorescent transferrin (magenta). Nuclei are stained with Hoechst 33342 (cyan). DNAJC13-KO cells display elongated, branched Transferrin-positive tubules (indicated by white arrowheads) compared with punctate endosomes in WT cells. Scale bars, 20 µm. A wider field view of these cells as well as the identical zoom version is including in **Data S6**. (B) Time-lapse imaging of DNAJC13-KO SUM159 cells expressing DNAJC13-mSG2 variants (green) after fluorescent Transferrin uptake (magenta). Representative zoomed-in regions and time points are shown. Dimeric DNAJC13 (WT and s2m) supports predominantly punctate Transferrin-positive endosomes, whereas monomeric variants (s1m1 and s1m2) show dynamic DNAJC13- and Transferrin-decorated tubules that emanate from endosomes and fail to undergo efficient fission (white arrows). Time after Transferrin addition are indicated. Scale bars, 2 µm. Additional views from the same experiment were shown in **Figure S7A**. (C) Representative cell suspensions in PBS from MNT-1-WT, DNAJC13-KO, and DNAJC13-KO cells rescued with DNAJC13-WT, DNAJC13(1-1486), DNAJC13-s1m2 (dimerization-defective mutant), or DNAJC13-8A (InsP_6_-binding-defective mutant), illustrating reduced pigmentation in DNAJC13-KO, DNAJC13(1-1486), s1m2 and 8A mutants. This experiment was performed independently from (Panel D) and (Panel E). (D) Quantification of melanin content in the indicated MNT-1 cell lines, normalized to total protein and expressed relative to MNT-1 WT. Bars show mean ± SD; individual data points are overlaid. p values (two-tailed t-test) are indicated above the corresponding comparisons. (E) Immunoblot analysis of DNAJC13 expression level in the MNT-1 cell lines used in (D). Anti-DNAJC13 and anti-β-actin (loading control) are shown.

### DNAJC13 dimerization and PHL2/3-domain integrity promotes melanosome maturation

A recent genetic screen revealed the involvement of endosomal trafficking and recycling proteins in melanosome maturation, including Retromer, Retriever, and WASH components.^30^ We therefore employed this system to further explore DNAJC13 structure-function relationships. We first validated that deletion of DNAJC13 with CRISPR-Cas9 in the human melanoma cell line MNT-1 (**Data S2.2**) results in loss of pigmentation, as assessed both by visual inspection of cells as well as by quantitative analysis of melanin (**Figure 7C-E**). Melanin production was fully rescued by re-expression of WT DNAJC13 at near endogenous levels. In contrast, expression of either a dimerization-defective mutant (DNAJC13-s1m2) or an InsP_6_-binding-defective mutant (DNAJC13-8A) at comparable levels failed to rescue pigmentation (**Figure 7C-E**). Taken together, our analysis indicates a critical role for DNAJC13 dimerization and InsP_6_-binding in multiple trafficking events linked with WASH and Retromer function.

## DISCUSSION

Our results reveal three major features of DNAJC13 that likely contribute to its endosomal regulatory functions. First, DNAJC13 forms an antiparallel homodimer that organizes multiple functional domains into a symmetric arch-like structure for the dimer core. This architecture imposes a defined spatial relationship between its N-terminal PHL1 domains, the J-domains, and the embedded GYF-like domains. The N-terminal PI(3)P-binding PHL1 domains are positioned in solvent-accessible conformations compatible with simultaneous interaction with a membrane surface. The J- and GYF-like domains are located adjacent to each other near the top of the arch; the hydrophobic binding groove in GYF, known in other GYF-containing proteins to interact with specific unstructured hydrophobic motifs,^35^ is solvent accessible. The proximity of the two sets of GYF and J-domains, related by C2-symmetry, raises the possibility that these domains function in tandem and/or across the dimer interface to facilitate interactions with other proteins. However, we note that the conformation of the J-domain in the composite structure is incompatible with formation of a functional interaction with HSP70, as HSP70 would clash with the other DNAJC13 protomer. Further studies are required to understand if the J-domain can adopt alternative conformations that would allow association with HSP70 proteins in the context of DNAJC13 dimers.

Second, our structural analysis of DNAJC13 reveals that InsP_6_ functions as a structural cofactor that likely stabilizes the tandem PHL2/3 domains into a compact tri-PHL module with PHL1. In vitro, disruption of InsP_6_ binding weakens DNAJC13 association with PI(3)P-containing GUVs when compared with WT DNAJC13 at equivalent protein concentrations. We also observed a reduction in association of DNAJC13 with Transferrin-positive endosomes in cells when DNAJC13 cannot bind InsP6. These data establish a mechanistic link between a soluble inositol phosphate and membrane targeting of an endosomal scaffold. While a single PH domain in Bruton’s tyrosine kinase has been shown to bind one face of InsP_6_,^53^ the use of a buried InsP_6_ to stabilize tandem PHL domains into a functional membrane-binding module appears to be unique.

Third, using proteomics, we demonstrated that DNAJC13 can function as an affinity handle for isolation of endosomes that are enriched in factors associated with recycling endosome trafficking. Compared with EEA1-positive endosomes, DNAJC13-positive organelles are depleted in canonical early and late endosomal RABs and in lysosomal components, but are instead enriched for WASH subunits, HSP70 chaperones, AP-4/RUSC machinery, and multiple recycling-associated factors, though such organelles likely reflect a continuum of states. Within this framework, Retromer and DNAJC13 itself appear to be retained on DNAJC13-positive membranes even when DNAJC13 dimerization is disrupted, whereas WASH recruitment is strongly dimer-dependent. Thus, DNAJC13 dimerization selectively supports WASH assembly without being essential for broader endosomal association or Retromer engagement, placing DNAJC13 functionally upstream of WASH in organizing actin-rich recycling domains. The requirement for DNAJC13 dimerization in WASH complex recruitment is mirrored by its role in controlling endosomal tubulation. Loss of DNAJC13 results in persistent tubules decorated with Transferrin, consistent with reduced rates of tubule fission. Dimerization-defective mutants rescue DNAJC13 localization to endosomes but not tubule hyper-elongation, indicating that simple recruitment of DNAJC13 to endosomal membranes is not sufficient to support its function. Instead, DNAJC13’s dimeric architecture is required to couple WASH-dependent actin assembly and/or fission to productive tubule scission and cargo recycling. Consistent with this model, we find that accumulation of melanin during melanocyte maturation requires both InsP_6_ binding and dimerization by DNAJC13, further highlighting that the dimerization mutant provides a powerful tool to mechanistically separate DNAJC13’s membrane targeting from its roles in endosomal tubulation, fission, and cargo recycling.

Although density within our structure ends within the third ARM domain, we employed Alphafold to generate a composite model for how the C-terminal 4^th^ and 5^th^ ARM domains may be oriented relative to the dimerization core. The overall organization (**Figure 1G**) suggests that engagement of PI(3)P on the membrane by PHL1 domains at the base of the arch would then place extended ARM domains across the dimer interface for each protomer. The functions of these likely flexible regions are unclear but the size of the arms in the extended conformation would allow DNAJC13 to potentially organize proteins across a membrane surface.

### Limitations of this study

While this work provides new insight into DNAJC13 structural mechanisms, several limitations remain. These include the absence of experimentally determined structures for the C-terminal ARM-IWN domains and the extreme C-terminus of DNAJC13, which is thought to be unstructured. This apparently reflects their flexibility relative to the defined core dimer structure. The predicted AlphaFold structures closely resemble the experimentally determined ARM-IWN structures, but the extent to which these regions interact with other proteins is unclear. Further studies are also necessary to understand the molecular surfaces on DNAJC13 that engage other endosomal machinery, including its interactions with sorting nexins, Retromer, and WASH components. Additionally, precisely how HSP70 proteins may engage DNAJC13 is unclear. Preparations of DNAJC13 from both human and insect cells contained HSP70 proteins. However, the extent to which these interactions reflect functional interaction with the J-domain or alternatively, a role for HSP70 in folding DNAJC13 itself, remains unknown. It is possible that HSP70 proteins are associated with the C-terminal regions of DNAJC13 that remain invisible in our cryo-EM maps. Thus, additional studies are necessary to understand how the DNAJC13 dimer engages multiple interacting partners to facilitate trafficking at recycling endosomes. Finally, while we demonstrate functional binding of InsP_6_ with DNAJC13, the extent to which InsP_6_ acts purely as a structural cofactor versus a regulated signal remains an open question. Our structural and biochemical studies provide a framework further elucidation of mechanisms underlying DNAJC13 function.

## RESOURCE AVAILABILITY

### Lead contact

Further information and requests for resources and reagents should be directed and will be fulfilled by the lead contact Wade Harper.

### Materials availability

Plasmids generated in this study are available from Addgene. Cell lines will be provided upon request.

### Data and code availability

Coordinates for the DNAJC13(1-1486)/InsP_6_ have been deposited at the Protein DataBank (PDB) under accession code 37FE, and the corresponding maps have been deposited in the Electron Microscopy DataBank (EMDB) under accession codes EMD-78135 (composite map), EMD-78131 (consensus map), EMD-78123, EMD-78128, EMD-78129, EMD-78130 (4 focused maps). All datasets will be made publicly available as of the date of the publication.

Integrated structure of full-length DNAJC13/InsP6 has been deposited in PDB-IHM with accession number 5-FDMP.

AlphaFold3 predictions of DNAJC13(1-1486) homodimer, DNAJC13 homodimer, DNAJC13/PI3P and DNAJC13/PI(3,5)P_2_ have been deposited in ModelArchive with accession number ma-z9nt8, ma-sgzb0, ma-germ0, and ma-vp0iy, as outlined in the Key Resources Table. All datasets will be made publicly available as of the date of publication.

All the MS proteomics data have been deposited to the ProteomeXchange Consortium through the PRIDE repository (http://www.proteomexchange.org/; project accession: PXD080914 and PXD080921). We used canonical protein entries from the human reference proteome database in our study (UniProt Swiss-Prot release 2024-01; https://ftp.uniprot.org/pub/databases/uniprot/previous_major_releases/). Code for proteomics data analysis and relevant figure generation is available on Github at https://github.com/taofu876/DNAJC13.git and on Zenodo at https://doi.org/10.5281/zenodo.21579490. Source data are provided with this paper.

Confocal microscopy raw data, quantification results, and R scripts used to generate the figures in this study are available at Zenodo under the following DOIs: 10.5281/zenodo.21579320, 10.5281/zenodo.21303971, 10.5281/zenodo.21312877, 10.5281/zenodo.21314099, and 10.5281/zenodo.21314844.

The data, code, protocols and key laboratory materials used and generated in this study are listed in a Key Resource Table The data, code, protocols and key laboratory materials used and generated in this study are listed in a Key Resource Table deposited on Zenodo (DOI: 10.5281/zenodo.21613970).

Any additional information and all the relevant raw data required to reanalyze the data reported in this paper is available from the lead contact upon request.

## ACKNOWLEDGMENTS

We thank Ellen Goodall, Jingjing Gao, Dawafuti Sherpa, Sichen (Susan) Shao and all Harper lab members for numerous discussions. Cryo-EM data collection was performed at the Harvard Cryo-EM Center for Structural Biology. Data processing was supported by SBGrid and O2 Cluster at Harvard Medical School. Mass Photometry was performed at the Center for Macromolecular Interactions (CMI). Confocal imaging was performed at the Core for Imaging Technology & Education Center (CITE). InsP_6_ detection by Mass Spectrometry was performed by Michael J. James at The Analytical Chemistry Core (ACC) at Harvard Medical School. This work was funded by Aligning Science Across Parkinson’s (ASAP) (J.W.H.), NIH R01NS110395 (J.W.H.), RO1NS083524 (J.W.H.), and NIH RO1 GM132129 (J.A.P.). Michael J Fox Foundation administers the grant ASAP-000282 on behalf of ASAP and itself. For the purpose of open access, the author has applied for a CC-BY public copyright license to the Author Accepted Manuscript (AAM) version arising from this submission. We acknowledge the Core for Imaging Technology and Education (Harvard Medical School) and the Cryo-EM Center for Structural Biology for imaging assistance.

## AUTHOR CONTRIBUTIONS

Conceptualization: T.F., J.W.H.; Investigation: T.F., C.L., F.V.H., J.A.P.; Analysis: T.F., C.L., J.W.H.; Writing—original draft: T.F., J.W.H.; Writing—reviewing and editing: T.F. and J.W.H. with input from all authors.

## DECLARATION OF GENERATIVE AI AND AI-ASSISTED TECHNOLOGIES IN THE WRITING PROCESS

During the preparation of this work, the authors used Harvard AI Sandbox to correct grammatical errors. After using this tool, the authors reviewed and edited the content as needed and take full responsibility for the content of the publication.

## METHODS

### KEY RESOURCES TABLE

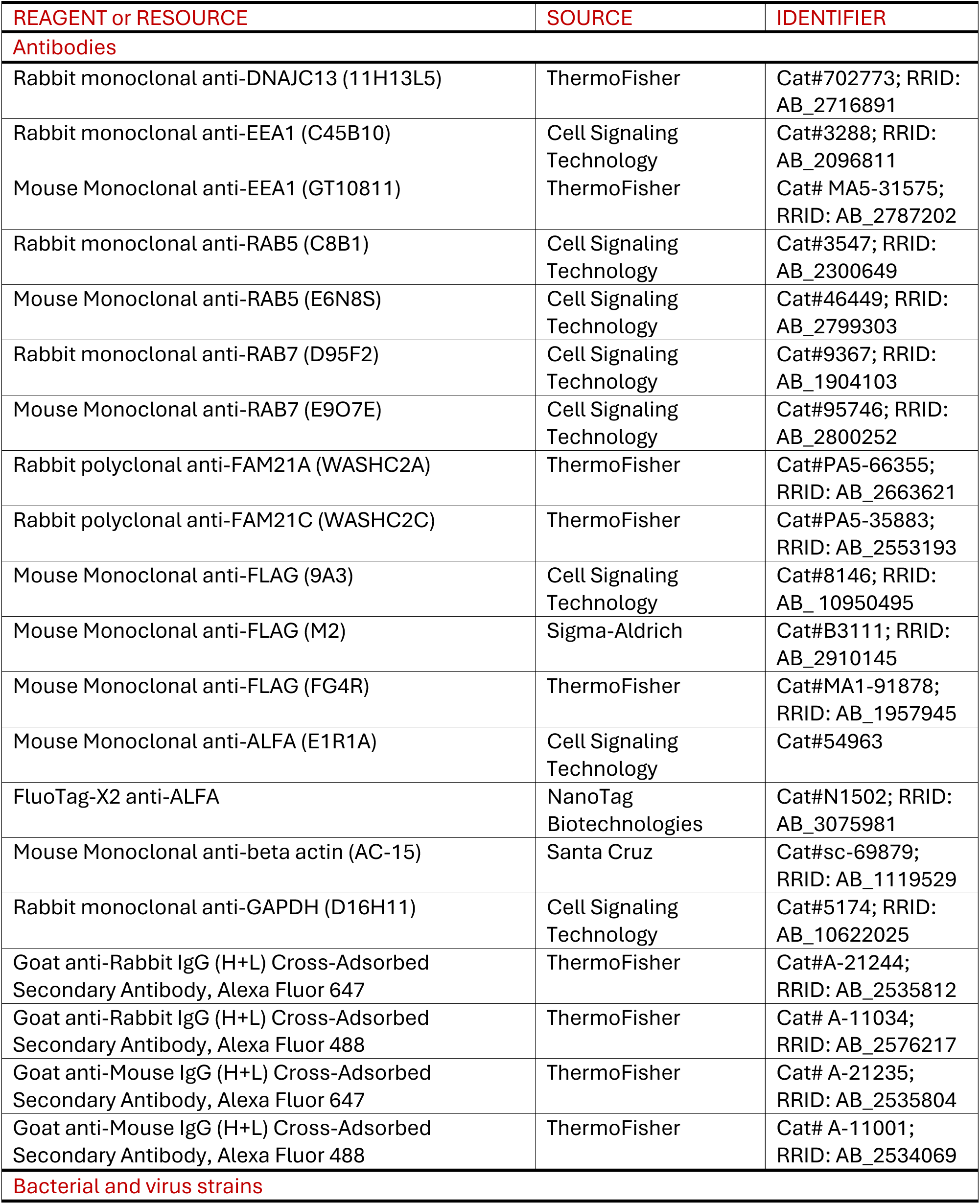
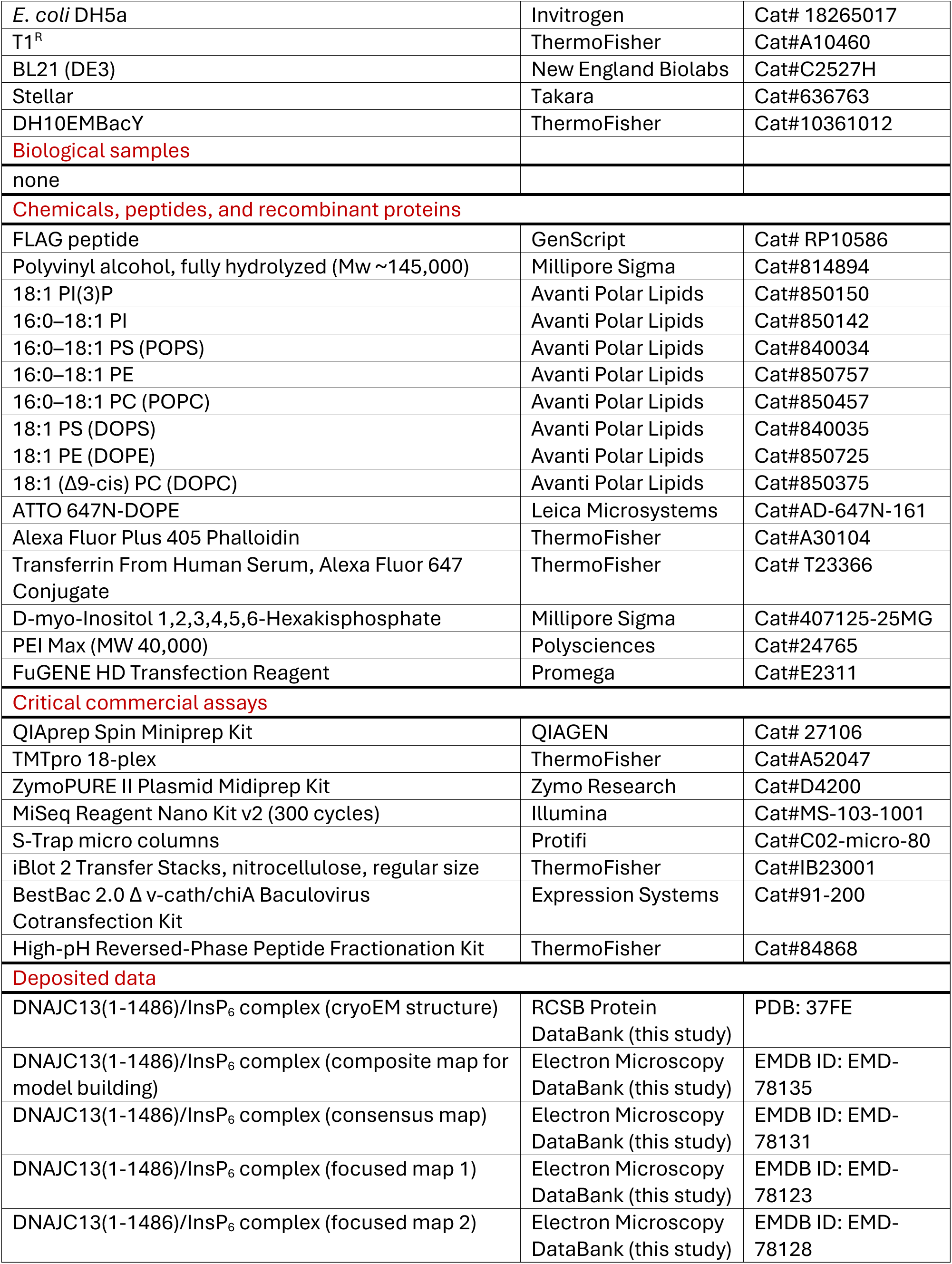
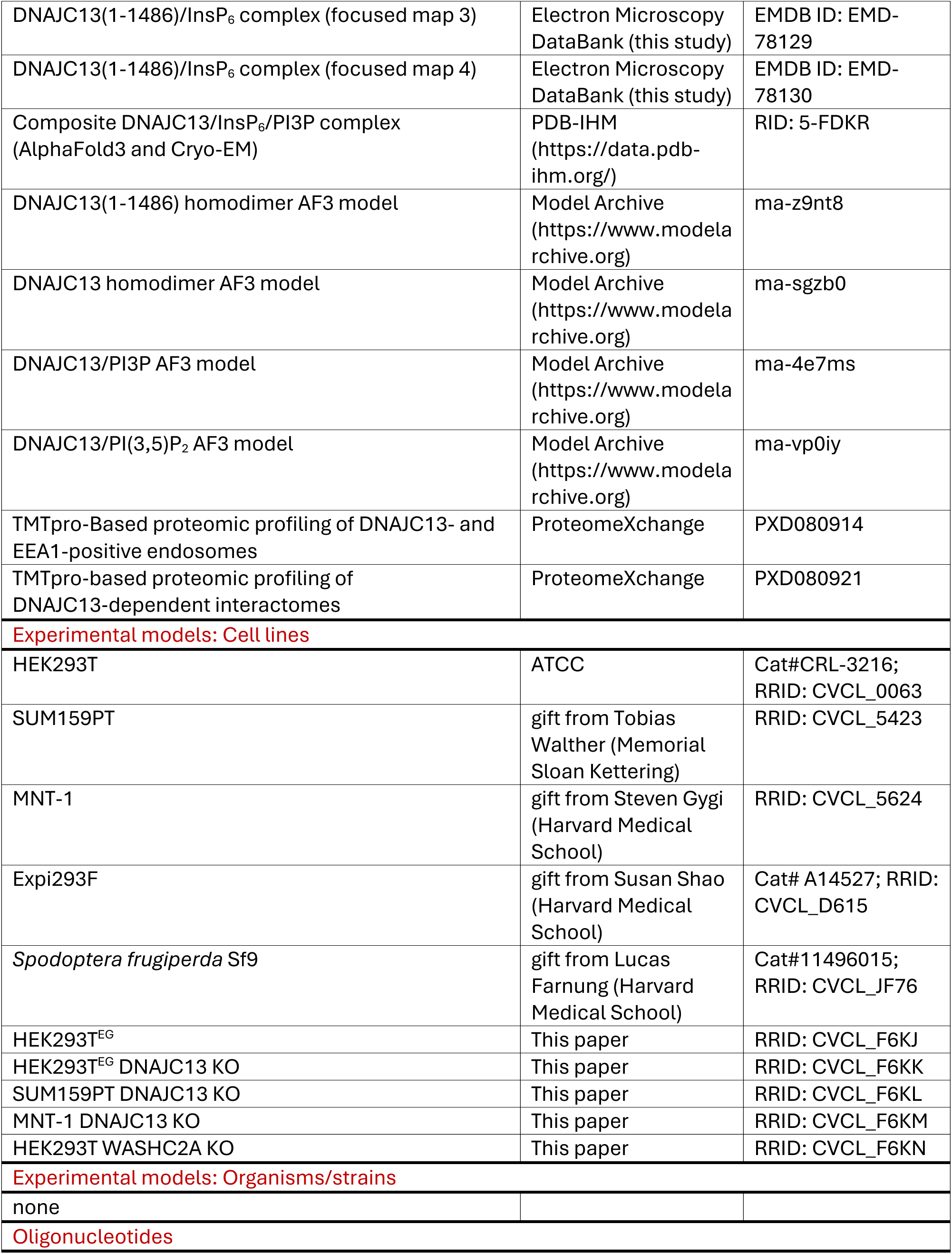
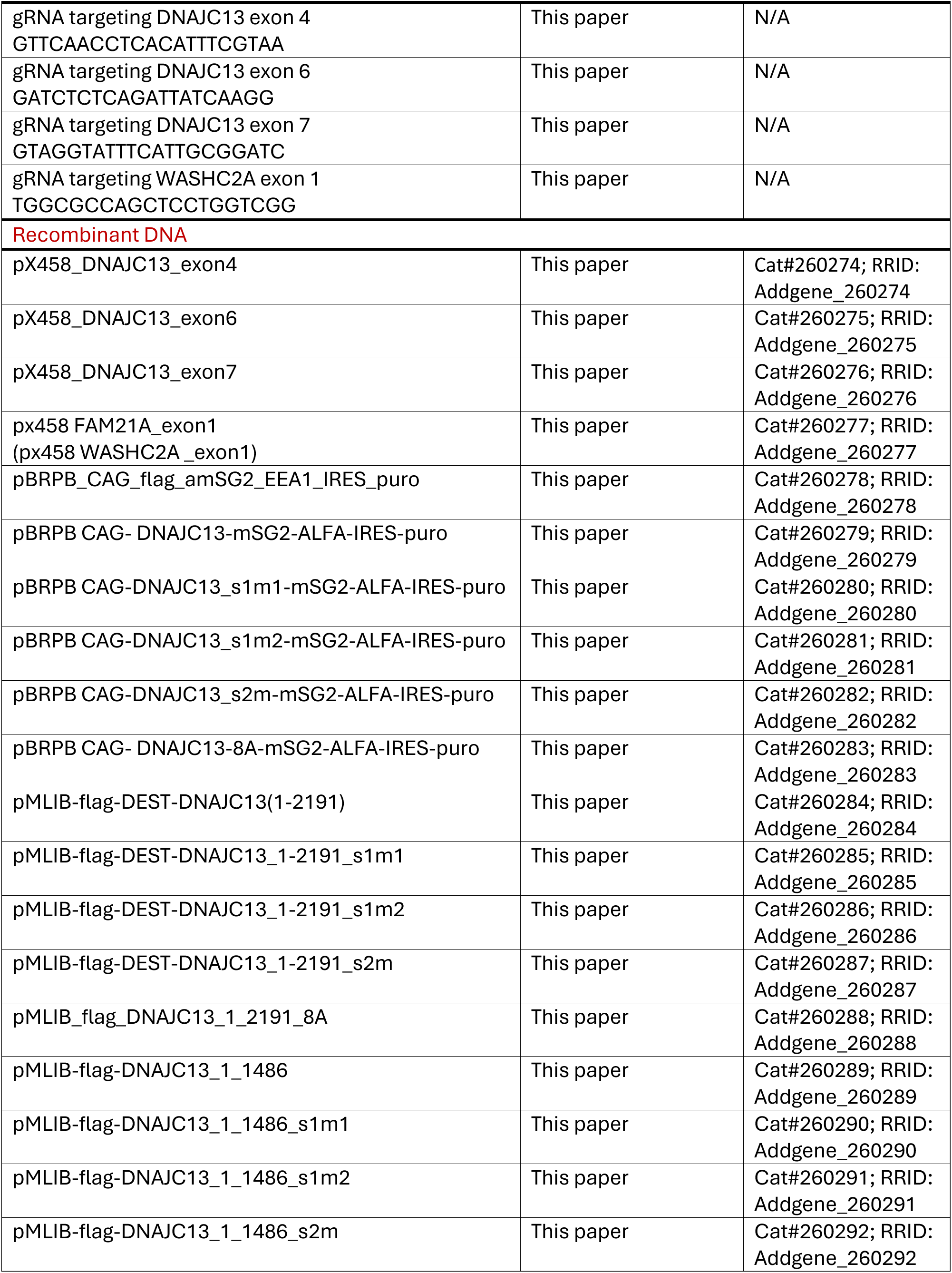
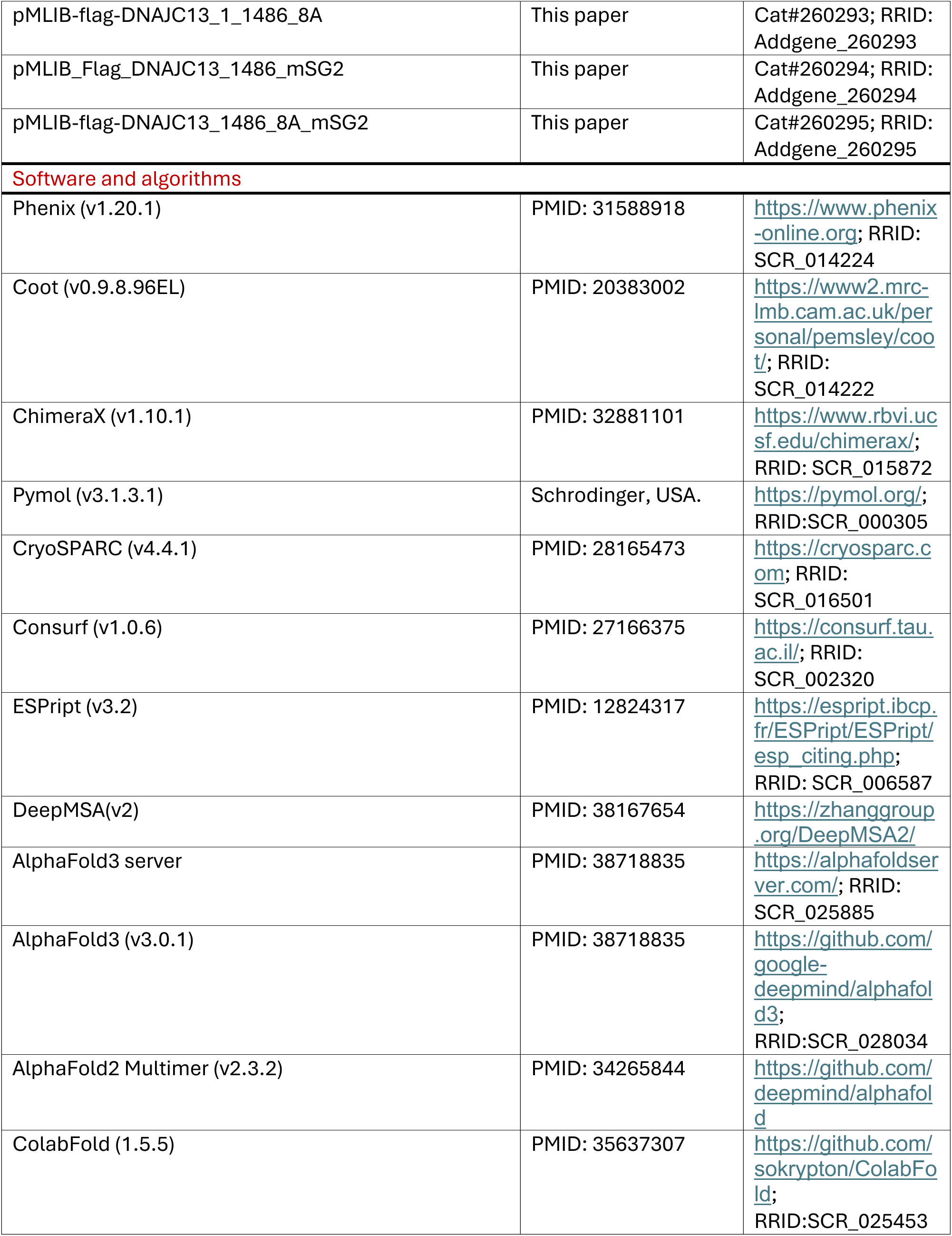
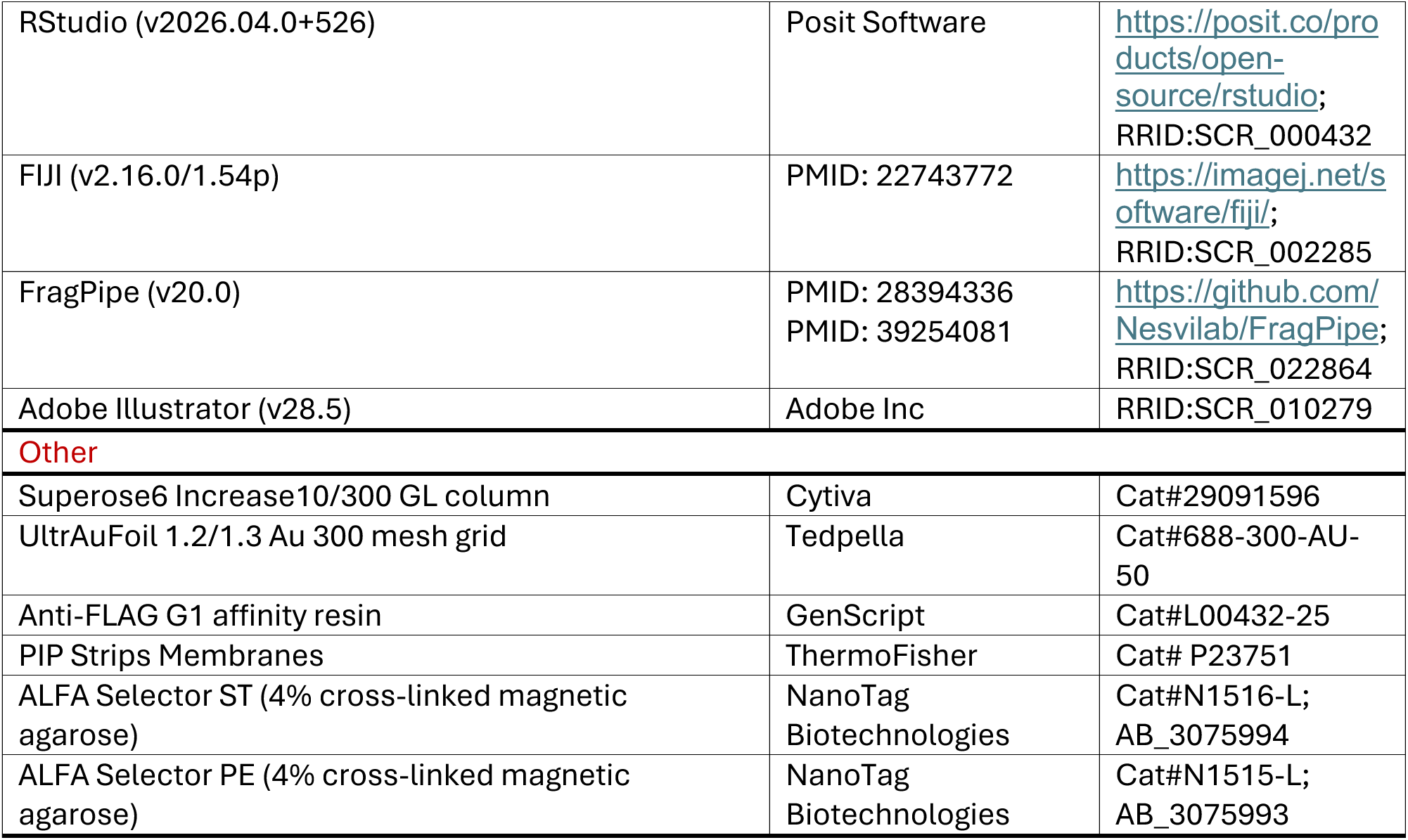

### EXPERIMENTAL MODEL AND STUDY PARTICIPANT DETAILS

#### Insect cells

*Spodoptera frugiperda* Sf9 cells (a kind gift from the laboratory of Lucas Farnung, Harvard Medical School) for baculoviral expression of recombinant proteins in insect cells were grown at 28 °C in ESF 921 medium (Expression Systems, Cat no. 96-001-01).

#### Bacterial cells

*Escherichia coli* strain BL21(DE3) was used for the overexpression of recombinant proteins. Bacterial cultures were initially grown at 37 °C and protein expression was induced with 0.8 mM isopropyl β-D-1-thiogalactopyranoside (IPTG). Following induction, the incubation temperature was immediately reduced to 18 °C, and cultures were maintained for an additional 20 h prior to harvesting.

#### Cell lines

HEK293T (ATCC, Cat. no. CRL-3216) and MNT-1 (a kind gift from the laboratory of Steven P. Gygi, Harvard Medical School) human cell lines were maintained in Dulbecco’s Modified Eagle Medium (DMEM; ThermoFisher Scientific, Cat. no. 11995065) supplemented with 10% (v/v) fetal bovine serum (FBS; Cytiva, Cat. no. SH30070.03), 100 µg/mL streptomycin, and 100 U/mL penicillin (ThermoFisher Scientific, Cat. no. 15140122) under standard humidified conditions at 37 °C with 5% CO_2_. SUM159PT human breast cancer cells (a kind gift from the laboratory of Tobias Walter, Memorial Sloan Kettering Cancer Center) were maintained in DMEM/F-12 medium (ThermoFisher Scientific, Cat no. 11330032) supplemented with 5 µg/mL insulin (Santa Cruz Biotechnology, Cat no. sc-360248), 1 µg/mL hydrocortisone (Sigma, Cat no. H0888-1G), 5% (v/v) FBS (Cytiva, Cat no. SH30070.03), 100 µg/mL streptomycin, and 100 U/mL penicillin (ThermoFisher Scientific, Cat no. 15140122). Expi293F human suspension cells (a kind gift from the laboratory of Susan Shao, Harvard Medical School) were maintained in Expi293 Expression Medium (ThermoFisher Scientific, Cat no. A1435101). Cultures were incubated in a regulated shaker incubator at 37°C with 8% CO_2_ and 70% humidity under continuous agitation at 125 rpm. All parental and engineered stable mammalian cell lines used in this study were routinely verified to be free of Mycoplasma contamination via MycoAlert Mycoplasma Detection Kit (Lonza, Cat no. LT07-318).

## METHOD DETAILS

### Molecular cloning

Plasmids used in this study were generated as previously described (see protocols.io: https://dx.doi.org/10.17504/protocols.io.5jyl8p7n8g2w/v1). Full-length cDNAs encoding human *DNAJC13* (wild type) and the *DNAJC13* 8A mutant were synthesized by Twist Bioscience (San Francisco, CA). Full length DNAJC13 and its corresponding truncation variants were cloned into pDONR223 or pDONR221 vectors using Gateway Cloning technology (ThermoFisher Scientific) to generate corresponding entry clones. Entry clones of EEA1 and WASHC2A were obtained from the human ORFeome v8.1 collection. Destination expression constructs were engineered using Gateway technology (ThermoFisher Scientific), Gibson assembly (New England Biolabs) or ClonExpress II One Step Cloning Kit (Vazyme), depending on the target vector architecture. For the generation of engineered stable cell lines, target sequences were transferred into modified piggyBac transposon expression vectors. For transient expression assays, mammalian expression constructs were generated within pcDNA3.1 or pMLIB vector backbones. Plasmids generated and used in this study have been deposited to Addgene and are listed in the key resource table.

### Baculovirus Generation and Recombinant Protein Expression in Insect Cells

*Spodoptera frugiperda* Sf9 cells culture were maintained as previously described (see protocols.io: https://dx.doi.org/10.17504/protocols.io.rm7vzyyrrlx1/v1). To generate recombinant baculoviruses, Sf9 cells were seeded into 6-well plates at a density of 1 × 10^6^ cells/well in a total volume of 2 mL of ESF 921 Insect Cell Culture Medium (Cat no. 96-001-01, Expression Systems). Recombinant baculoviruses were generated utilizing the BestBac 2.0 Δ v-cath/chiA Baculovirus Cotransfection Kit (Expression Systems, Cat no. 91-200) according to the manufacturer’s instructions. Briefly, linearized baculovirus DNA and the pBacPAK-DNAJC13(1-1486) plasmid were co-transfected into adherent Sf9 cells using the Expres^2^ TR transfection. Following incubation at 28 °C for 5 days, the conditioned culture medium containing the P0 generation virus was harvested. To amplify viral infectivity and titer, the harvested P0 virus was used to inoculate a 40 mL suspension culture of Sf9 cells at a density of 1 × 10^6^ cells/ml. At 72 h post-infection, the P1 viral generation was collected by pelleting the cells at 500 × *g* for 5 min and reserving the supernatant. The harvested P1 virus was subsequently used to infect a second 40 mL suspension culture of Sf9 cells maintained at 2 × 10^6^ cells/mL. Cultures were incubated for 72 h to yield the P2 generation stock. For each viral generation, infection efficiency was monitored by tracking cell diameters via a Countess 3 Automated Cell Counter (ThermoFisher Scientific, Cat no. AMQAX2000). Uninfected Sf9 cells typically exhibited an average diameter of approximately 16 μm, whereas fully infected cells swelled to a characteristic diameter of approximately 20 μm, indicating high titer baculovirus propagation. Viral amplification cycles were repeated sequentially until cell diameters reached this target threshold. All generated viral stocks were stored at 4 °C and protected from light. For large-scale protein expression, 500 μL of high-titer P2 or P3 baculovirus stock was added to a 50 mL suspension culture of Sf9 cells (2 × 10^6^ cells/mL), representing a 1:100 volumetric dilution. Cells were harvested by centrifugation at 72 h post-infection, and the resulting pellets were stored at -80 °C for downstream protein purification.

### Recombinant protein expression in mammalian cells

Recombinant protein expression in mammalian cells was performed as previously described (see protocols.io: https://dx.doi.org/10.17504/protocols.io.bp2l61x1kvqe/v1). Human embryonic kidney suspension Expi293F cells (a kind gift from the laboratory of Susan Shao, Harvard Medical School) were cultured and maintained in Expi293 Expression Medium (ThermoFisher Scientific, Cat no. A1435101) using sterile 125 mL Corning Erlenmeyer cell culture flasks (Sigma, Cat no. CLS431143-1EA). For routine subculturing, cells were systematically seeded at a baseline density of 2 × 10^6^ cells/mL and incubated at 37 °C in a humidified shaker containing 8% CO_2_ under continuous agitation at 125 rpm. High-purity expression plasmids were prepared at a large scale utilizing the ZymoPURE II Plasmid Midiprep Kit (Zymo Research, Cat no. D4200). Linear polyethylenimine “Max” (PEI MAX, MW 40,000; Polysciences, Cat no. 24765) prepared as 1 mg/mL sterile aqueous stock solution was used as the transfection reagent. For a standard 500 mL suspension culture adjusted to a working density of 2 × 10^6^ cells/mL, a 1.5 mg of total purified plasmid DNA and 4.5 mL of PEI MAX stock solution were diluted separately into two individual 25 mL volumes of Expi293 medium. These solutions were then combined to form a 50 mL transfection mixture and incubated at room temperature for 10-15 min to facilitate stable polyplex assembly. The resulting transfection mixture was subsequently added directly into the 500 mL cell culture under gentle, continuous swirling to ensure homogeneous polyplex distribution. Cells were harvested 72 h post-transfection by centrifugation at 1,000 × *g* for 3 min. The resulting cell pellets were immediately flash-frozen and stored at -80 °C for protein purification.

### FLAG-Affinity Protein Purification

Following harvest, cell pellets expressing FLAG-tagged recombinant proteins from bacteria, insect or mammalian expression systems were resuspended in ice-cold PBS. Cell lysis was performed on ice using a Qsonica sonicator (Cat no. Q500-110) equipped with a standard probe, utilizing a pulsed cycle of 10 s on and 20 s off. Sonicator settings were optimized for each host system as follows: bacterial suspensions were disrupted at 60% amplitude for 5 min; insect cells at 50% amplitude for 3 min; and mammalian cells at 40% amplitude for 1 min. To clarify the crude lysates, a two-step sequential centrifugation workflow was employed. Lysates were initially cleared by centrifugation at 4,000 × *g* for 10 min at 4 °C. The resulting supernatants were subsequently collected and subjected to high-speed centrifugation at 12,000 × *g* for 10 min at 4 °C to completely pellet remaining insoluble debris. Anti-FLAG G1 Affinity Resin (GenScript, Cat no. L00432-25) was equilibrated in gravity-flow columns by washing three times with 10 bed-volumes of PBS. The clarified supernatants were immediately loaded onto the columns and recirculated through the resin bed at least three times at 4 °C to ensure maximal protein binding. To remove non-specifically bound proteins, the columns were washed extensively a minimum of three times with 10 bed-volumes of ice-cold PBS. Target proteins were subsequently eluted by three sequential incubations with 5 mL of 0.1 mg/mL FLAG peptide (GenScript, Cat no. RP10586) diluted in PBS. The final eluates were pooled, collected, and concentrated utilizing centrifugal ultrafiltration filters for downstream applications.

### Size Exclusion Chromatography

Size exclusion chromatography (SEC) was performed on an ÄKTA Pure system (Cytiva) as previously described^54^, with minor modifications. Briefly, purified DNAJC13 (1-2191) proteins were loaded onto a Superose 6 Increase 10/300 GL column (Cytiva, Cat no. 29091596) equilibrated with an SEC buffer consisting of 20 mM HEPES (pH 7.5), 100 mM NaCl, and 0.002% (w/v) lauryl maltose neopentyl glycol (LMNG; Anatrace, Cat no. NG310). Peak fractions were pooled, concentrated to 1.0 mg/mL, and immediately utilized for cryo-EM grid preparation or mass photometry.

### Mass photometry

Mass photometry experiments were performed at the Center for Macromolecular Interactions (CMI) at Harvard Medical School using a Refeyn TwoMP mass photometer equipped with an Accurion vibration-isolation bench (Refeyn Ltd., UK). The instrument was powered on and allowed to equilibrate at room temperature for 45-60 min prior to data acquisition. Glass coverslips were alternately washed with isopropanol and ultra-pure water for three complete cycles before insertion into the instrument holder. Mass calibration was performed using a standard protein mix provided by CMI containing thyroglobulin (3 µM) and bovine serum albumin (BSA, 10 µM) in PBS supplemented with 5% glycerol. Data was collected using Refeyn AcquireMP software, and subsequent data analysis was performed using Refeyn DiscoverMP software. For each measurement, light-scattering events resulting from single-molecule binding at the glass-water interface were recorded in a 1-min movie. The calculated molecular weights of individual particles were plotted as mass histograms and quantified by fitting the peaks to a Gaussian distribution.

### Cryo-EM grid preparation and data collection

Cryo-EM grid preparation was performed essentially as previously described (see protocols.io: https://dx.doi.org/10.17504/protocols.io.n2bvj3dexlk5/v1). Specifically, a total of 3 μL of 1mg/ml DNAJC13(1-2191) purified as above was vitrified in ethane on glow-discharged UltrAuFoil R1.2/1.3 grids (Quantifoil) using a Vitrobot Mark IV (ThermoFisher Scientific) set at 22°C and 100% humidity with a 5 second wait time, 20 second blot time, and +5 blot force. Grids were screened for suitable ice using a 200 kV Talos Arctica equipped with a Gatan K3 direct electron detector. The dataset was collected using a Titan Krios (ThermoFisher Scientific) operating at 300 kV and equipped with a imaging filter with a 10-eV slit width and a Falcon4i direct electron detector (ThermoFisher Scientific) in counting mode at a nominal magnification of 165,000× corresponding to a calibrated pixel size of 0.736 Å. Semi-automated data collection was performed with EPU. For the DNAJC13(1-2191) dataset, a 3.28-second exposure was fractionated into 56 frames, resulting in a total exposure of 50.01 e-/Å2. The defocus targets were -1.0 to - 2.2 μm. Movies on the Titan Krios were collected in TIFF format.

### Cryo-EM data processing and model building

Data processing for this dataset was performed in CryoSPARC v4.4.1 (**Data S1.1**).^55,56^ 21786 movies were imported into CryoSPARC and were processed using patch motion correction followed with patch CTF estimation, and micrographs with severe contamination or poor CTF fits were removed. Particles were initially picked using the blob picker tool with a minimum particle diameter of 200 Å and a maximum particle diameter of 250 Å. Picked particles were extracted and underwent several rounds of 2D classification. High quality 2D classes were selected and used as templates for a further round of particle picking using CryoSPARC’s template picker tool. 18551 micrographs were subjected to automated particle picking using templates generated from blob picking. These newly picked particles were extracted with a box size of 512 pixels and down sampled to a box size of 64 pixels for the initial classification steps. The initial extracted particles underwent several rounds of heterogeneous refinement using multiple reference volumes generated initially by ab initio reconstruction. The best class containing 660,405 particles were selected for another round of heterogeneous refinement and full extract at box size of 512 pixels followed with homogeneous refinement, global CTF refinement, reference-based motion correction and non-uniform refinement produce a reconstruction (599,851 particles) at an overall resolution of 2.39 Å. 3D Variability Analysis (3DVA) and 3D Flexible Refinement (3DFlex) were both performed in CryoSPARC on the consensus particle stacks. The first component from 3DVA is shown in Supplementary Movie 1. 3D classification with different focused masks were carried out and additional non-uniform refinement and local refinement were performed, leading to four locally refined maps at 2.5-2.7 Å resolution. Finally, these refinements were combined to generate a composite map used for model building. An initial model for the DNAJC13 was from AlphaFold3 prediction. The model was manually adjusted in Coot v.0.9.8 in between iterative rounds of real space refinement in Phenix v.1.19.2. Figures were made using ChimeraX v.1.10.1 and PyMOL v3.1.3.1.

### AlphaFold modeling

For DNAJC13, structures of the DNAJC13(1-1486) dimer, full-length DNAJC13 dimer, and DNAJC13 monomer without ligand were generated using AlphaFold2-Multimer (v2.3.2) on the O2 High Performance Compute Cluster at Harvard Medical School and/or the AlphaFold3 server implemented on the Google platform, as indicated in the figure legends. Protein models containing PI(3)P or PI(3,5)P_2_ were generated using AlphaFold3 on the O2 High Performance Compute Cluster at Harvard Medical School.

### CRISPR-Cas9 gene editing

To generate DNAJC13 and WASHC2A knockout (KO) cell lines in HEK293T and SUM159 backgrounds, cells were transiently transfected with a pX458 expression vector (Addgene, Cat no. 48138) encoding Cas9 and a target-specific single guide RNA (sgRNA). Transfections were performed using polyethylenimine (PEI) or FuGENE HD (Promega, Cat no. E2311) transfection reagent according to the manufacturers’ instructions. Following a 24 h incubation, GFP-expressing cells were isolated via fluorescence-activated cell sorting (FACS), and single cells were deposited into individual wells of 96-well plates containing full growth medium. Single-cell clones were subsequently expanded and screened for successful gene ablation. For the MNT-1 cell line, DNAJC13-KO clones were generated via lentiviral transduction. Lentiviral particles were packaged using the lentiCRISPRv2-Opti backbone (Addgene, Cat no. 163126). Stably transduced MNT-1 cells were selected by incubation with 2 µg/mL puromycin for 4-5 days. Puromycin-resistant cells were then single-cell sorted into 96-well plates via FACS. All expanded single-cell clones from HEK293T, SUM159, and MNT-1 backgrounds were initially screened and confirmed by next-generation sequencing using the Illumina MiSeq platform, and target protein depletion was further validated via immunoblotting (**Data S2.2**).

The following sgRNA sequences were used to target the indicated gene:

DNAJC13, 5′-TTCAACCTCACATTTCGTAA-3′ (exon4), 5′-TAGGTATTTCATTGCGGATC-3′ (exon7); WASHC2A, 5′- TGGCGCCAGCTCCTGGTCGG-3′ (exon1).

### *In Vitro* InsP_6_-induced Cleavage Assay

To evaluate endogenous InsP_6_ bound to DNAJC13, autocatalytic cleavage assays were performed utilizing the *Clostridium difficile* Toxin A glucosyltransferase-cysteine protease domain (g-CPD) as a molecular biosensor, adapting the InsP_6_-induced cleavage assay described previously^40^ and detailed in protocols.io (DOI: 10.17504/protocols.io.e6nvwxddzgmk/v1). Recombinant DNAJC13(1-1486) wild-type (WT) and mutant proteins were purified from mammalian suspension cultures. Initial protein stocks were measured via NanoDrop spectrophotometry and systematically normalized to equivalent molar concentrations based on relative band intensity quantified from SDS-PAGE Coomassie staining or immunoblotting. Time-course autoprocessing reactions were executed at 37 °C across distinct experimental conditions maintaining identical final sample volumes and a 1:1 molar ratio of the g-CPD biosensor to the respective DNAJC13 variants. Aliquots were withdrawn at designated intervals (t = 0, 20, 40, 60, and 80 min), immediately quenched with 4 × Laemmli sample loading buffer, and thermally denatured at 85°C for 5 min. Equal volumes of the resulting denatured fractions were resolved via SDS-PAGE and transferred to membranes for western blot analysis. SDS-PAGE and western blot runs were executed in at least three technical replicates. For each individual experimental condition, intact g-CPD precursor and the cleaved CPD fragment were systematically normalized to the baseline at t = 0 min.

### Protein-Bound Inositol Hexaphosphate (InsP_6_) Extraction and LC-MS/MS Detection

Mass spectrometry-based detection of InsP_6_ was performed by Michael J. James at the Analytical Chemistry Core (ACC), Harvard Medical School. To quantify protein-bound inositol hexaphosphate (InsP_6_), equivalent molar amounts of purified WT and mutant DNAJC13(1-1486) recombinant proteins were thermally denatured at 85 °C for 10 min to liberate bound InsP_6_ from bound protein. Following denaturation, the released chemical fraction was isolated, and quantitative detection was executed utilizing a Vanquish Ultra-High Performance Liquid Chromatography (UHPLC) system coupled inline to an Exploris 240 Orbitrap mass spectrometer equipped with an electrospray ionization (ESI) source (ThermoFisher Scientific), following an adapted method described previously.^57^ Chromatographic separation of the target samples was achieved by injecting a 20 µL sample volume onto a Waters zHILIC hybrid column (2.1 × 100 mm) maintained at a regulated temperature of 45 °C. The binary mobile phase configuration consisted of 20 mM ammonium carbonate supplemented with 0.1% ammonium hydroxide in water (Mobile Phase A) and 95% (v/v) acetonitrile in water Mobile Phase B). Elution was performed at a continuous flow rate of 0.4 mL/min utilizing a linear gradient descending from 80% B to 10% B over a 5 min runtime window, followed by structured post-gradient washing and re-equilibration steps to eliminate carryover. For mass spectrometric detection, the ESI source was operated in negative ion mode with full scan MS acquisition range of 220-700 *m*/*z*. InsP_6_ chemical species were identified via high-resolution monitoring of the characteristic doubly charged precursor ion [M-2H]^2-^ at 328.9229 *m*/*z*. For tandem mass spectrometry (MS/MS or MS^2^) product ion scanning, fragmentation collision energies were pre-optimized via direct syringe infusion of a synthetic InsP_6_ chemical reference standard (Sigma, Cat no. 407125-25MG). Under identical chromatographic running parameters, the precursor coordinate at 328.9229 *m*/*z* was isolated and subjected to subsequent Product Ion Scans utilizing an optimized Normalized Collision Energy (NCE) of 30%. Absolute concentrations of liberated InsP_6_ were calculated by interpolating raw peak area integrations against a concurrent five-point external standard calibration curve run systematically alongside each experimental sample batch.

### PIP strip overlay assay

Phosphoinositide binding of DNAJC13 was assessed using nitrocellulose-based PIP Strips (ThermoFisher Scientific, Cat no. P23751) according to the manufacturer’s instructions. PIP Strips were equilibrated in Tris-buffered saline containing 0.1% (v/v) Tween-20 (TBS-T; 10 mM Tris-HCl pH 8.0, 150 mM NaCl, 0.1% Tween-20) for 10 min and then blocked for 1 h in TBS-T supplemented with 3% (w/v) fatty acid-free BSA (TBS-T/3% BSA) with gentle rocking. Blocked membranes were incubated with purified FLAG-DNAJC13 proteins diluted to 0.5 μg/mL in fresh TBS-T/3% BSA overnight at 4 °C with gentle agitation. Following incubation, membranes were washed three times for 5 min each in TBS-T. Membranes were incubated for 2 h in TBS-T/3% BSA containing mouse anti-FLAG conjugated with HRP primary antibody. After three additional 10-min washes in TBS-T, signals were visualized by enhanced chemiluminescence and imaged on a Bio-Rad ChemiDoc imager.

### SDS-PAGE immunoblotting and quantitative analysis

Immunoblotting was performed essentially as previously described (see protocols.io: https://dx.doi.org/10.17504/protocols.io.n2bvje64wgk5/v1). Briefly, cell pellets were resuspended in RIPA lysis buffer (ThermoFisher Scientific, Cat no. 89901) supplemented with Benzonase (Millipore, no. 71205-3). BCA assay kit (ThermoFisher Scientific, Cat no. 23227) was used to determine protein concentration, and normalized lysate amounts were boiled in 1× SDS containing Laemmli sample buffer. Lysates were run on 4-20% Criterion TGX Stain-Free Protein Gel (Bio-Rad, Cat no. 5678095). After electrophoresis, gels were electro-transferred onto a nitrocellulose membrane by using iBlot 2 Gel Transfer Device (ThermoFisher Scientific, Cat no. IB21001) and iBlot™ 2 Transfer Stacks (ThermoFisher Scientific, Cat no. IB23001). Membranes were blocked with 5% non-fat milk for 1 h, and incubated with primary antibody overnight at 4 °C and subsequently with HRP-conjugated or fluorescently-labeled secondary antibodies for 1 h at room temperature. After washing, blot images were acquired in a Bio-Rad ChemiDoc imager. Images were processed with Bio-Rad Image Lab software (version 6.1.0; RRID:SCR_014210). Western blot quantification analysis were performed in FIJI by using macro script as described: dx.doi.org/10.17504/protocols.io.7vghn3w. Quantified western blots are the mean of 3 technical repeats, with statistical analysis performed using RStuido. Full, uncropped versions of blots are provided in **Document S2**.

### Transferrin uptake assays

For endocytic cargo internalization assays, adherent SUM159 human breast cancer cells were incubated at 37 °C in complete culture medium supplemented with 20 µg/mL of Alexa Fluor 647-conjugated Transferrin (ThermoFisher Scientific, Cat no. T23366) for the time durations indicated in the respective figure legends. Following the internalization pulse, the endocytic reaction was immediately terminated by rapidly rinsing the cell monolayers with PBS, followed by the immediate addition of 4% (w/v) paraformaldehyde (PFA) in PBS. Cells were fixed for 15 min at room temperature. Following fixation, cell monolayers were washed three times with PBS and processed for downstream immunofluorescence and confocal imaging analysis as described in the Immunofluorescence and Live-Cell Imaging method sections.

### Immunofluorescence and Live-Cell Imaging

Spinning disk confocal imaging was performed at the Core for Imaging Technology & Education (CITE), Harvard Medical School, essentially as previously described (see protocols.io: https://dx.doi.org/10.17504/protocols.io.bp2l6jyj1vqe/v1). For immunofluorescence staining, cells were fixed with 4% (w/v) paraformaldehyde (PFA) in PBS for 20 min at room temperature and permeabilized with 0.1% (v/v) Triton X-100 in PBS for 15 min. After three washes with PBS, cells were blocked with 2% (w/v) BSA in PBS for 60 min at room temperature. Primary antibodies were diluted 1:200 in 1% (w/v) BSA in PBS, and 200 μL of the antibody solution was added to each well for overnight incubation at 4°C. Following primary antibody incubation, cells were washed three times with PBS containing 0.05% (v/v) Tween-20 (PBST). Cells were then incubated with 200 μL of Alexa Fluor-conjugated secondary antibodies diluted 1:200 in 1% BSA in PBS for 1 h at room temperature in the dark. After three additional washes with PBST, nuclei were counterstained with 5 μg/mL Hoechst 33342 for 5 min. Samples were maintained in PBS and imaged using a NIKON 60×/1.42NA oil objective on a Nikon Ti2 Eclipse microscope equipped with a Yokogawa CSU-X1 spining disk system. Fluorophores were excited sequentially utilizing a Nikon LUN-F XL solid-state laser combiner with 405 nm (80 mW), 488 nm (80 mW), 561 nm (65 mW), and 640 nm (60 mW) laser lines. Excitation light was directed through a Semrock Di01-T405/488/568/647 dichroic mirror. Fluorescence emission was collected using Chroma Technology bandpass filters: ET455/50m (for 405 nm ex), ET525/50m (488 nm ex), ET 620/60m (561 nm ex), and ET700/75m (640 nm ex). Images were acquired with a Hamamatsu ORCA-Fusion BT sCMOS camera controlled by NIS-Elements image acquisition software (version AR, RRID:SCR_014329). Laser intensities and exposure times were kept constant across all experimental groups. For live-cell imaging, cells were seeded onto sterile 24-well glass-bottom plates (Cellvis, Cat no. P24-1.5H-N) or µ-Slide 8-well glass-bottom chamber slides (ibidi, Cat no. 80807). During live-cell image acquisition, cells were maintained at 37°C and 5% CO_2_ using an OkoLab environmental control system and imaged using a Yokogawa CSU-X1 spinning disk system on a Nikon Ti2 Eclipse microscope equipped with a NIKON 60×/1.42NA oil objective. Fluorophores were excited sequentially using a Nikon LUN-F XL solid-state laser combiner with 405 nm (80 mW), 488 nm (80 mW), 561 nm (65 mW), and 640 nm (60 mW) laser lines. Images were acquired with a Hamamatsu ORCA-Fusion BT sCMOS camera controlled by NIS-Elements image acquisition software. Laser intensities and exposure times were kept constant across all experimental groups. Image processing was performed using ImageJ/Fiji. For comparative datasets, brightness and contrast adjustments were applied identically across experimental groups using the same display settings.

### Giant Unilamellar Vesicles (GUVs) Preparatioin and Recruitment Assays

Giant Unilamellar Vesicles (GUVs) were prepared using the hydrogel-assisted swelling method as previously described (see protocols.io: https://dx.doi.org/10.17504/protocols.io.3byl4b398vo5/v1). Briefly, PVA (MW 145,000; Sigma, no. 8148940101) was dissolved in ultra-pure water with heating at 90°C for 30 min to make a 5% (w/v) aqueous solution. A thin layer of the 5% PVA solution was applied to cover the bottom of sterile 12-well glass-bottom plates (Cellvis, no. P12-1.5H-N) and dried overnight in a chemical fume hood to form a uniform film. Lipid mixtures were prepared in chloroform at a total stock concentration of 1 mg/mL. Atto647N-labeled 1,2-dioleoyl-sn-glycero-3-phosphoethanolamine (Atto647N-DOPE; Leica, Cat no. AD 647N-161) was used as the GUV membrane dye and dissolved in a 80:20 chloroform/methanol mixture to a 1mg/ml stock concentration. All the other lipids for GUV preparation were purchased from Avanti Polar Lipids. For phosphatidylinositol 3-phosphate [PI(3)P]-containing GUVs, the lipid composition consisted of 70% (w/w) 1,2-dioleoyl-sn-glycero-3-phosphocholine (DOPC, Cat no. A80375C/0025/AM01M), 20% (w/w) 1,2-dioleoyl-sn-glycero-3-phosphoethanolamine (DOPE, Cat no. A80725C/0025/AM01M), 5% (w/w) 1,2-dioleoyl-sn-glycero-3-phospho-L-serine (DOPS, Cat no. A80035C/0010/AM01M), and 5% (w/w) 1,2-dioleoyl-sn-glycero-3-phospho-(1’-myo-inositol-3’-phosphate) (18:1 PI3P, Cat no. A85162P/0100/4V11U), supplemented with 0.3% (w/w) Atto647N-DOPE. For control (non-PI3P) GUVs, 18:1 PI3P was replaced with an equivalent weight ratio of 5% (w/w) 1-palmitoyl-2-oleoyl-sn-glycero-3-phosphoinositol (16:0-18:1 PI, Cat no. A85079P/0100/4V11U). For vesicle formation, 10-15 µL of the respective lipid mixture was uniformly deposited onto the dried PVA film. The solvent was evaporated under a stream of inert gas for 10 min to form a dry lipid film. The films were then hydrated by adding 200 µL of a 400 mOsm sucrose solution to each well. Swelling was allowed to proceed for 1 h at room temperature. The resulting GUVs were harvested by gentle pipetting and utilized within 12 h of preparation. GUV recruitment assays were performed essentially as previously described (see protocols.io: https://dx.doi.org/10.17504/protocols.io.bxm2pk8e). Protein membrane recruitment experiments were performed in 96-well glass bottom plates (Cellvis, P96-1.5H-N). Wells were coated with 2 mg/mL BSA for 30 min and washed three times with PBS buffer (pH 7.5). Recombinant DNAJC13(1-1486)-mSG2 or specified mutants were added to a final concentration of 80nM. To initiate the recruitment reaction, 10 µL of GUVs were added into each well to achieve a final reaction volume of 80 µL. After 5-min incubation, during which random fields of view were selected, time-lapse images were acquired at 37°C using a Yokogawa CSU-X1 spinning disk confocal on a Nikon Ti-E motorized microscope equipped with a Nikon Apochromat Plan 60×/1.40 N.A. immersion oil objective lens, and the Perfect Focus System was used to maintain focus over time. Consistent laser intensities and exposure times were maintained across all experimental groups. Technical replicates were performed at least three times for each experimental condition. Imaging processing, including identical brightness and contrast adjustments across comparative datasets, was performed using ImageJ/Fiji. To quantify fluorescence intensities on the GUV membrane, time-lapse dual-channel TIFF images were processed using a custom automated analysis pipeline written in Python 3. This pipeline was adapted and modified from the open-source GUV tracking framework developed by the laboratory of Johannes Schöneberg (https://github.com/JohSchoeneberg/GUV_Tracking_Quantification). Initial vesicle center coordinates (*x, y*) were supplied as seed inputs, and GUV tracking across frames was executed on the membrane dye channel. For each frame, a dynamic intensity threshold was calculated between the GUV center and the surrounding background. The GUV boundary was mapped by extracting orthogonal intensity profiles across the active coordinate center and computing the full-width at half-maximum boundaries. The detected vesicle radius was multiplied by a factor of 1.1 to encapsulate the total boundary, and a circular mask with a 12-pixel thickness was generated to extract the regions of interest (ROIs). Background intensities for the membrane and cargo channels were determined by pooling all pixels within 30×30 pixel regions at the four corners of the image frame. Net fluorescence intensities were calculated by subtracting these local background metrics from the mean ROI intensities. For kinetic analysis, values were normalized to the initial frame. Quality control diagnostics were output automatically as binary threshold masks and bounding-box overlays to verify spatial tracking fidelity. Individual intensity trajectories were calculated, averaged, and reported as mean ± SEM.

### Melanin quantification

Melanin was quantified by following a previously published method^58^ with minor modifications (see protocols.io: https://dx.doi.org/10.17504/protocols.io.yxmvmd339v3p/v1). Cells were plated in 6-well plates (2-wells per replicate), and cells were grown for an additional three days before harvest. On the day of harvest, cells were washed with cold DPBS and scraped into lysis buffer (1.5% Triton X-100 with protease inhibitors) and incubated on ice for 20 minutes with intermittent vortexing. Lysates were then centrifuged at 17,000 × *g* for 20 min at 4°C. Supernatants were transferred to a new tube and protein concentration was measured using BCA, and pellets containing melanin were washed in 100% ethanol by vortexing. After ethanol wash, pellet were spun down by centrifugation at 17,000 × *g* for 5 minutes at 4°C. Ethanol was discarded and samples were air-dried at 37°C for 30 minutes. Pellets were resuspended in 250 µL of 1M NaOH/10% DMSO and incubated at 80 °C for 1 h with occasional vortexing. After incubation, samples were cooled to room temperature and centrifuged at 17,000 × *g* for 10 min at room temperature. 200 µL of sample was transferred to a 96-well plate and absorbance at 490 nm was measured on a microplate reader.

### Affinity Purification of Endosomes (Endo-IP)

To enable the affinity-based isolation of specific endosomal populations, knockout cell lines were engineered to stably express ALFA-mSG2-EEA1 or DNAJC13-mSG2-ALFA using the PiggyBac transposon system. Organelle immunoprecipitation was performed as previously described (see protocols.io: https://dx.doi.org/10.17504/protocols.io.ewov14pjyvr2/v2). Briefly, cells were grown to confluence in 15 cm plates, and all subsequent manipulation steps were strictly executed on ice or at 4°C. Cells were rinsed with ice-cold PBS and harvested by gentle scraping, and collected by centrifugation at 500 × *g* for 5 min at 4 °C. Cells pellets were resuspended in ice-cold KPBS buffer (25 mM KCl, 38.4 mM KH_2_PO_4_, 61.6 mM K_2_HPO_4_, pH 7.25) supplemented with cOmplete protease and PhosSTOP phosphatase inhibitor cocktails. Cells were mechanically disrupted via 30 strokes in a 2 mL Type B Dounce tissue grinder (DWK Life Sciences, Cat no. 885300-0002). To yield the post-nuclear supernatant (PNS), the crude lysate was initially centrifuged at 500 × *g* for 3 min at 4°C to pellet unlysed cells and nuclei; the resulting supernatant was collected and further cleared by centrifugation at 1,000 × *g* for 5 min at 4°C. The cleared PNS fraction was then incubated with pre-washed magnetic anti-ALFA beads (NanoTag Biotechnologies, Cat no. N1516-L) for 50 min at 4 °C under constant end-to-end rotation. Endosome-bound beads were isolated via a magnetic rack and washed three times with ice-cold KPBS. Endosomes were eluted from the resin by incubation at 37 °C for 30 min in an elution buffer containing 20 mM HEPES (pH 7.5), 100 mM NaCl, and 0.5% (v/v) NP-40. The resulting eluates were subsequently processed for immunoblotting or quantitative proteomics.

### On-Column Proteolytic Digestion and TMTpro Labeling for Quantitative Proteomics

Protein samples from isolated endosomes were prepared using the S-Trap micro-spin column digestion protocol according to the manufacturer’s instructions (Protifi, Cat no. C02-micro-80). Tandem Mass Tag pro (TMTpro) multiplexing and peptide processing strategies were executed as previously described (see protocols.io: https://dx.doi.org/10.17504/protocols.io.rm7vzjej4lx1/v1). Briefly, each sample was mixed with an equal volume of 2× lysis buffer consisting of 10% (w/v) SDS and 100 mM triethylammonium bicarbonate (TEAB; pH 8.5). Proteins were reduced by incubation with 5 mM tris(2-carboxyethyl) phosphine (TCEP) at 55°C for 30 min and subsequently alkylated with 15 mM iodoacetamide for 30 min at room temperature in the dark. Samples were acidified by adding phosphoric acid and then mixed with S-Trap binding buffer (90% methanol, 100 mM TEAB, pH 7.55). The resulting mixture was loaded onto the micro-spin columns and washed four times with 200 µL of S-Trap binding buffer utilizing a manifold vacuum system. For proteolytic digestion, proteins were first digested on-column with 0.5 µg of Lys-C at 37°C overnight in a humidified chamber, followed by a subsequent 6 h incubation with 0.5 µg of trypsin. Peptides were sequentially eluted from the columns via three consecutive centrifugation steps: first with 50 mM TEAB (pH 8.5), followed by 0.2% (v/v) formic acid in ultra-pure water, and finally with 50% (v/v) acetonitrile. Eluates were pooled and dried to completeness using a SpeedVac vacuum concentrator. Dried peptides were reconstituted in 50 µL of 200 mM 4-(2-hydroxyethyl)-1-piperazinepropanesulfonic acid (EPPS; pH 8.0) containing 30% (v/v) acetonitrile and labeled with 5 µL of a 25 µg/µL TMTpro reagent stock for 1.5 h at 25°C. The labeling reactions were quenched by adding 0.5% (w/v) hydroxylamine for 15 min at room temperature. To ensure optimal and balanced peptide representation across multiplexed channels without sacrificing high-yield samples, a 3 µL aliquot from each channel was pooled in an equal volume ratio for an initial ratio-check analysis. Based on the ratio-check profile, the remaining samples were precisely pooled^59^: biological replicates within the same experimental group were adjusted using pro-rata volumes to achieve equivalent peptide loading, whereas systematic differences in total peptide yields between distinct groups (e.g., negative controls versus treatment groups) were intentionally preserved to maintain high mass spectrometry sensitivity. The combined multiplexed peptide pools were then dried under vacuum. Multiplexed peptide pools were desalted using custom-packed C18 StageTips, dried, and finally resuspended in 5% (v/v) acetonitrile containing 5% (v/v) formic acid prior to liquid chromatography-tandem mass spectrometry (LC-MS/MS) analysis.

### Liquid Chromatography-Mass spectrometry (LC-MS) data acquisition and analysis

Quantitative proteomic analysis was performed essentially as previously described (see protocols.io: https://dx.doi.org/10.17504/protocols.io.bys6pwhe). Briefly, IP samples (label free) were analyzed using an Orbitrap Exploris480 mass spectrometer (ThermoFisher Scientific) coupled with an nLC-1200 liquid chromatograph. Peptides were separated on a 100μm microcapillary column packed with 20cm of Accucore C18 resin (2.6 μm, 150 Å, Thermo). A 90-min linear gradient from 5% to 20% ACN in 80min, to 36% at 83min and 98% at 85min in 0.125% formic acid was used at 0.3µL/min. MS1 spectrum was acquired on the Orbitrap (resolution 60,000, scan range 350-1,350 m/z, standard automatic gain control (AGC) target, auto maximum injection time). Peptide fragmentation was achieved by high-energy collisional dissociation (HCD) at 28% normalized collision energy. MS2 was acquired on the Orbitrap (resolution 15,000, isolation window 1.2 m/z, 200% normalized AGC, 96ms maximum injection time). FAIMS Pro was set to -30, -50 and -70V compensation voltages (CV). A cycle time of 1 sec is used for each CV. Raw data files were processed using FragPipe 23.1.

TMTpro-labeled samples were analyzed using an Orbitrap Eclipse mass spectrometer (ThermoFisher Scientific) coupled with a Vanquish Neo liquid chromatograph. Peptides were separated on a 100μm microcapillary column packed with 20cm of Accucore C18 resin (2.6 μm, 150 Å, Thermo). A 90-min linear gradient from 5% to 20% ACN in 80min, to 36% at 83min and 98% at 85min in 0.125% formic acid was used at 0.3µL/min. MS1 spectrum was acquired on the Orbitrap (resolution 60,000, scan range 350-1,350 m/z, 100% automatic gain control (AGC) target, auto maximum injection time). Peptide fragmentation was achieved by high-energy collisional dissociation (HCD) at 36% normalized collision energy. MS2 was acquired on the Orbitrap (resolution 50,000, isolation window 0.7 m/z, first mass 110 m/z, 200% normalized AGC, 120ms maximum injection time). FAIMS Pro was set to -40, -60 and-80V. A cycle time of 1 sec is used for each CV.

TMTpro-MS raw data files were processed using MSConvert^60^ and searched utilizing the Comet^61^ search engine against the human canonical database (UniProt Swiss-Prot, October 2024 archive), which included reversed decoy sequences and common contaminants. Peptide mass tolerance was set to 50 ppm and fragment ion tolerance to 0.03 Da. These search windows were optimized to maximize spectral matching sensitivity in conjunction with target-decoy linear discriminant analysis (LDA).^62^ Static modifications were designated as TMTpro (+304.207 Da) on lysine residues and peptide N-termini, and carbamidomethylation (+57.021 Da) on cysteine residues. Oxidation of methionine residues (+15.995 Da) was permitted as a variable modification. LDA was executed^63^ to filter peptide-spectrum matches (PSMs) to a 2% false discovery rate (FDR)^64^, and quantified proteins were required to have a minimum of 1 unique peptide. TMTpro reporter ion intensities were extracted by selecting the maximum peak intensity within a ±0.003 Da window around the theoretical *m*/*z* and corrected for isotopic impurities. To ensure high-quality quantification, PSMs were filtered to retain only those with a combined sum signal-to-noise ratio 200 across all multiplexed channels and a precursor isolation specificity/purity cutoff ≥ 50%.^45^ Downstream data summarization, missing value imputation (MBimpute = TRUE), and global normalization were executed using the MSstatsTMT R package.^65,66^ Protein-level summarization was conducted utilizing the default ‘msstats’ setting. Differential abundance testing was performed using a moderated *t*-test, and *p*-values were adjusted for multiple hypothesis testing using the Benjamini-Hochberg (BH) method.

## QUANTIFICATION AND STATISTICAL ANALYSIS

For quantitation of immunoblots, protein detection was performed using the ChemiDoc MP Imaging System with fluorescently labeled secondary antibodies. Densitometric values are presented as mean ± SD from three technical replicates and were quantified in ImageJ/FIJI(v2.16.0/1.54p). Co-localization analysis of fluorescently labeled proteins were performed in ImageJ/FIJI(v2.16.0/1.54p) using the JaCoP plugin. Statistical analyses were carried out in RStudio, and the specific tests used are indicated in each figure legend.

---

**Figure S1.**
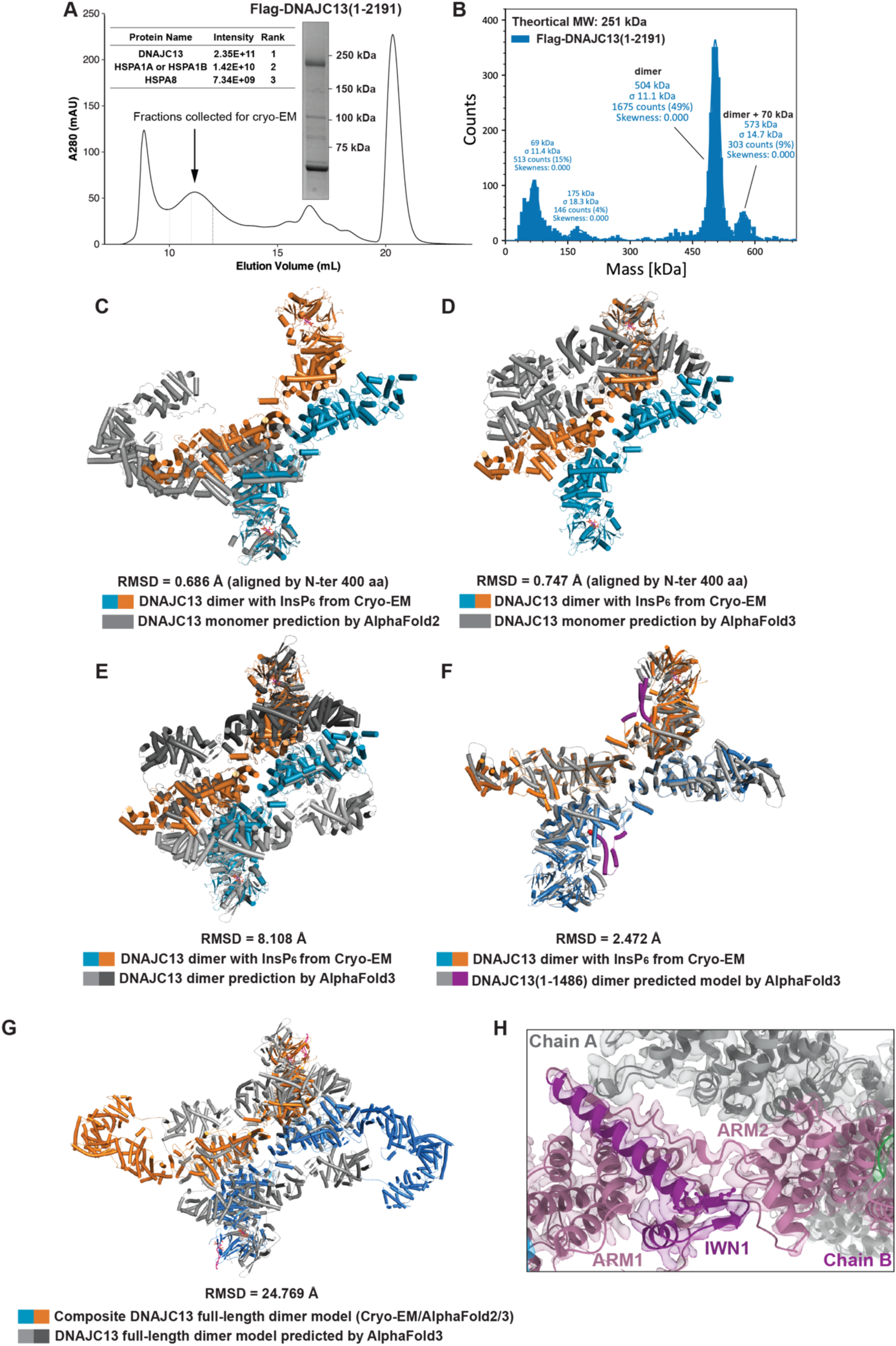
Structural Alignment and Comparative Analysis of the DNAJC13(1-1486) Cryo-EM Structure and AlphaFold Predictions, Related to Figure 1. (A) Size-exclusion chromatography profile of Flag-DNAJC13(1-2191) purified from Expi293F cells. UV absorbance at 280 nm is plotted against elution volume. Inset, Coomassie-stained SDS-PAGE of the peak fraction. Size markers indicate mass in kDa. Inset: Mass-spectrometry analysis of purified DNAJC13(1-2191) from Expi293F cells. Proteins identified listed with their label-free intensities and abundance rank. DNAJC13 is the most abundant species, with HSPA1A or HSPA1B and HSPA8 detected as the most abundant co-purifying chaperones. (B) Mass photometry of Flag-DNAJC13(1-2191) purified from Expi293F cells. The theoretical molecular weight of the monomer is 251 kDa. The major species corresponds to a dimer (504 kDa), with a minor population of dimer plus a 70-kDa complex, indicating that DNAJC13 forms a dimer in solution. (C-F) Structural overlays of the experimentally determined DNAJC13(1-1486) homodimer cryo-EM model (colored in blue and orange) with corresponding predictive structural models (colored in gray): (Panel C) AlphaFold2 predicted monomer model, (Panel D) AlphaFold3 predicted monomer model, (Panel E) AlphaFold3 predicted full-length dimer model, and (Panel F) AlphaFold3 predicted DNAJC13(1-1486) dimer model. (G) Structural overlays of the composite DNAJC13 homodimer model by merging Cryo-EM and AlphaFold2/3 (colored in blue and orange) with AlphaFold3 predicted DNAJC13 full-length dimer model. (H) Packing of the IWN1 motif, a fold apparently unique to DNAJC13, with the subsequent ARM2 domain. Structural elements in Chain B are shown in magenta while chain A is shown in grey.

**Figure S2.**
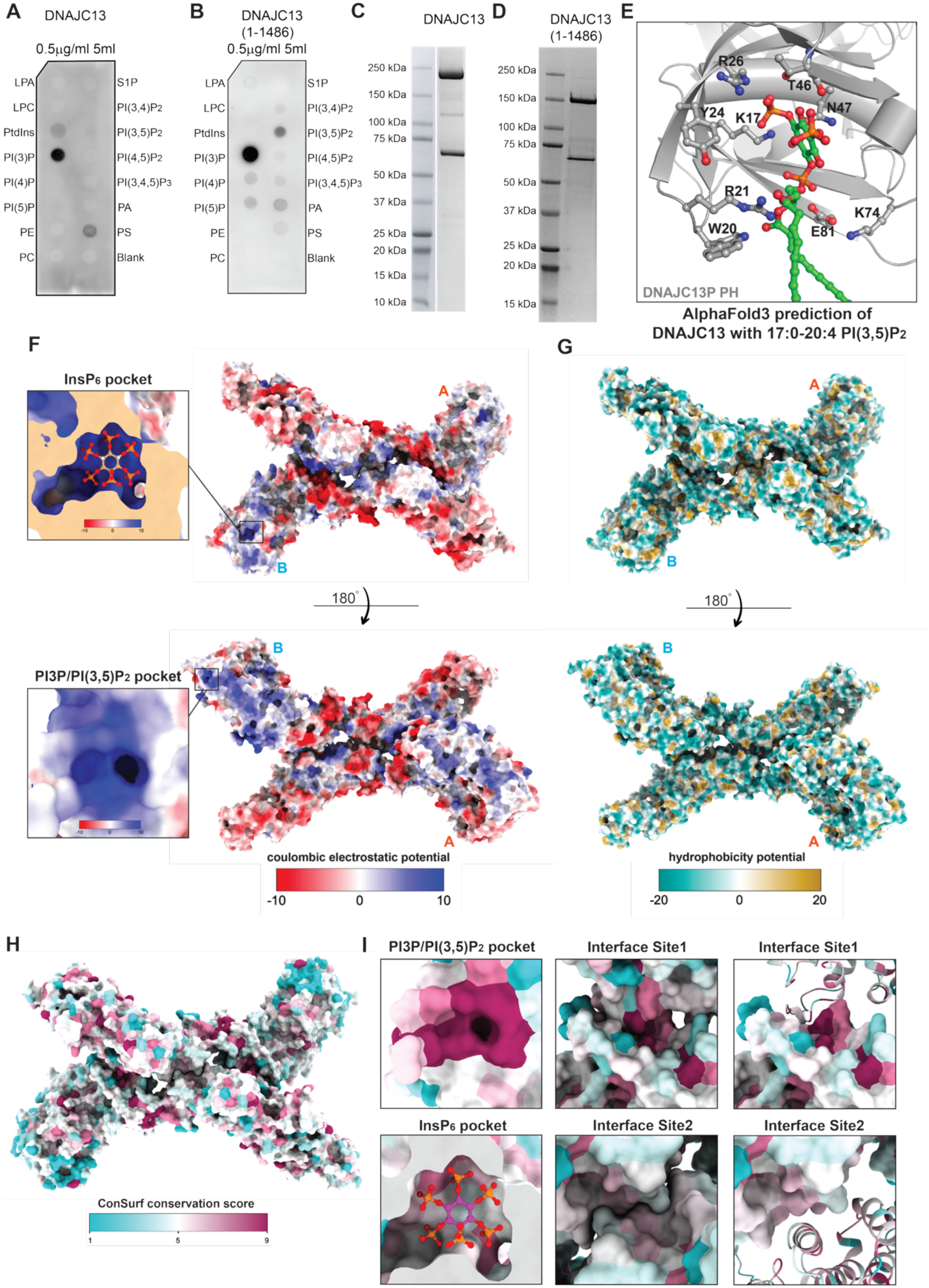
Surface properties, conservation, and phosphoinositide binding of DNAJC13, related to Figure 2. (A, B) Lipid-protein overlay assays (PIP strips) using full-length DNAJC13 (Panel E) or DNAJC13(1-1486) (Panel F). Proteins (0.5 µg/ml) were incubated with membrane-spotted lipids as indicated, and bound protein was detected by immunoblotting, revealing preferential binding to PI3P. (C, D) Coomassie-stained SDS-PAGE gels showing the purity of full-length DNAJC13 (Panel C) and DNAJC13(1-1486) (Panel D) used in the lipid-binding assays. (E) AlphaFold3 predicted model of the DNACJC13 with 17:0-20:4 PI(3,5)P_2_, illustrating the specificity of the PI3P-binding pocket. Key basic and polar residues forming the predicted binding are shown as sticks around the PI(3,5)P_2_ headgroup. (F) Coulombic electrostatic potential of the DNAJC13(1-1486) dimer cryo-EM model shown in two orientations related by 180°. Insets highlight the highly basic InsP_6_ pocket formed by PHL2-PHL3 and the basic pocket on PHL1 that binds PI3P. (G) Hydrophobicity surface of the same model and orientations as in (Panel A), colored from hydrophilic (cyan) to hydrophobic (yellow). (H) Surface representation colored by ConSurf conservation scores, from variable (cyan) to highly conserved (magenta). (I) Close-up views of conserved surface areas. Top left, conserved cavity corresponding to the PI3P pocket on PHL1. Bottom left, moderately conserved InsP_6_ pocket between PHL2 and PHL3. Top middle/right, highly conserved patches at dimer interface Site 1 between the two protomers (surface and ribbon views). Bottom middle/right, intermediately conserved patches at dimer interface Site 2 (surface and ribbon views).

**Figure S3.**
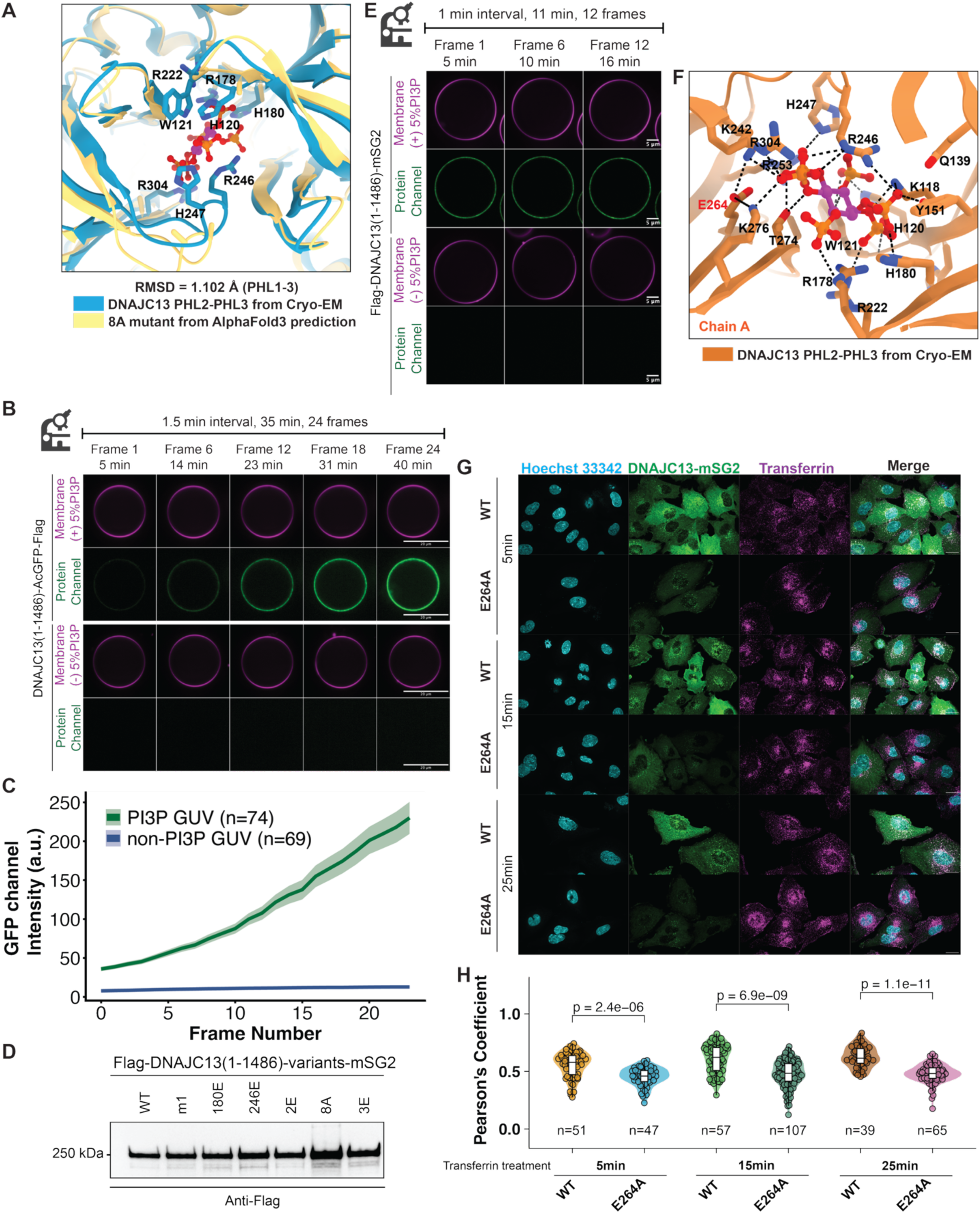
InsP6 binding is required for efficient recruitment of DNAJC13 to PI3P-containing membranes, related to Figure 3. (A) Structural comparison of the PHL2-PHL3 InsP_6_-binding pocket from the cryo-EM model (blue) and the 8A mutant predicted by AlphaFold3 (gold). Residues mutated in the 8A variant are shown as sticks at the corresponding positions in WT and 8A; the overall RMSD for PHL1-3 is indicated. (B) Time-lapse confocal imaging of DNAJC13(1-1486)-AcGFP-Flag incubated with GUVs containing 5% PI3P or lacking PI3P. DNAJC13 accumulates robustly and progressively on PI3P-containing membranes but not on PI3P-deficient GUVs. Magenta, fluorescent lipid membrane dye; green, DNAJC13(1-1486)-AcGFP-Flag. Representative frames from a 35-min imaging series (1.5-min interval, 24 frames total) are shown. Scale bars, 20 µm. (C) Quantification of protein binding to PI3P-containing (dark green) versus non-PI3P (light green) GUVs over time. GFP channel intensity at the membrane is plotted as mean ± SEM for the indicated number of GUVs (n). (D) Immunoblot of input Flag-DNAJC13(1-1486)-mSG2 variant proteins used in GUV binding assays, probed with anti-FLAG antibody, confirming comparable protein levels for all constructs. Note: the blot contains variant proteins that were not employed in GUV binding assays. (E) Time-lapse images of PI3P-containing GUVs incubated with DNAJC13(1-1486)-mSG2. Protein (green) rapidly and stably decorates the PI3P-positive membrane but not PI3P-deficient membrane (magenta). Images were acquired every 1 min for 11 min. (F) Close-up view of the InsP_6_-binding pocket in chain A of the cryo-EM structure, highlighting residues that coordinate InsP_6_. E264 forms hydrogen bonds with K276 and K242 in PHL3, positioning these lysines to engage InsP_6_ phosphates. (G) Confocal images of SUM159 cells expressing DNAJC13-mSG2 WT or the E264A mutant (green, see **Data S3B** for immunoblots) after 5, 15, or 25 min of fluorescent Transferrin uptake (magenta). Nuclei are labeled with Hoechst 33342 (cyan). Compared with WT, E264A shows reduced punctate localization and diminished overlap with Transferrin-positive endosomes. Scale bars, 20 µm. (H) Pearson’s correlation analysis of DNAJC13-Transferrin colocalization for WT and E264A at the indicated time points. Violin plots show the distribution of Pearson’s coefficients with median and interquartile ranges; n indicates the number of cells analyzed. p values (two-tailed t-test) are shown above the corresponding comparisons.

**Figure S4.**
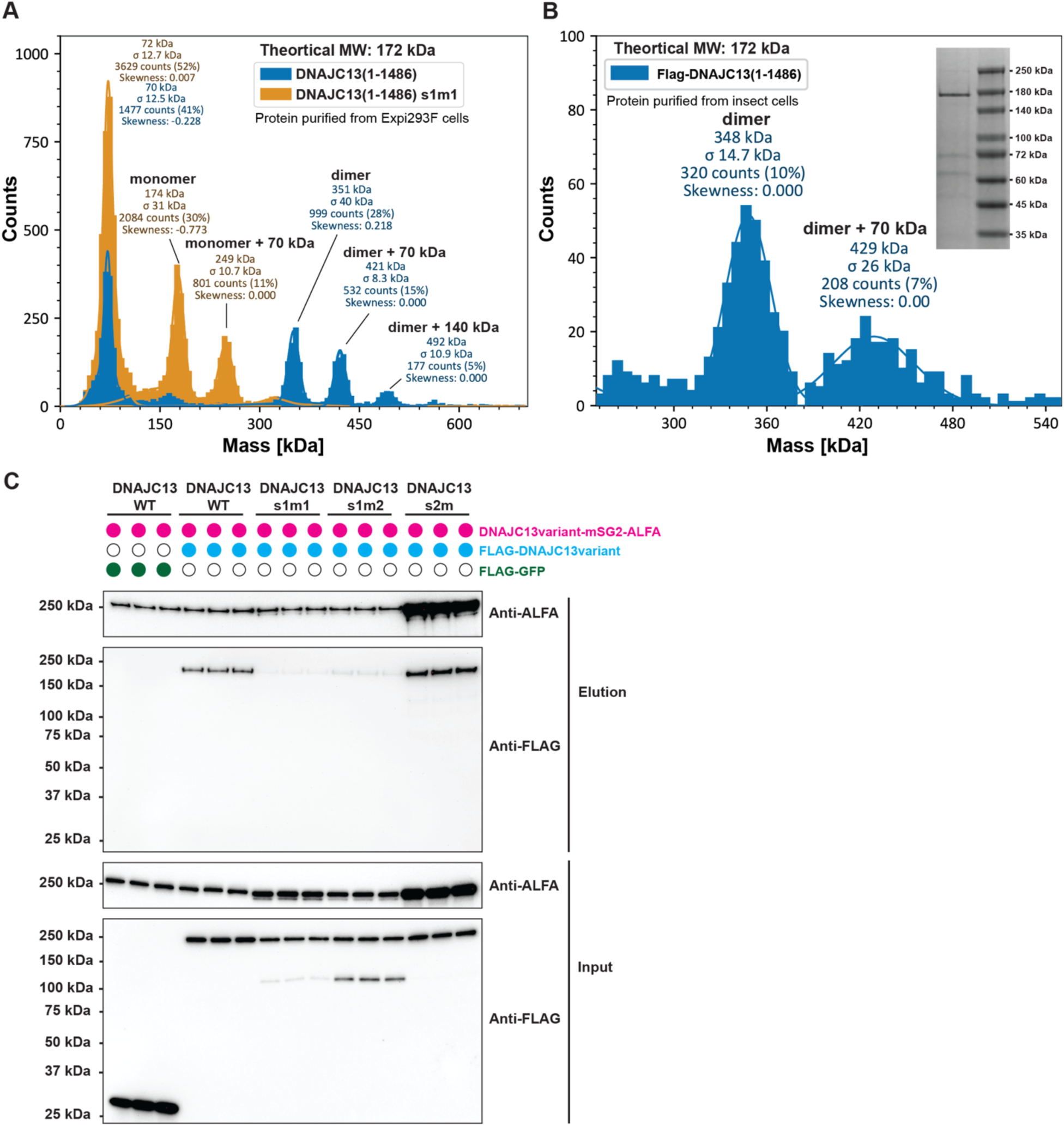
Mass photometry and co-immunoprecipitation demonstrate that DNAJC13 forms a stable homodimer, related to Figure 4. (A) Mass photometry of Flag-DNAJC13(1-1486) (orange) and the site-1 interface mutant s1m1 (blue) purified from Expi293F cells. DNAJC13(1-1486) (theoretical monomer mass 172 kDa) is observed as dimer, and dimer plus one or two 70-kDa binding partners. In contrast, s1m1 produces peaks corresponding almost exclusively to monomer and monomer plus a 70-kDa complex, indicating that s1m1 disrupted dimerization. (B) Mass photometry of Flag-DNAJC13(1-1486) purified from insect cells. A predominant peak at 348 kDa is consistent with a dimer, with a smaller population of dimer plus a 70-kDa binding partner. Inset, Coomassie-stained SDS-PAGE of the protein. (C) Pull-down assay testing dimer formation of DNAJC13 variants in DNAJC13-KO HEK293T cells. ALFA-tagged DNAJC13-mSG2 variants were co-expressed with FLAG-tagged DNAJC13 variants or FLAG-GFP (negative control, open circles) in HEK293T cells. Top panels: anti-ALFA and anti-FLAG immunoblot of proteins eluted from ALFA resin (Elution fraction). Bottom panels: corresponding input lysates. Equal volumes of input or eluate were loaded for all conditions.

**Figure S5.**
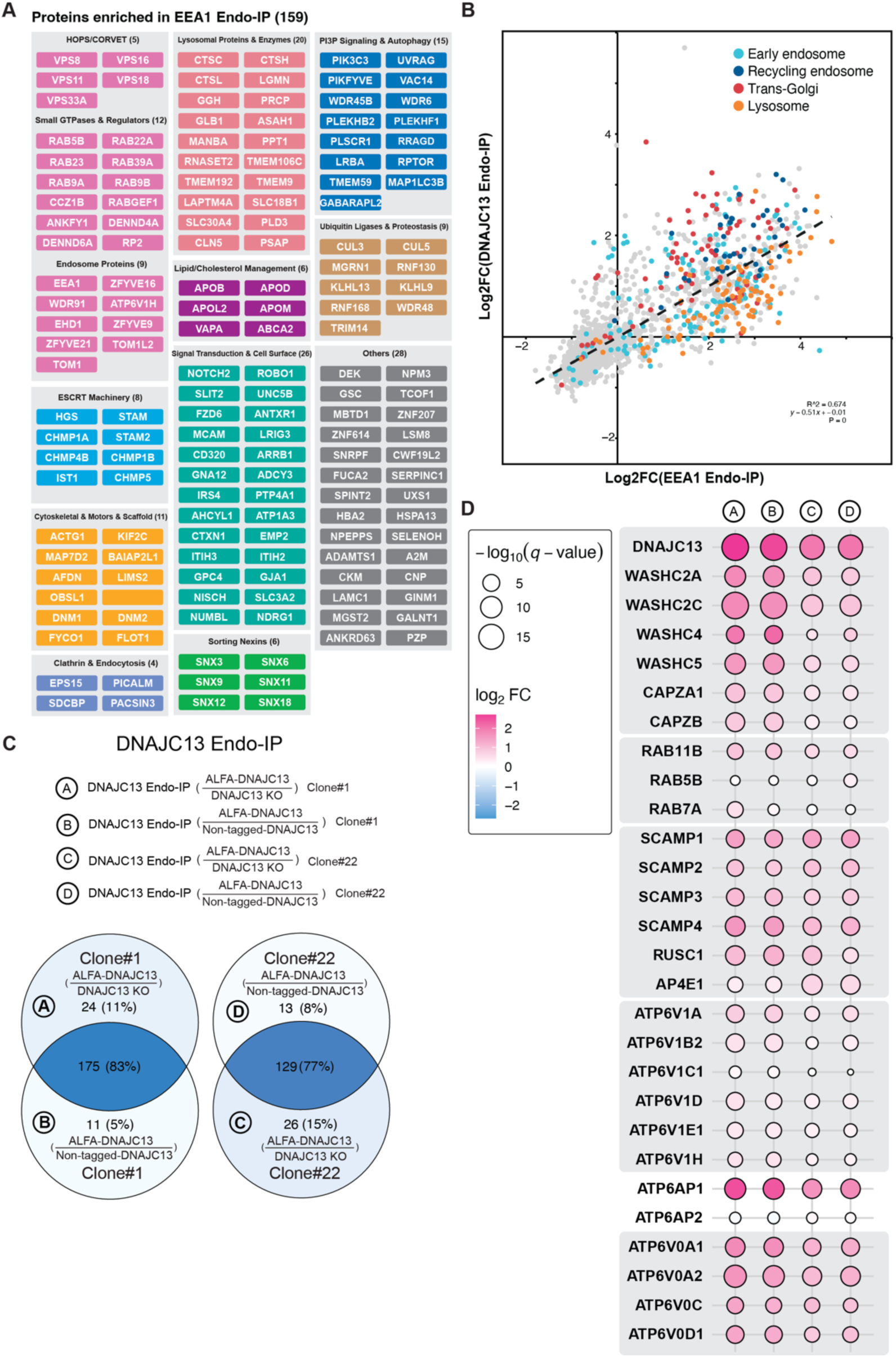
DNAJC13 Endo-IP preferentially enriches TGN and recycling-endosome components, related to Figure 5. (A) Functional annotation of proteins selectively enriched in EEA1 Endo-IP (FC > 1.5, q < 0.01) as compared with DNAJC13 Endo-IP. (B) Global comparison of protein enrichment between EEA1 Endo-IP and DNAJC13 Endo-IP. Each point represents a protein, plotted by its log_2_ fold change in the EEA1 Endo-IP (x-axis) versus the DNAJC13 Endo-IP (y-axis). Proteins annotated as early endosome, trans-Golgi, recycling endosome, or lysosome (as employed previously^45^) are highlighted in cyan, red, navy, and orange, respectively; all other proteins are shown in gray. The dashed line shows the linear regression across all proteins, with the regression equation, R^2^, and P value indicated. (C, D) Reproducibility across DNAJC13 Endo-IPs using either Clone #1 or #22 cells expressing ALFA-tagged DNAJC13 with or without expression of untagged DNAJC13, as outlined in Panel C. Panel D shows a bubble heat map comparing the relative enrichment of selected proteins in the various DNAJC13 Endo-IP samples from Panel C. Circle color indicates log₂ fold change and circle size represents -log₁₀(q-value). Proteins include WASH complex subunits and partners, V-ATPase subunits, recycling-endosome markers (RAB11B), and TGN-endosomal trafficking factors (RUSC1/2, AP-4 subunits).

**Figure S6.**
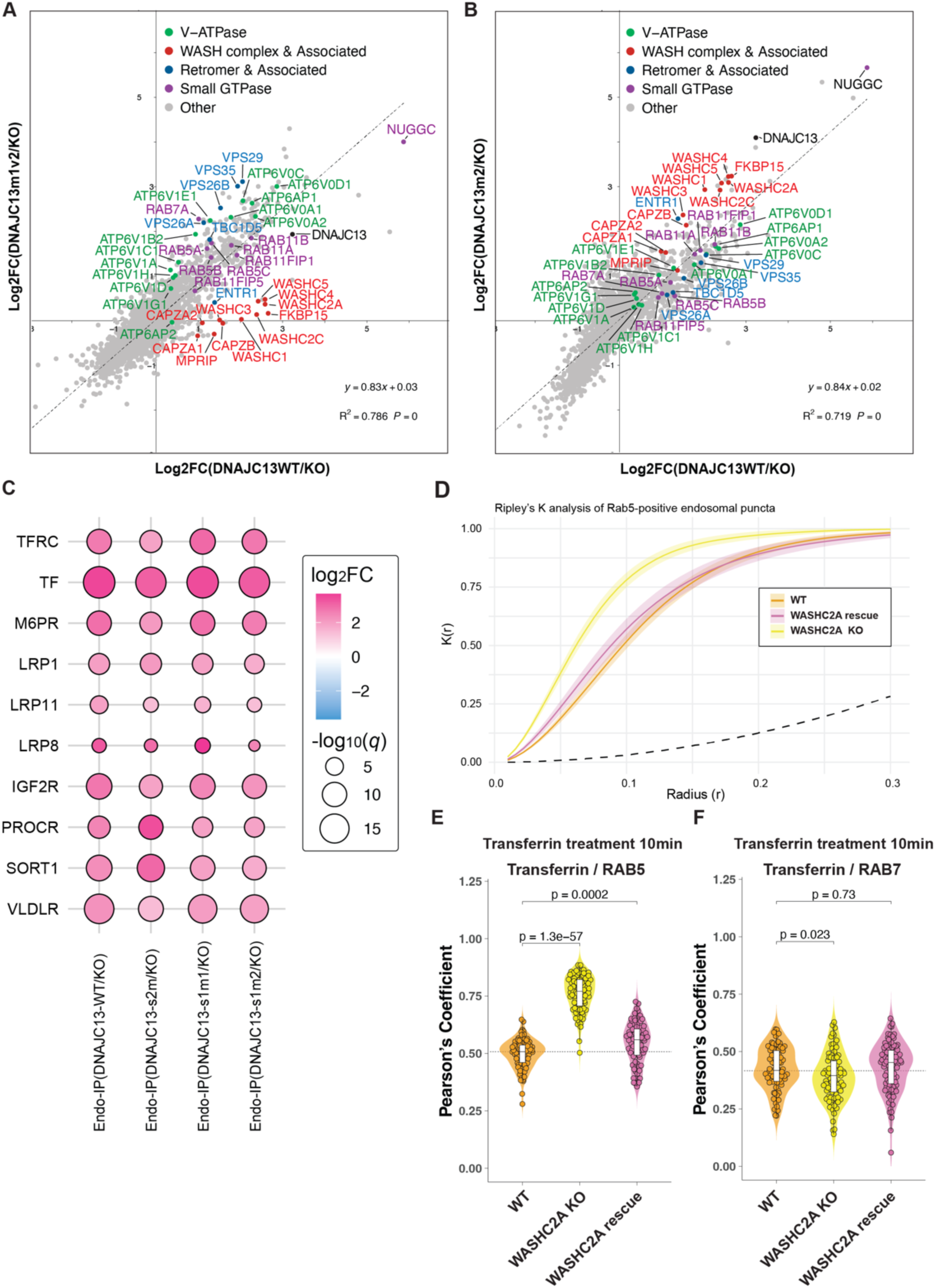
DNAJC13 dimerization is required for the WASH complex and V-ATPase assembly, related to Figure 6. (A, B) Comparison of DNAJC13 Endo-IP proteomes obtained with DNAJC13-WT/KO versus the dimerization-defective mutants DNAJC13-s1m1/KO (A) or DNAJC13-s1m2/KO (B). Scatter plots show log_2_ fold changes (WT/KO, x-axis) versus log_2_ fold changes (s1m1/KO or s1m2/KO, y-axis) for all detected proteins. V-ATPase subunits (green), WASH complex and associated proteins (red), retromer and associated proteins (blue), and small GTPases (purple) are highlighted. Linear-regression fits, equations, R^2^, and P values are shown. (C) Bubble heatmap summarizing enrichment of individual V-ATPase subunits in DNAJC13 Endo-IPs. Data are shown for two clonal lines (Clone#1 and Clone#22) expressing DNAJC13-WT, using non-tagged DNAJC13 or DNAJC13 KO cells as controls. Circle color indicates log_2_ fold change, and circle size reflects the -log_10_(q-value), illustrating the consistent enrichment of a greater number of V_0_ subunits compared to V_1_ subunits in DNAJC13-positive endosomes. (D) Ripley’s K(r) was computed from the centroid coordinates of Rab5 puncta within single-cell ROIs for the indicated groups. For each ROI, puncta coordinates were linearly normalized to a common global bounding box (0-1 in x and y), and K(r) was calculated across radii r = 0.01–0.3 in normalized units (area fixed to 1). Bold lines and shaded ribbons show the group mean ±95% confidence interval. The dashed line indicates the expectation under complete spatial randomness (CSR, K(r) = πr²), such that values above the dashed line reflect spatial clustering of Rab5-positive endosomal puncta relative to CSR. (E, F) Quantification of Transferrin colocalization with RAB5 (D) or RAB7 (E) in WT, WASHC2A-KO, and WASHC2A-rescued cells after 10-min Transferrin uptake. Violin plots show the distribution of Pearson’s coefficients for Transferrin/Rab5 or Transferrin/Rab7 colocalization; median and interquartile ranges are indicated. n denotes the number of cells analyzed, and p values (two-tailed t-test) are shown above the corresponding comparisons.

**Figure S7.**
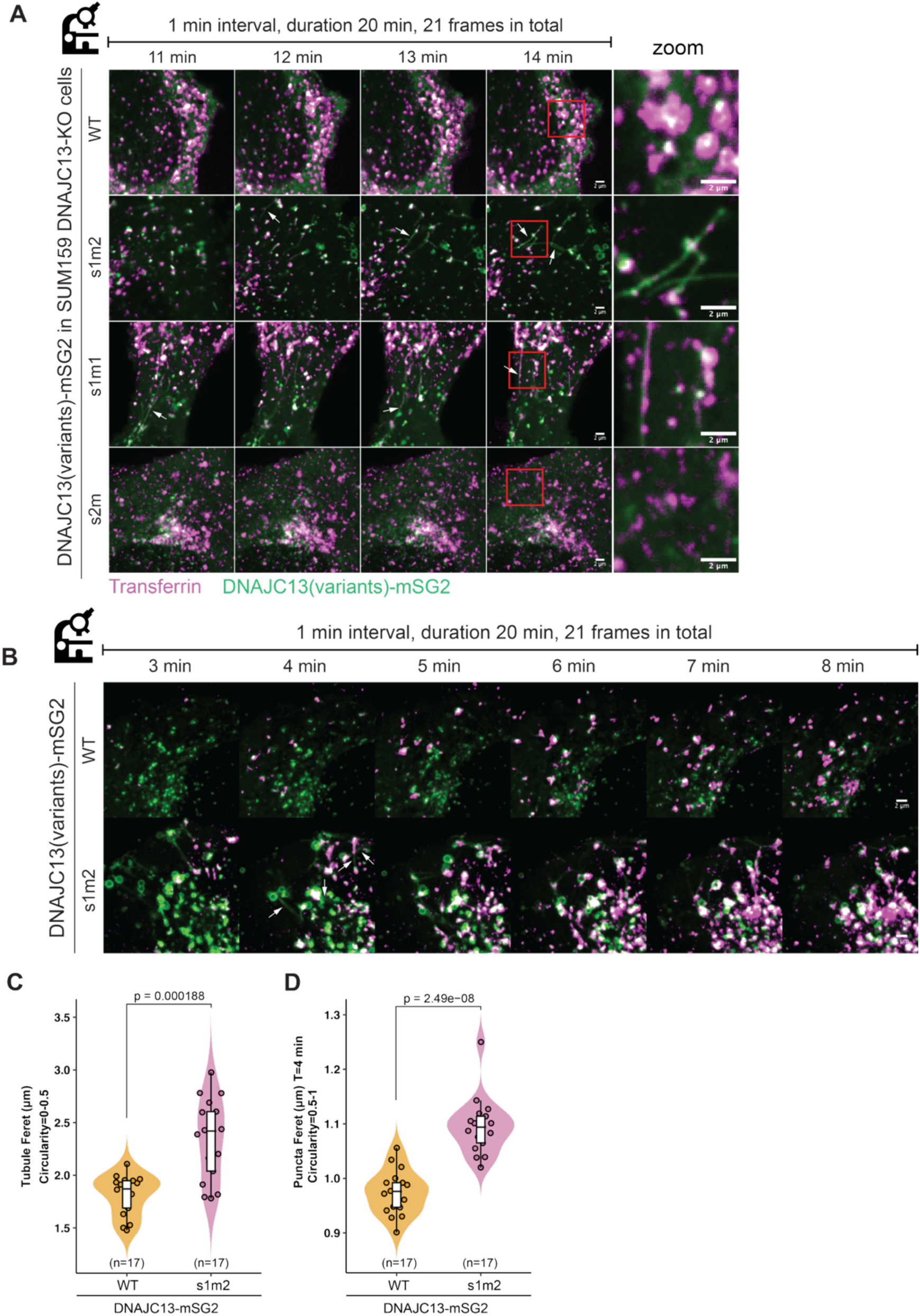
DNAJC13 dimerization controls endosomal tubulation. (A) Time-lapse imaging of DNAJC13^-/-^ SUM159 cells expressing DNAJC13-mSG2 variants (green) following fluorescent Transferrin uptake (magenta). DNAJC13-WT, the dimeric mutant (s2m), and the monomeric mutants (s1m1 and s1m2) were imaged at 1-min intervals for 20 min; representative fields from 11-14 min are shown, with zoomed views of the regions outlined at 14 min on the right. Dimeric DNAJC13 (WT and s2m) predominantly decorates punctate endosomes, whereas monomeric variants (s1m1 and s1m2) form branched, highly dynamic DNAJC13- and Transferrin-positive tubules that emanate from endosomes and fail to undergo efficient fission. The red box in time 14 indicates the region shown in the rightmost panels. Scale bars, 2 μm. Additional views from the same experiment are shown in Figure 7B. (B-D) Independent time-lapse experiment in DNAJC13^-/-^ SUM159 cells expressing DNAJC13-mSG2 variants (green) following fluorescent Transferrin uptake (magenta), imaged at 1-min intervals for 20 min. (Panel B) Representative fields for DNAJC13-WT-mSG2 and DNAJC13-s1m2-mSG2 between 3 and 8 min. (C, D) Quantification of DNAJC13(variants)-mSG2-positive structures at t = 4 min, separated by object circularity into tubular (circularity 0-0.5) and punctate (circularity 0.5-1) populations. (Panel C) Violin plot show the distribution of tubule length (Feret diameters) for tubular objects (circularity 0-0.5) in WT and s1m2 cells. (Panel D) Violin plot show the distribution of puncta ferret diameter for punctate objects (circularity 0.5-1) in WT and s1m2 cells. N indicates the number of cells analyzed per group. Groups were compared using a two-sided Wilcoxon rank-sum test (Mann-Whitney U test) on Feret values and the p-values are shown above the plots.

