## Supplementary material for "Structural Mechanisms of DNAJC13 Dimeric Assembly and InsP6 binding in Recycling Endosome Regulation": Table S1

**Table S1. Cryo-EM data collection, refinement and validation statistics**

|  | DNAJC13(1-1486)/InsP_6_ (PDB 37FE)  Consensus map (EMD-78131) |
| --- | --- |
| **Data collection and processing** |  |
| Magnification | 165,000 |
| Voltage (kV) | 300 |
| Electron exposure (e–/Å^2^) | 50.01 |
| Defocus range (μm) | -1.2 to -2.2 |
| Pixel size (Å) | 0.736 |
| Symmetry imposed | C1 |
| Initial particle images (no.) | 3,676,632 |
| Final particle images (no.) | 599,851 |
| Map resolution (Å)  FSC threshold | 2.4  0.143 |
| Map resolution range (Å) | 2.1-36.5 |
| **Refinement** |  |
| Initial model used (PDB code) | AlphaFold |
| Model resolution (Å)  FSC threshold | 2.7  0.5 |
| Model resolution range (Å) | 2.8-40.8 |
| Map sharpening *B* factor (Å^2^) | -77 |
| Model composition  Non-hydrogen atoms  Protein residues  Ligands | 19842  2614  IHP:2 |
| *B* factors (Å^2^)  Protein  Ligand | 93.84  119.96 |
| R.m.s. deviations  Bond lengths (Å)  Bond angles (°) | 0.010  1.191 |
| Validation  MolProbity score  Clashscore  Poor rotamers (%) | 1.59  9.28  0.00 |
| Ramachandran plot  Favored (%)  Allowed (%)  Disallowed (%) | 97.54  2.46  0.00 |

Composite map (EMD-78135) was generated from consensus map (EMD-78131) and 4 focused maps (EMD-78123, EMD-78128, EMD-78129, EMD-78130) was used for model building and figures in this study. Final model statistics for PDB 37FE were calculated against the consensus (EMD-78131).
