## Supplementary material for "Structural Mechanisms of DNAJC13 Dimeric Assembly and InsP6 binding in Recycling Endosome Regulation": Document S1_Data S1-S6

**Contents**

**Data S1.1. Additional details of the cryo-EM data-processing workflow in CryoSPARC, representative micrograph and 2D classifications, and map resolution analysis**

**Data S1.2. Quality of maps and models**

**Data S1.3. IWN repeats and ARM domain analysis**

**Data S1.4. GYF-like domains analysis and J-domain IP-MS analysis**

**Data S2.1. Structural and sequence determinants of $InsP_6$ and PI(3)P binding by DNAJC13 PH-like domains**

**Data S2.2. Generation and validation of DNAJC13 and WASHC2A knockout cell lines**

**Data S2.3. Colocalization analysis of DNAJC13 dimer-interface, and PH-like domain mutations with Transferrin**

**Data S2.4. Replicate cleavage assays used for quantification of $InsP_6$ release from DNAJC13 variants in Figure 2D**

**Data S2.5. Validation of $InsP_6$ standard**

**Data S3. Western blots to confirm expression levels of DNAJC13 variants in cells**

**Data S4. Validation of ALFA-tagged DNAJC13 and EEA1 cell lines for Endo-IPs**

**Data S5. WASHC2A loss disrupts early endosome distribution and promotes clustering of Transferrin-positive endosome**

**Data S6. DNAJC13 is required for proper Transferrin endocytosis and endosomal distribution in SUM159 cells**

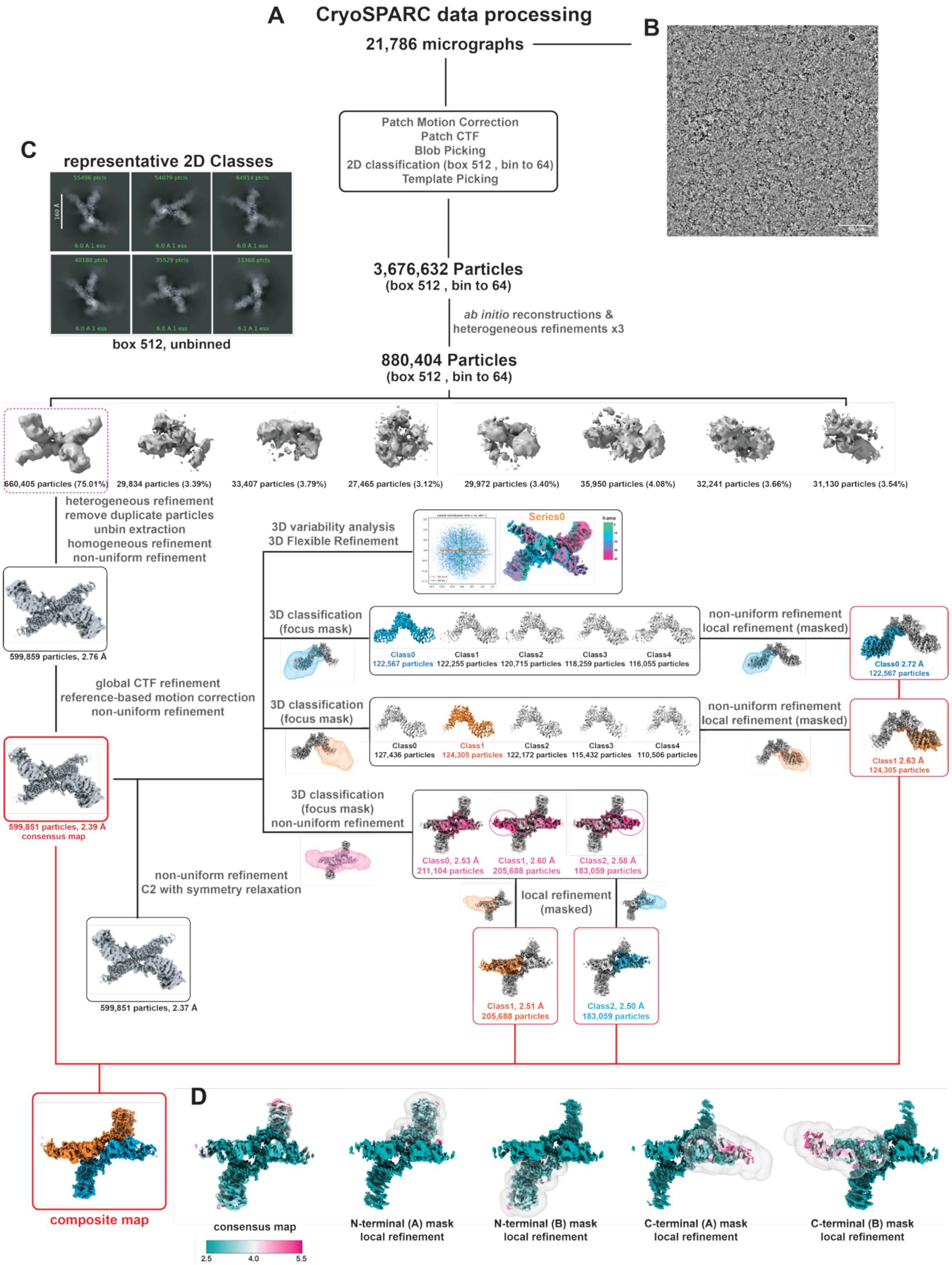


**Data S1.1. Additional details of the cryo-EM data-processing workflow in CryoSPARC, representative micrograph and 2D classifications, and map resolution analysis.**

(A) Cryo-EM data analysis of DNAJC13(1-2191) in CryoSPARC. Flow chart of the data processing strategy using CryoSPARC to obtain a 2.4 Å consensus reconstruction and a composite map for model building.

(B) Representative motion-corrected 10 Å lowpass filtered micrograph of vitrified DNAJC13(1-2191) on a R1.2/1.3 UltrAuFoil grid. Scale bar represents 50 nm.
(C) Representative Cryo-EM single-particle 2D classes, generated in CryoSPARC, revealing an elongated, highly asymmetric particle with multiple branched arms, giving an arch-shaped dimeric structure. The arms appear flexible and variably oriented around a dense central hub, consistent with a dynamic, multi-arm architecture. Scale bar represents 160 Å.
(D) The indicated cryo-EM maps colored by local resolution

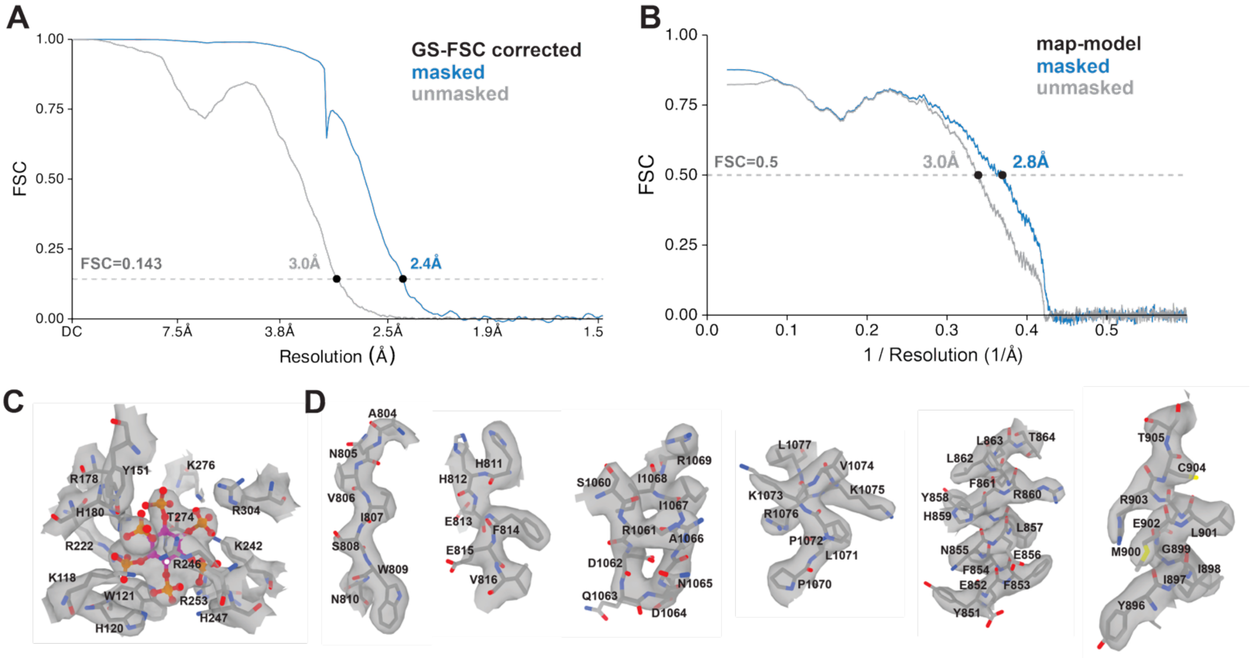


**Data S1.2. Quality of maps and models**

(A) Fourier shell correlation (FSC) vs. resolution (Å) curves for the indicated cryo-EM maps. Resolution was estimated at FSC=0.143 (grey dash line).

(B) Model vs. map FSC curves.

(C) Representative cryo-EM density in the composite map for the $InsP_6$-binding pocket residues within the PHL2-PHL3 region with corresponding atomic model.

(D) Representative cryo-EM density in the composite map for selected residues at the Site 1 and Site 2 dimerization interfaces with corresponding atomic model.

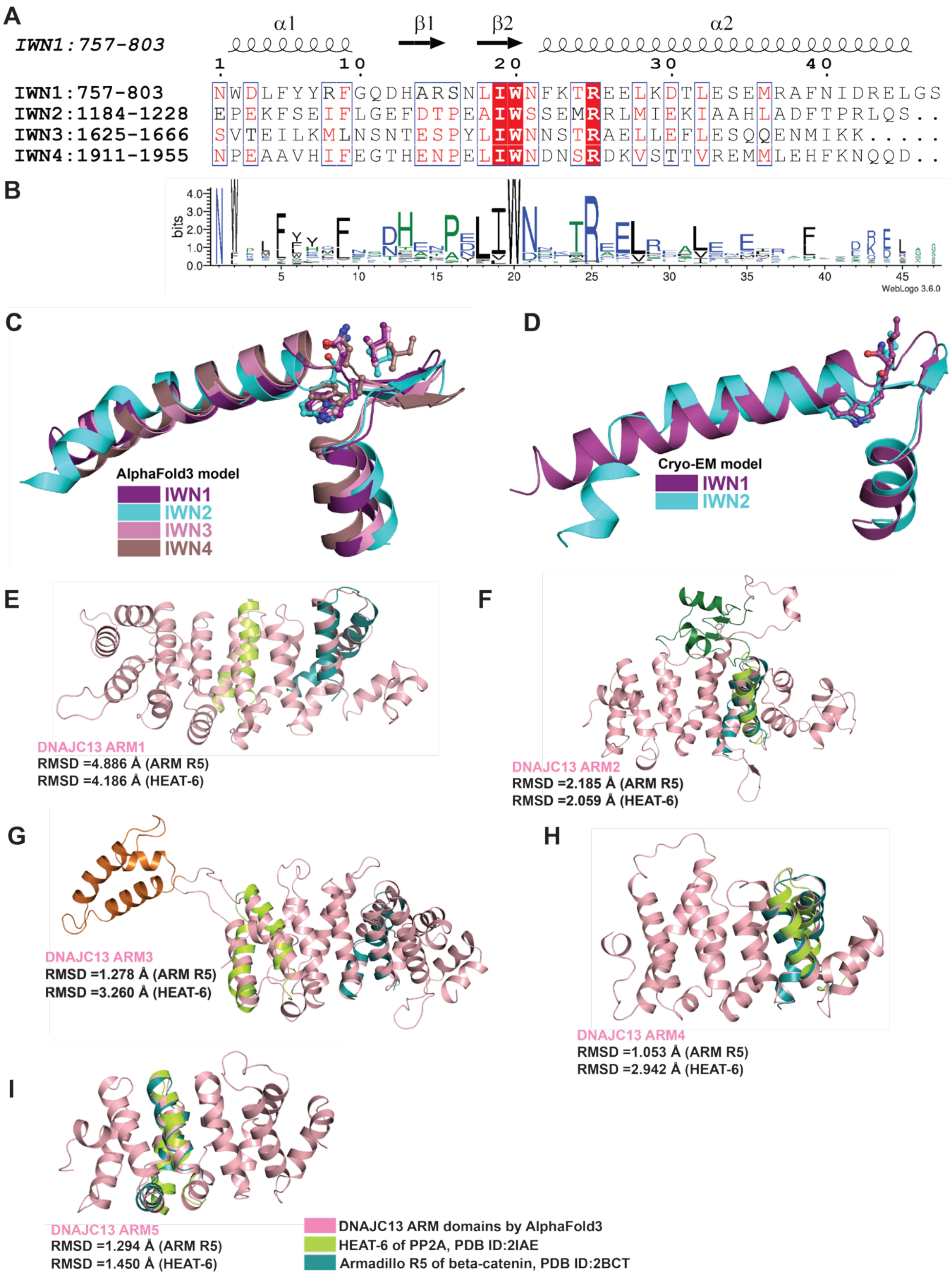


**Data S1.3. IWN repeats and ARM domain analysis**

(A) Multiple sequence alignment of the four IWN repeats. Cryo-EM secondary structure of IWN1 repeat is shown above. Strictly conserved residues are highlighted in red; conserved positions are boxed in blue.

(B) Sequence conservation of the indicated DNAJC13 IWN1 repeat. Position-specific amino-acid conservation is represented as a sequence logo generated with WebLogo 3.6.0 from the DeepMSA2 multiple sequence alignment server.

(C) Superposition of IWN1-4 from the AlphaFold3 DNAJC13 full-length model, colored as indicated. Side chains of strictly conserved residues I-W-[N/S] are shown as sticks.

(D) Superposition of the cryo-EM model of IWN1 and IWN2 colored as indicated. Side chains of strictly conserved residues I-W-[N/S] are shown as sticks.

(E-I) Structural comparison of DNAJC13 ARM1-5 domains with canonical ARM and HEAT repeat unit. AlphaFold3 models of individual DNAJC13 ARM domains (pink) were superposed with HEAT-6 of PP2A (green; PDB 2IAE) and the armadillo repeat R5 of β-catenin (cyan; PDB 2BCT). Panels show ARM1 (E), ARM2 (F), ARM3 (G), ARM4 (H), and ARM5 (I). Root-mean-square deviation (RMSD) values for each superposition are indicated in each panel for comparison with armadillo repeat R5 of β-catenin and HEAT-6 of PP2A.

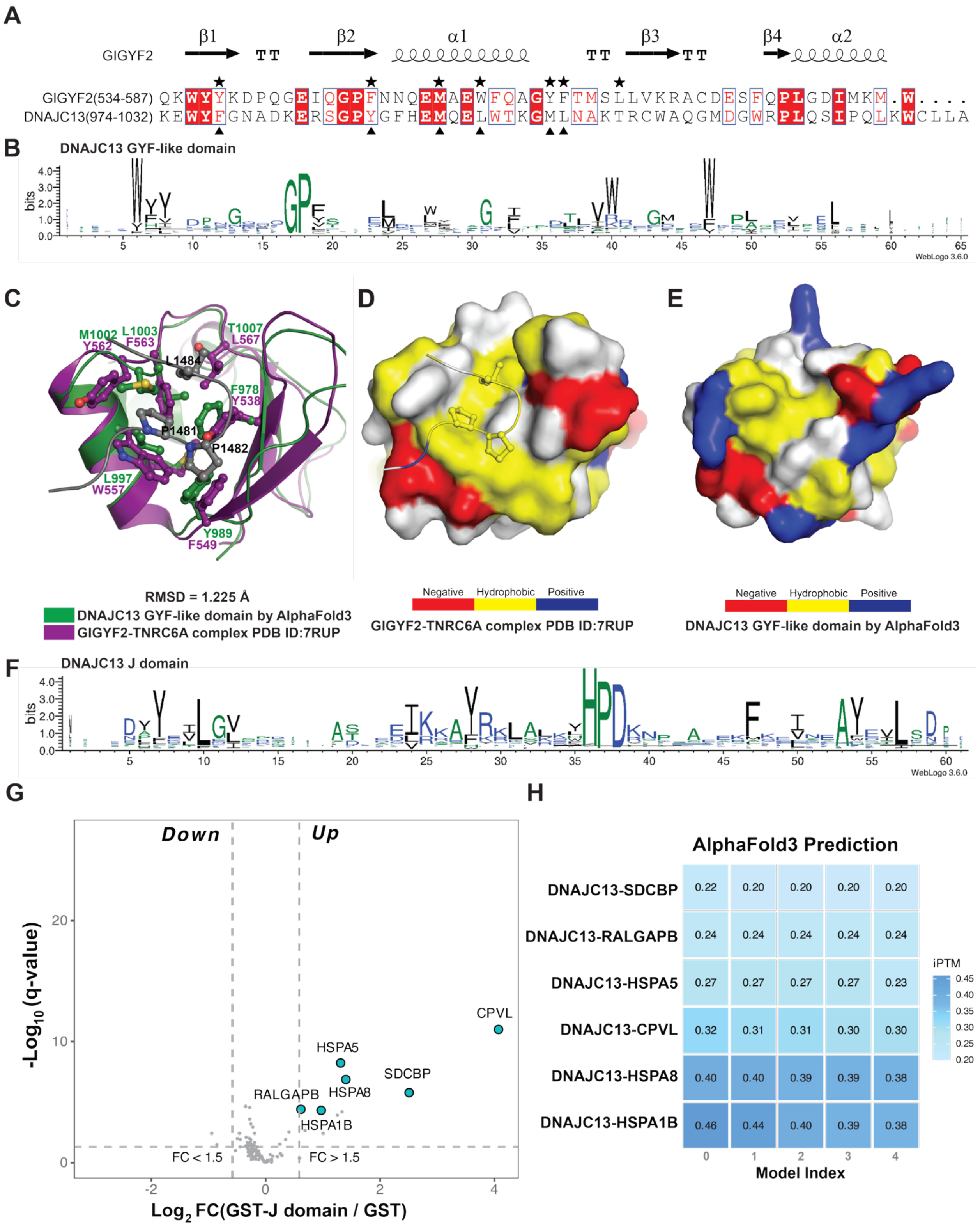


**Data S1.4. GYF-like domains analysis and J-domain IP-MS analysis**

(A) Structure-based sequence alignment of the GYF domain of GIGYF2 with the AlphaFold3-predicted GYF-like domain of DNAJC13. Secondary-structure elements derived from the GIGYF2 GYF domain are shown above. The canonical GYF motif has the consensus sequence GP[YF]xxxx[MV]xxWxxx[GN]YF.

Residues in GIGYF2 that form the hydrophobic groove for the PPGL motif of TNRC6A are indicated by black stars, and DNAJC13 residues that form a similar hydrophobic groove are indicated by black triangles.
(B) Sequence conservation of the indicated DNAJC13 GYF-like domain. Position-specific amino-acid conservation is represented as a sequence logo generated with WebLogo 3.6.0 from the DeepMSA2 multiple sequence alignment server.
(C) Structural superposition of the AlphaFold3 model of the DNAJC13 GYF-like domain (green) with the GIGYF2-TNRC6A complex (purple and grey; PDB 7RUP). Side chains forming the peptide-binding groove are shown as sticks; the RMSD is indicated.
(D, E) Combined surface and ribbon-stick representations showing the hydrophobic binding interface between the GIGYF2 GYF domain and the TNRC6A PPGL motif (Panel D), and the corresponding surface of the DNAJC13 GYF-like domain (Panel E). Surfaces are colored by amino-acid type: hydrophobic residues in yellow, positively charged residues in blue, negatively charged residues in red, and uncharged polar residues in white, illustrating a potential hydrophobic pocket in the DNAJC13 GYF-like domain that could accommodate hydrophobic motifs.
(F) Sequence conservation of the indicated J domain of DNAJC13. Position-specific amino-acid conservation is represented as a sequence logo generated with WebLogo 3.6.0 from the DeepMSA2 multiple sequence alignment server.
(G) Volcano plot of GST pulldown followed by quantitative mass spectrometry using the GST-J domain as bait. $Log_2$ FC (GST-J domain/GST) is plotted against $-log_{10}$(q-value); dashed lines indicate thresholds for fold change (FC > 1.5) and statistical significance (q < 0.05), and selected enriched interactors are labeled.
(H) AlphaFold3 prediction confidence (shown as iPTM values) for complexes between DNAJC13 and the binding partners identified in (G).

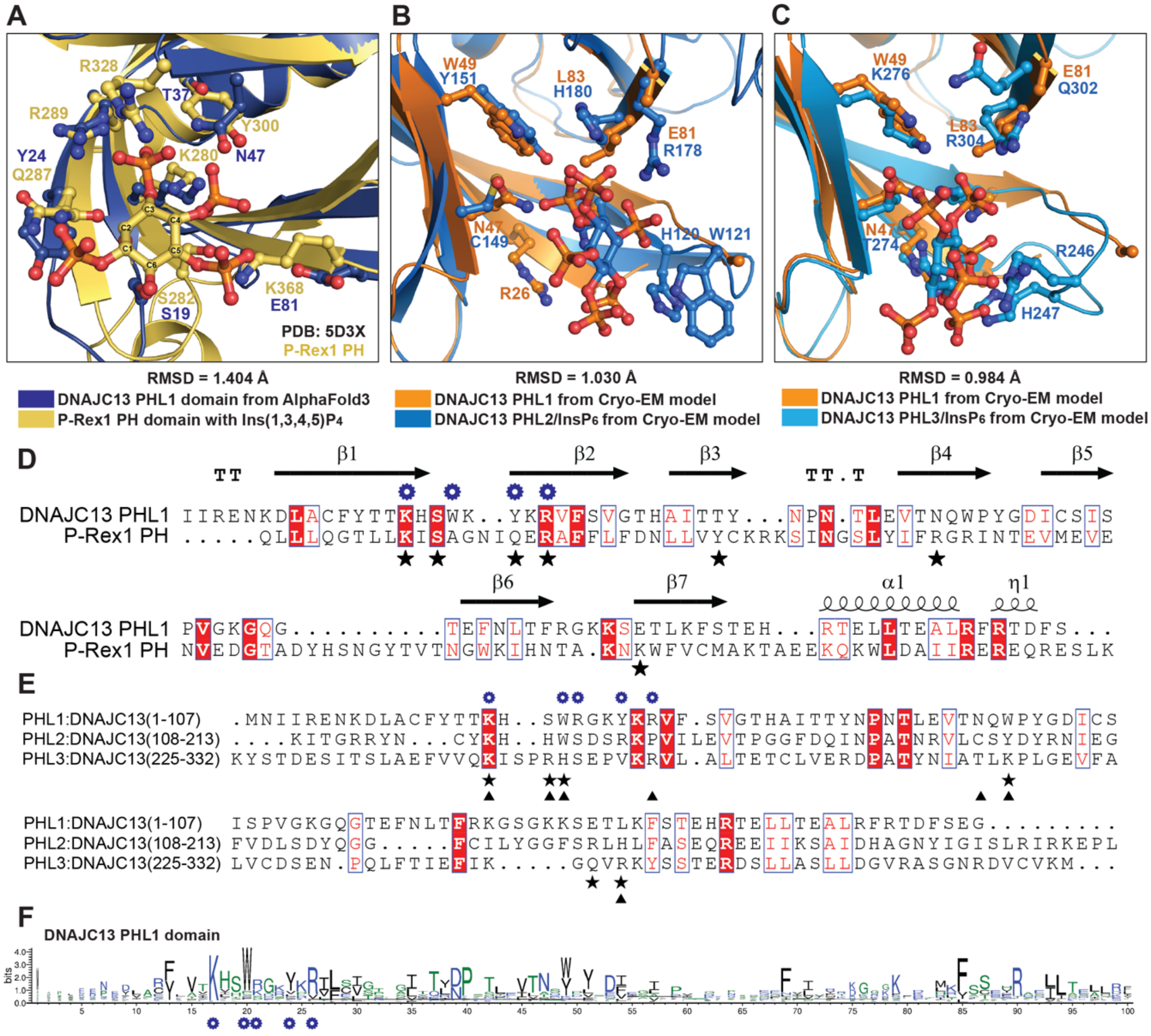


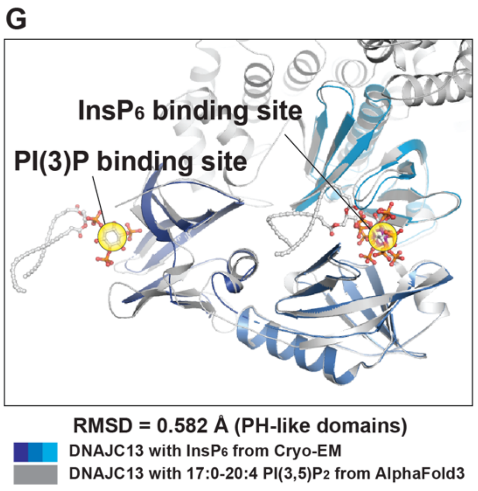


**H**

| $IP_4$ phosphate position | P-Rex1 PH binding-site residues | DNAJC13 PHL1 residues at equivalent positions |
|---|---|---|
| C1 | Q287 | Y24 |
| C3 | K280, R289, R328 | K17, R26, N47 |
| C4 | K280, S282, Y300 | K17, S19, T37 |
| C5 | K368 | E81 |

**I**

| $InsP_6$ phosphate position | DNAJC13 binding-site residues |
|---|---|
| C1 | R178, T274, K276 |
| C2 | Y151, R178, H180, R246 |
| C3 | K118, R178, R222 |
| C4 | K118, H120, W121, R246, H247 |
| C5 | K242, H247, R253 |
| C6 | K242, T274, K276, R304 |

**Data S2.1. Structural and sequence determinants of $InsP_6$ and PI(3)P binding by DNAJC13 PH-like domains**

(A) Superposition of the DNAJC13 N-terminal PH-like domain1 (PHL1) predicted by AlphaFold3 (blue) with the PH domain of P-Rex1 bound to Ins(1,3,4,5)$P_4$ (yellow; PDB 5D3X). Side chains that coordinate the inositol phosphate in P-Rex1 (yellow labels) and the corresponding residues in DNAJC13 (blue labels) are shown as sticks, together with the bound Ins(1,3,4,5)$P_4$ ligand (yellow carbon atoms). The RMSD between the two domains is 1.404 Å.

(B) Superposition of the cryo-EM structures of DNAJC13 PHL1 domain (orange) and DNAJC13 PHL2-$InsP_6$ domain (marine). Conserved basic and polar residues implicated in inositol phosphate binding are shown as sticks and labeled in the corresponding colors. The RMSD between PHL1 and PHL2 is 1.030 Å.

(C) Superposition of the cryo-EM structures of DNAJC13 PHL1 domain (orange) and DNAJC13 PHL3-$InsP_6$ domain (sky blue). Conserved basic and polar residues implicated in inositol phosphate binding are shown as sticks and labeled in the corresponding colors. The RMSD between PHL1 and PHL3 is 0.984 Å.

(D) Structure-based sequence alignment of the DNAJC13 PHL1 domain with the P-Rex1 PH domain, with PHL1 secondary-structure elements indicated above the alignment. Residues important for DNAJC13 binding to PI(3)P are marked with blue gears, and residues required for P-Rex1 PH binding to Ins(1,3,4,5)$P_4$ are marked with black stars. The PI(3)P-binding residues in PHL1 were inferred from the DNAJC13/17:0-20:4 PI(3,5)$P_2$ AlphaFold3 model and its structural alignment with the P-Rex1 PH/Ins(1,3,4,5)$P_4$ complex shown in (A).

(E) Sequence alignment of PHL1-3 from DNAJC13. Residues involved in PI(3)P binding in PHL1 are marked with blue gears, residues involved in $InsP_6$ binding in PHL2 are marked with black stars, and residues involved in $InsP_6$ binding in PHL3 are marked with black triangles.

(F) Sequence conservation of the indicated PHL1 domain of DNAJC13. Position-specific amino-acid conservation is represented as a sequence logo generated with WebLogo 3.6.0 from the DeepMSA2 multiple sequence alignment server.

(G) Overall superposition of the three DNAJC13 PH-like domains bound to $InsP_6$ in the cryo-EM structure (dark blue, marine, sky blue) with the AlphaFold3 model of the PH-like domains bound to two copies of PI(3,5)$P_2$ (grey), showing the relative locations of the predicted PI(3)P and observed $InsP_6$ binding sites; RMSD value between PH-like domains is 0.582 Å.

(H) Table summarizing the correspondence between Ins(1,3,4,5)$P_4$-binding residues in the P-Rex1 PH domain and the equivalent positions in DNAJC13 PHL1, illustrating the molecular mechanism by which DNAJC13 specifically binds PI(3)P.

(I) Table summarizing the DNAJC13 residues in the PHL domains that contact each phosphate group of $InsP_6$.

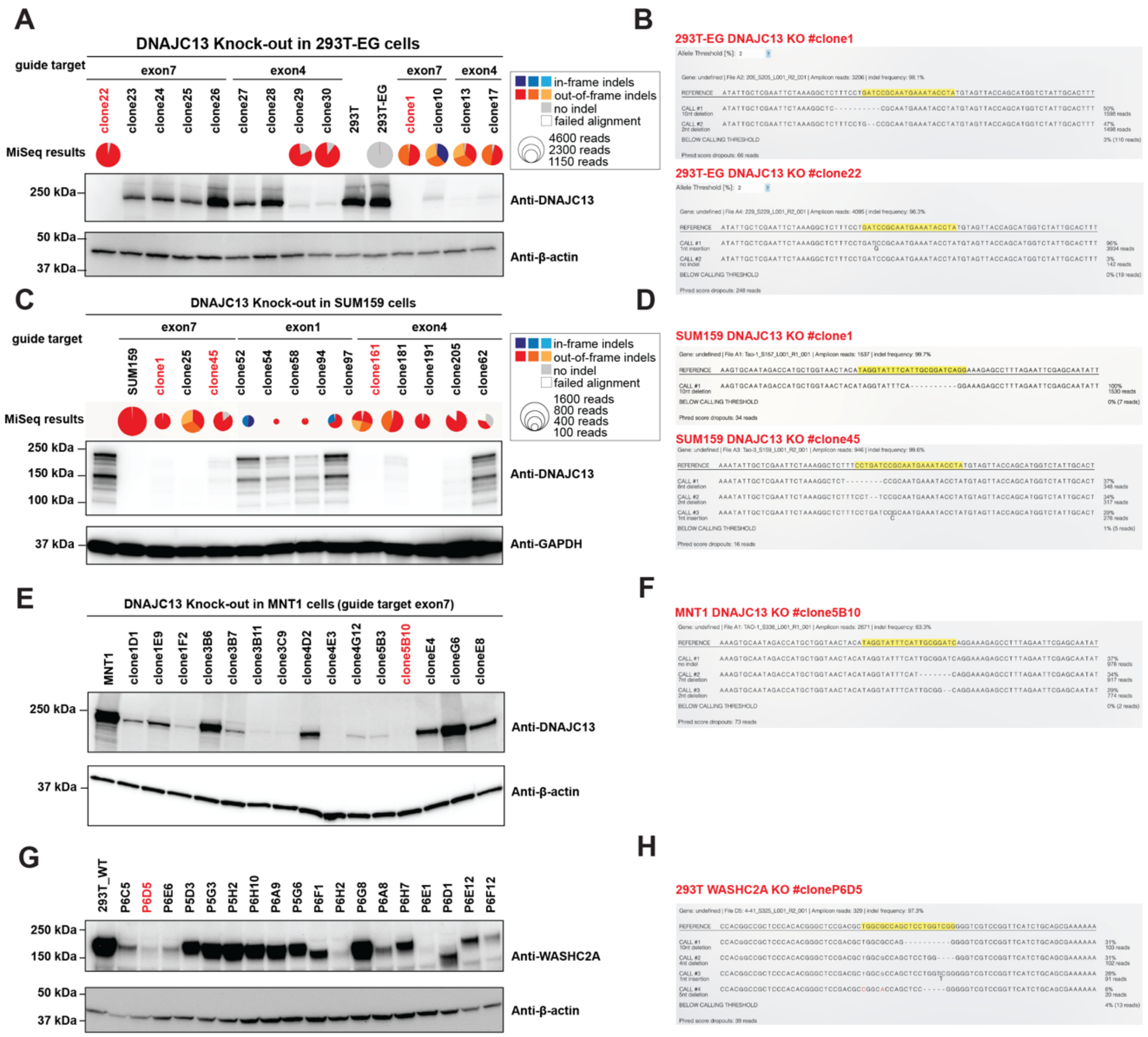
A
DNAJC13 Knock-out in 293T-EG cells
guide target
exon7
exon4
exon7
exon4
clone22
clone23
clone24
clone25
clone26
clone27
clone28
clone29
clone30
293T
293T-EG
clone1
clone10
clone13
clone17
MiSeq results
in-frame indels
out-of-frame indels
no indel
failed alignment
4600 reads
2300 reads
1150 reads
250 kDa
Anti-DNAJC13
50 kDa
37 kDa
Anti-β-actin
B
293T-EG DNAJC13 KO #clone1
293T-EG DNAJC13 KO #clone22
C
DNAJC13 Knock-out in SUM159 cells
guide target
exon7
exon1
exon4
SUM159
clone1
clone25
clone45
clone52
clone54
clone58
clone94
clone97
clone161
clone181
clone191
clone205
clone62
MiSeq results
in-frame indels
out-of-frame indels
no indel
failed alignment
1600 reads
800 reads
400 reads
100 reads
250 kDa
150 kDa
100 kDa
Anti-DNAJC13
37 kDa
Anti-GAPDH
D
SUM159 DNAJC13 KO #clone1
SUM159 DNAJC13 KO #clone45
E
DNAJC13 Knock-out in MNT1 cells (guide target exon7)
MNT1
clone1D1
clone1E9
clone1F2
clone3B6
clone3B7
clone3B11
clone3C9
clone4D2
clone4E3
clone4G12
clone5B3
clone5B10
cloneE4
cloneG6
cloneE8
250 kDa
Anti-DNAJC13
37 kDa
Anti-β-actin
F
MNT1 DNAJC13 KO #clone5B10
G
293T_WT
P6C5
P6D5
P6E6
P5D3
P5G3
P5H2
P6H10
P6A9
P5G6
P6F1
P6H2
P6G8
P6A8
P6H7
P6E1
P6D1
P6E12
P6F12
250 kDa
150 kDa
Anti-WASHC2A
50 kDa
37 kDa
Anti-β-actin
H
293T WASHC2A KO #cloneP6D5

**Data S2.2. Generation and validation of DNAJC13 and WASHC2A knockout cell lines**
(A-B) Immunoblot analysis of DNAJC13 expression in $293T^{EG}$ cells edited with the indicated CRISPR guides targeting exon 7 or exon 4 of DNAJC13. Parental $293T^{EG}$ cells and individual clones are shown. Pie charts summarize MiSeq results for each clone (fractions of in-frame indels, out-of-frame indels, no indel, and failed alignment; circle size indicates total read number). Anti-β-actin serves as a loading control (A). Representative MiSeq alignments for indicated $293T^{EG}$ DNAJC13 knockout clones showing the edited sequences is presented in (B).
(C-D) Immunoblot analysis of DNAJC13 expression in SUM159 cells edited with the indicated CRISPR guides. Parental SUM159 cells and individual clones are shown with corresponding MiSeq summary pie charts. GAPDH serves as a loading control (C). Representative MiSeq alignments for SUM159 DNAJC13 knockout clones showing the edited sequences are shown in (D).
(E-F) Immunoblot analysis of DNAJC13 expression in MNT1 cells targeted with a CRISPR guide against exon 7. Parental MNT1 cells and individual clones are shown; anti-β-actin serves as a loading control (E). Representative MiSeq alignment for an MNT1 DNAJC13 knockout clone showing the edited sequences is presented in (F).
(G-H) Immunoblot analysis of WASHC2A expression in HEK293T cells edited with the indicated CRISPR guides. Parental HEK293T cells and individual clones are shown; anti-β-actin serves as a loading control (G). Representative MiSeq alignment for a HEK293T $WASHC2A^{-/-}$ clone showing the edited sequences is presented in (H).

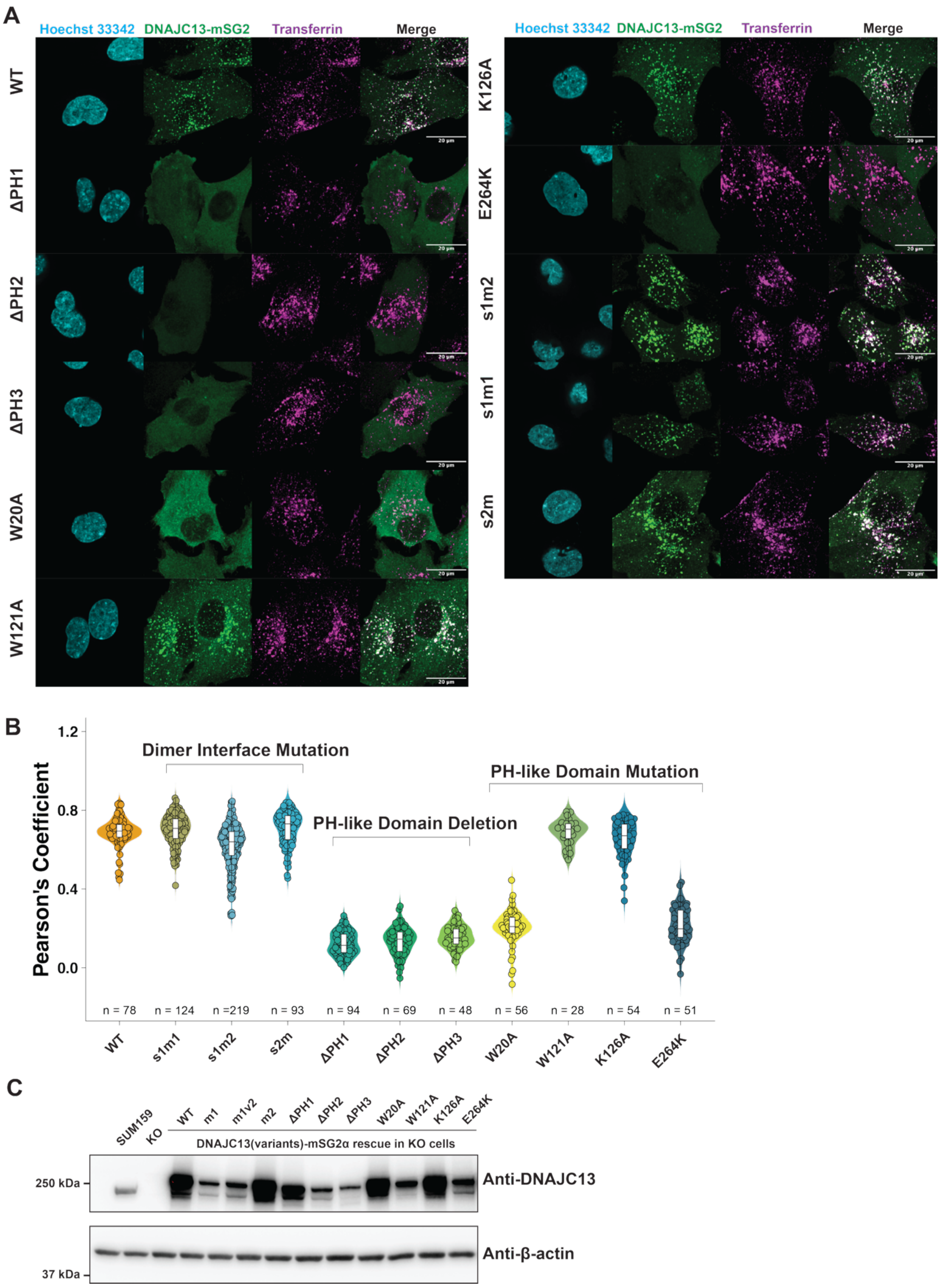


**Data S2.3. Colocalization analysis of DNAJC13 dimer-interface, and PH-like domain mutations with Transferrin**

(A) Representative confocal images of DNAJC13$^{-/-}$ SUM159 cells expressing mSG2-tagged DNAJC13 WT or the indicated mutants. Cells were subjected to a 10-min fluorescent transferrin uptake (magenta). DNAJC13-mSG2 variants are shown in green and nuclei are stained with Hoechst 33342 (cyan); merged images show transferrin distribution relative to each DNAJC13 variant. Mutations include dimer-interface mutants (s1m1, s1m2, s2m), PH-like domain deletions (ΔPH1–ΔPH3), PH-like domains point mutants (W20A, W121A, K126A, E264K), and Parkinson's-disease-linked mutations (V722L, N855S, R1266Q, T1895M). Scale bars, 20 μm.

(B) Quantification of transferrin colocalization with DNAJC13-mSG2 variants, expressed as Pearson's correlation coefficients for the indicated variants. Each point represents a single cell; violin plots show the distribution of values, and n indicates the number of cells analyzed per condition.

(C) DNAJC13 expression in SUM159 DNAJC13-KO cells, WT-rescued cells, and 8A-mutant rescued cells. DNAJC13 was detected with anti-DNAJC13 antibody; β-actin serves as a loading control. Uncropped raw images are shown in **Document S2**.

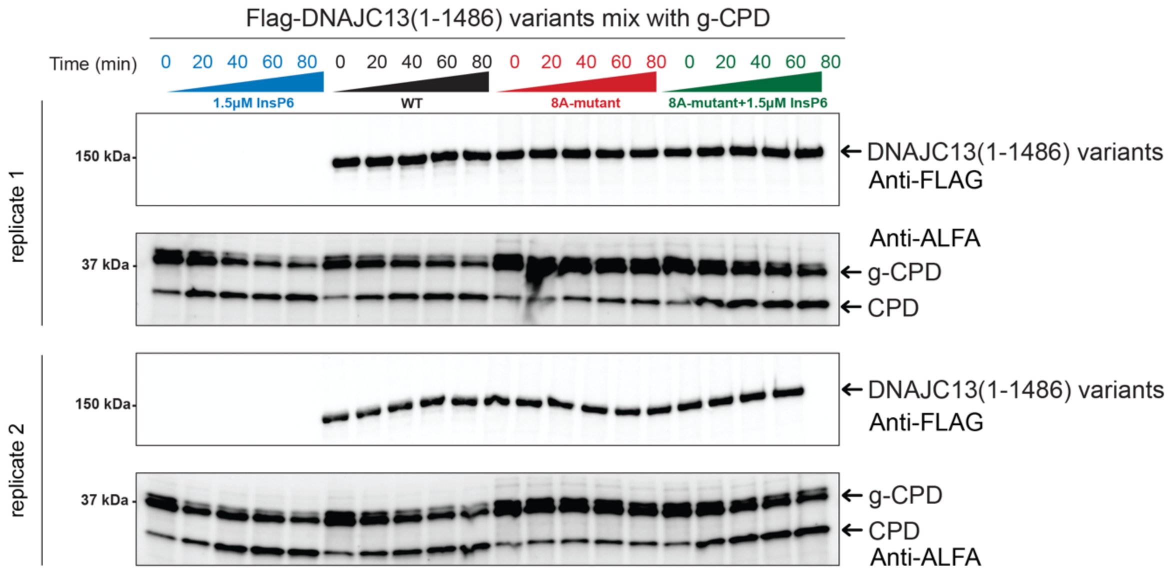


**Data S2.4. Replicate cleavage assays used for quantification of $InsP_6$ release from DNAJC13 variants in Figure 2D**

Time-course cleavage assays of g-CPD activated by $InsP_6$ released from Flag-DNAJC13(1-1486) variants. Reactions were performed with 1.5 μM $InsP_6$ alone, WT DNAJC13(1-1486), the 8A mutant, or the 8A mutant supplemented with 1.5 μM $InsP_6$, and samples were collected at the indicated times (0-80 min). Three independent technical replicates are shown; replicate 3 is presented as the representative blot in **Figure 2C**. Upper panels, immunoblots for DNAJC13(1-1486) variants probed with anti-Flag antibody; lower panels, immunoblots for g-CPD and CPD probed with Alexa-647-conjugated ALFA nanobody. Corresponding band intensities were quantified, and the results are shown in **Figure 2D**.

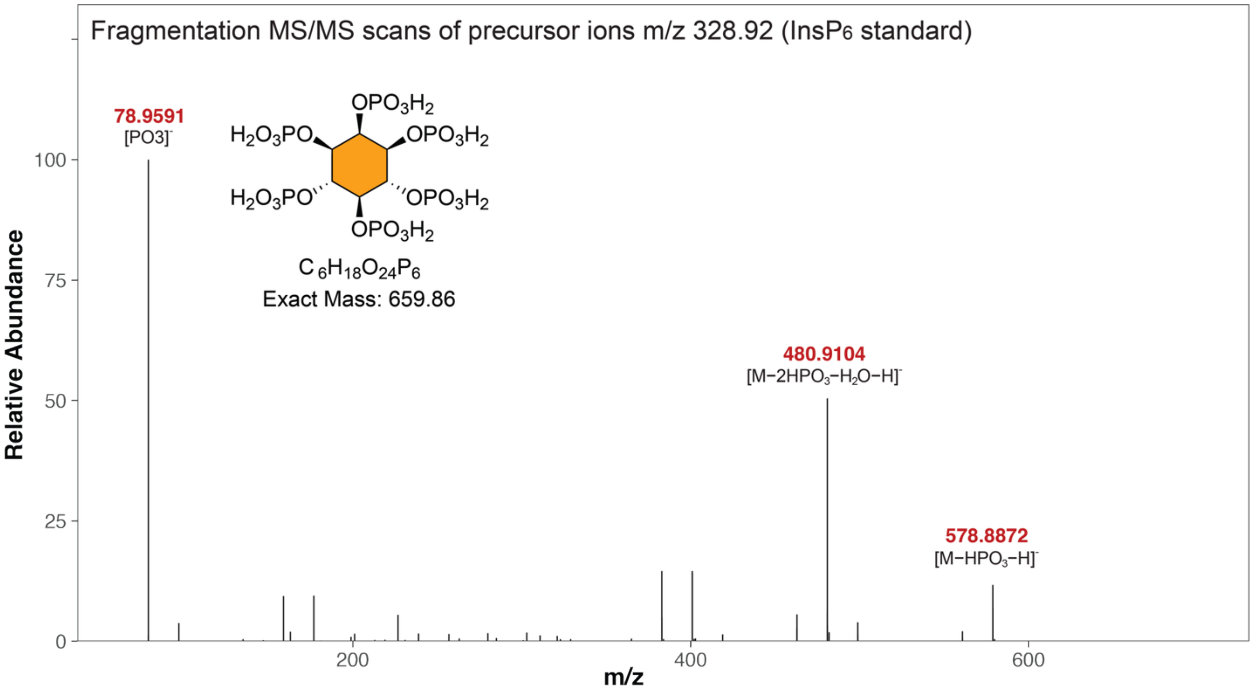


**Data S2.5. Validation of $InsP_6$ standard**

Fragmentation MS/MS spectrum of the $InsP_6$ standard (precursor ion m/z 328.92). Major fragment ions corresponding to successive phosphate losses are indicated, and the chemical structure and exact mass (659.86 Da) of $InsP_6$ are shown.

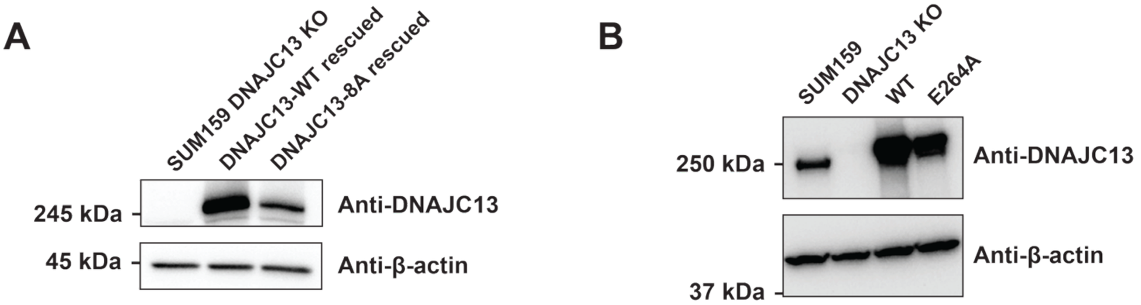


**Data S3. Western blots to confirm expression levels of DNAJC13 variants in cells**

(A) Immunoblot analysis of DNAJC13-KO SUM159 cells rescued with the indicated mSG2-tagged DNAJC13 variants. DNAJC13 was detected with anti-DNAJC13 antibody; β-actin serves as a loading control. Related to confocal imaging in **Figure 3C**.

(B) DNAJC13 expression in parental SUM159 cells, DNAJC13-KO cells, DNAJC13-WT rescued cells, and DNAJC13-E264A rescued cells. DNAJC13 was detected with anti-DNAJC13 antibody; β-actin serves as a loading control. Related to confocal imaging in **Figure S3G**.

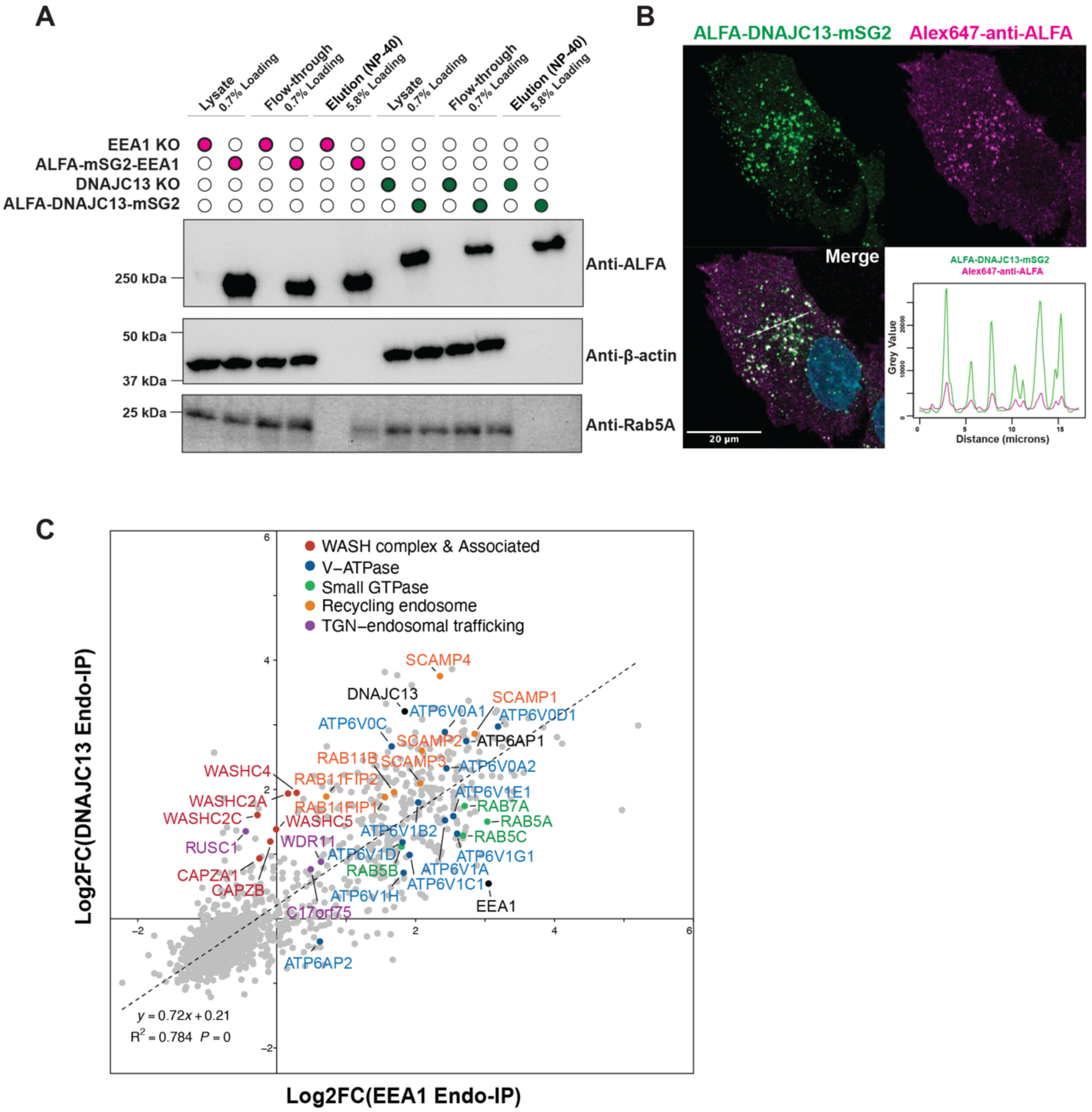


**Data S4. Validation of ALFA-tagged DNAJC13 and EEA1 cell lines for Endo-IPs**

(A) Immunoblot analysis of EEA1 Endo-IP and DNAJC13 Endo-IP. Lysate, flow-through, and NP-40 elution fractions (0.7% and 5.8% loadings as indicated) were probed with anti-ALFA to detect the ALFA-tagged baits, anti-β-actin as a loading control, and anti-Rab5 to verify enrichment of early endosomes.

(B) Confocal images of DNAJC13-KO SUM159 cells expressing ALFA-DNAJC13-mSG2 (green) stained with Alexa647-conjugated anti-ALFA nanobody (magenta). The merged image and corresponding line-scan intensity profile demonstrate strong colocalization between the fluorescent protein tag and the ALFA epitope signal, confirming the specificity of the ALFA nanobody for ALFA tag. Scale bar, 20 µm.

(C) Correlation scatter plot of log2 FC (rescue/KO) for proteins detected in EEA1 Endo-IP (x-axis) versus DNAJC13 Endo-IP (y-axis). Data were compiled from two independent sets of EEA1 Endo-IP and DNAJC13 Endo-IP experiments, each quantified in separate TMTpro multiplexes. Selected proteins are colored and annotated by functional class (WASH complex and associated proteins, V-ATPase, small GTPases, recycling-endosome markers, and TGN-endosomal trafficking factors). Dashed line indicates the linear regression fit.

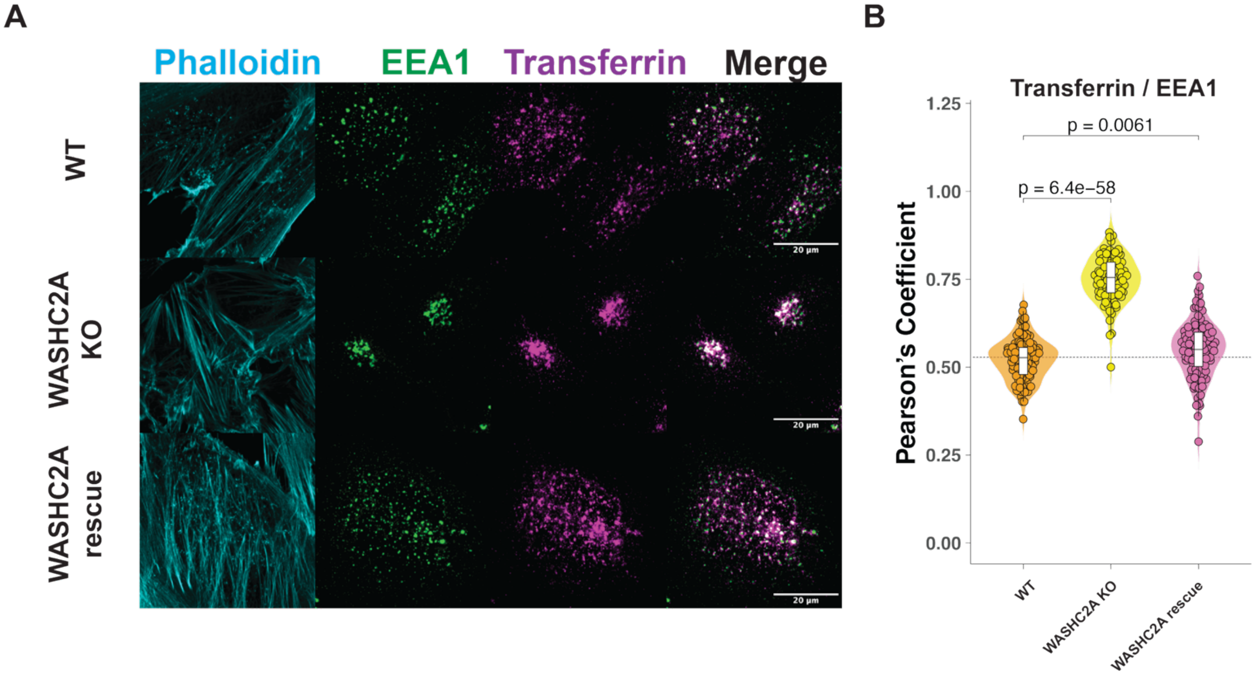


**Data S5. WASHC2A loss disrupts early endosome distribution and promotes clustering of Transferrin-positive endosome**

(A) Confocal images of SUM159 WT cells, WASHC2A-KO and rescued cells after a 10-min uptake of fluorescent Transferrin (magenta). Early endosomes are labeled with EEA1 (green, as indicated), and F-actin is labeled with phalloidin (cyan). Merged images show Transferrin distribution relative to endosomal markers for each genotype. Scale bars, 20 µm. Images for WT, WASHC2A-KO, and WASHC2A-rescued cells co-stained with RAB5 and RAB7 are selected to shown in **Figure 6C and 6D**.

(B) Quantification of Transferrin colocalization with EEA1, expressed as Pearson's correlation coefficients for the indicated genotypes. Each point represents one cell; violin plots show the distribution of values. Pearson's correlation coefficients for WT, WASHC2A-KO, and WASHC2A rescued cells of Transferrin with RAB5 or RAB7 are selected to shown in **Figure S6D** and **S6E**.

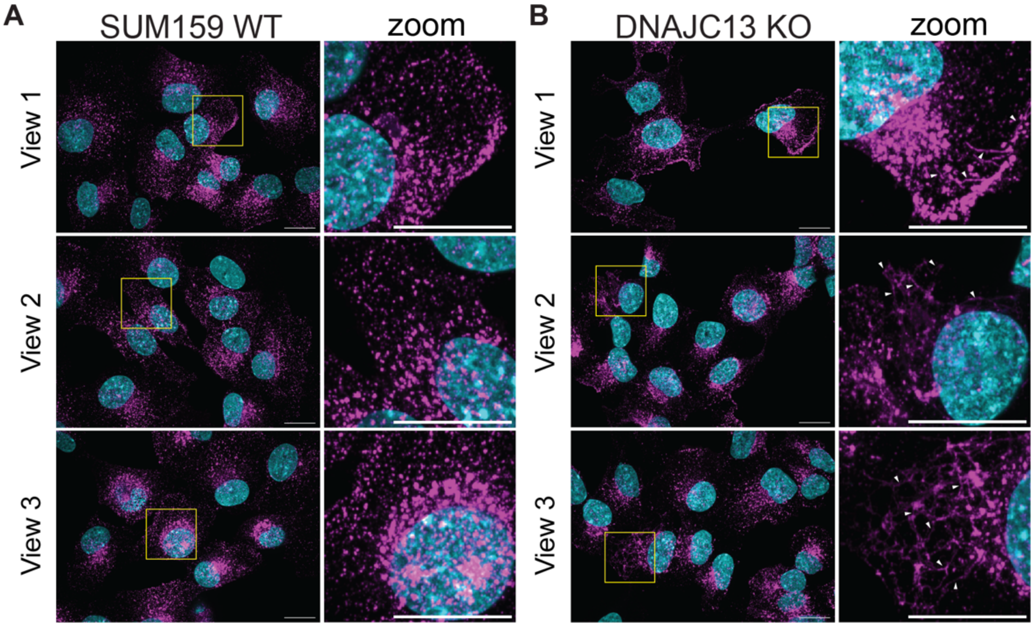


**Data S6. DNAJC13 is required for proper Transferrin endocytosis and endosomal distribution in SUM159 cells**

(A, B) Representative confocal images of SUM159 WT (Panel A) and DNAJC13-KO (Panel B) cells following a 10-min uptake of fluorescent Transferrin (magenta). Nuclei are stained with Hoechst 33342 (cyan). Three independent fields of view are shown for each condition. Yellow boxes indicate zoomed regions. Scale bars, 20 µm. The zoomed image in View 3 was selected as representative view of the phenotype shown in **Figure 7A**.
