## Supplementary material for "Structural Mechanisms of DNAJC13 Dimeric Assembly and InsP6 binding in Recycling Endosome Regulation": Document S2

Raw images of western blots included in this study. Page 1/6

Raw data for Figure 2C and Data S2.4

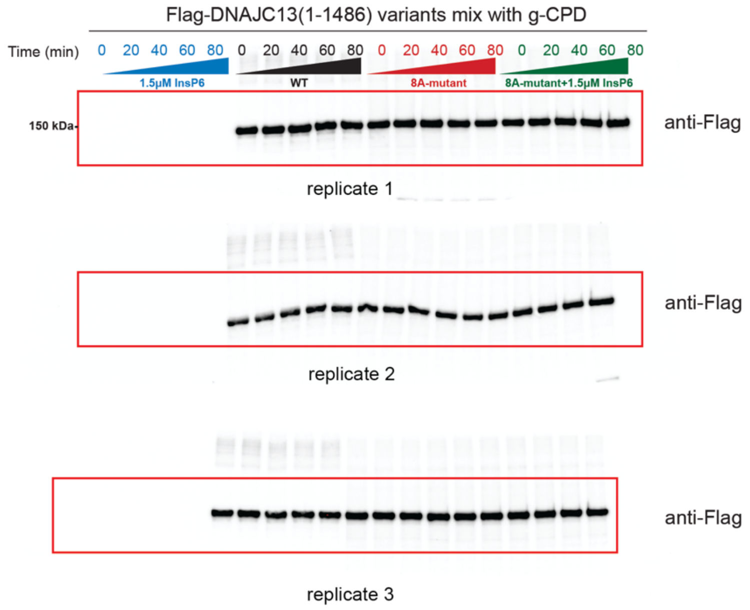


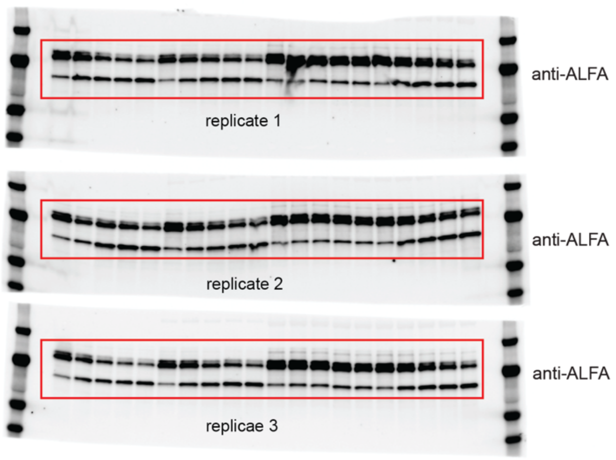

Raw images of western blots included in this study. Page 2/6

Raw data for Figure 7E

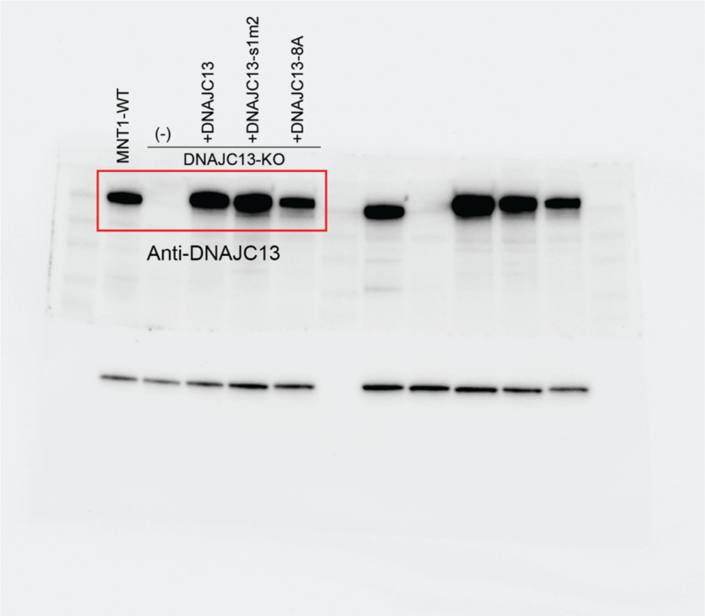


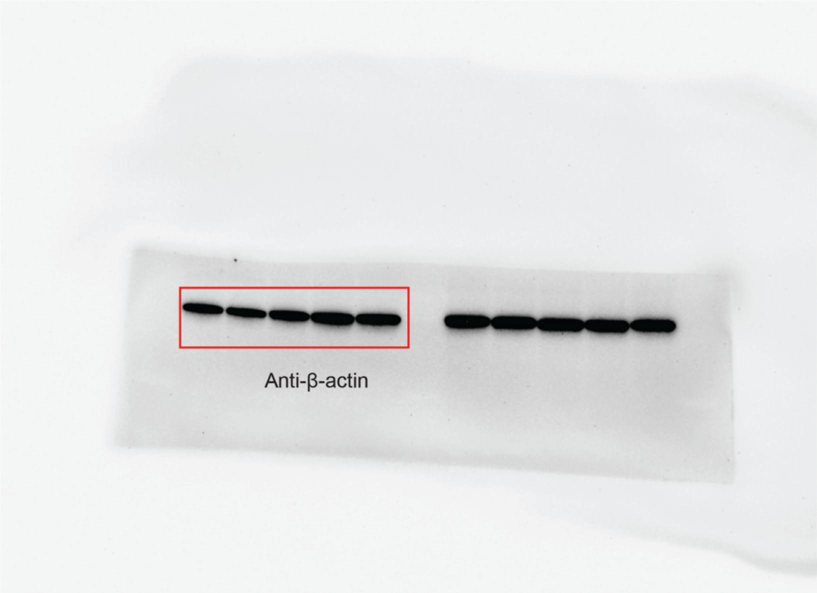

Raw images of western blots included in this study. Page 3/6

Raw data for Data S2.3C

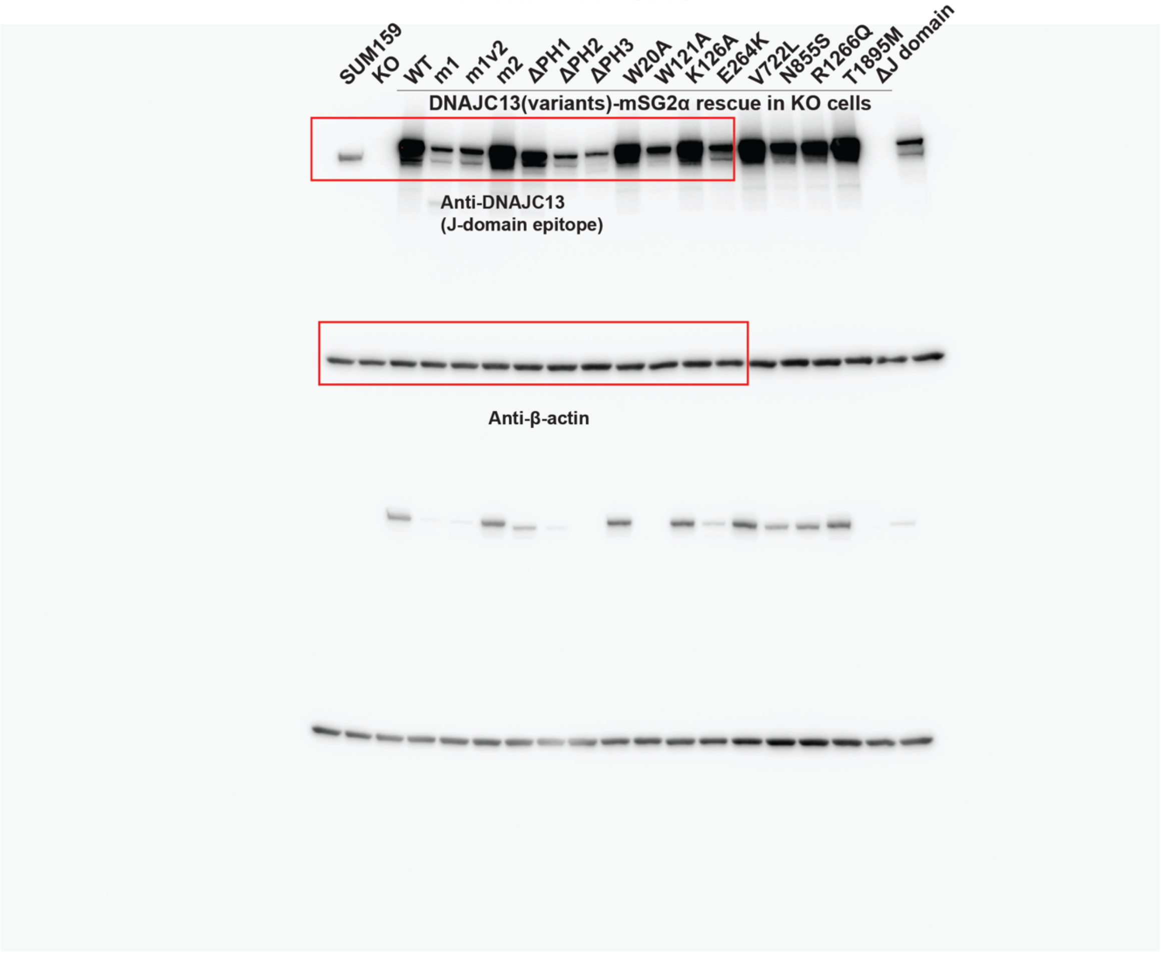

Raw images of western blots included in this study. Page 4/6

Raw data for Data S3A

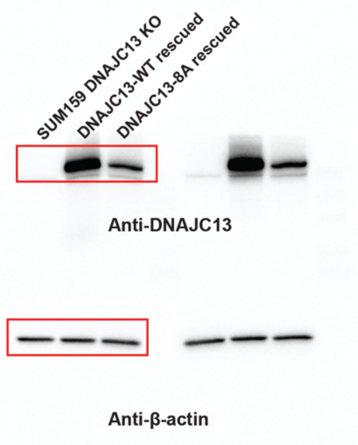


Raw data for Data S3B

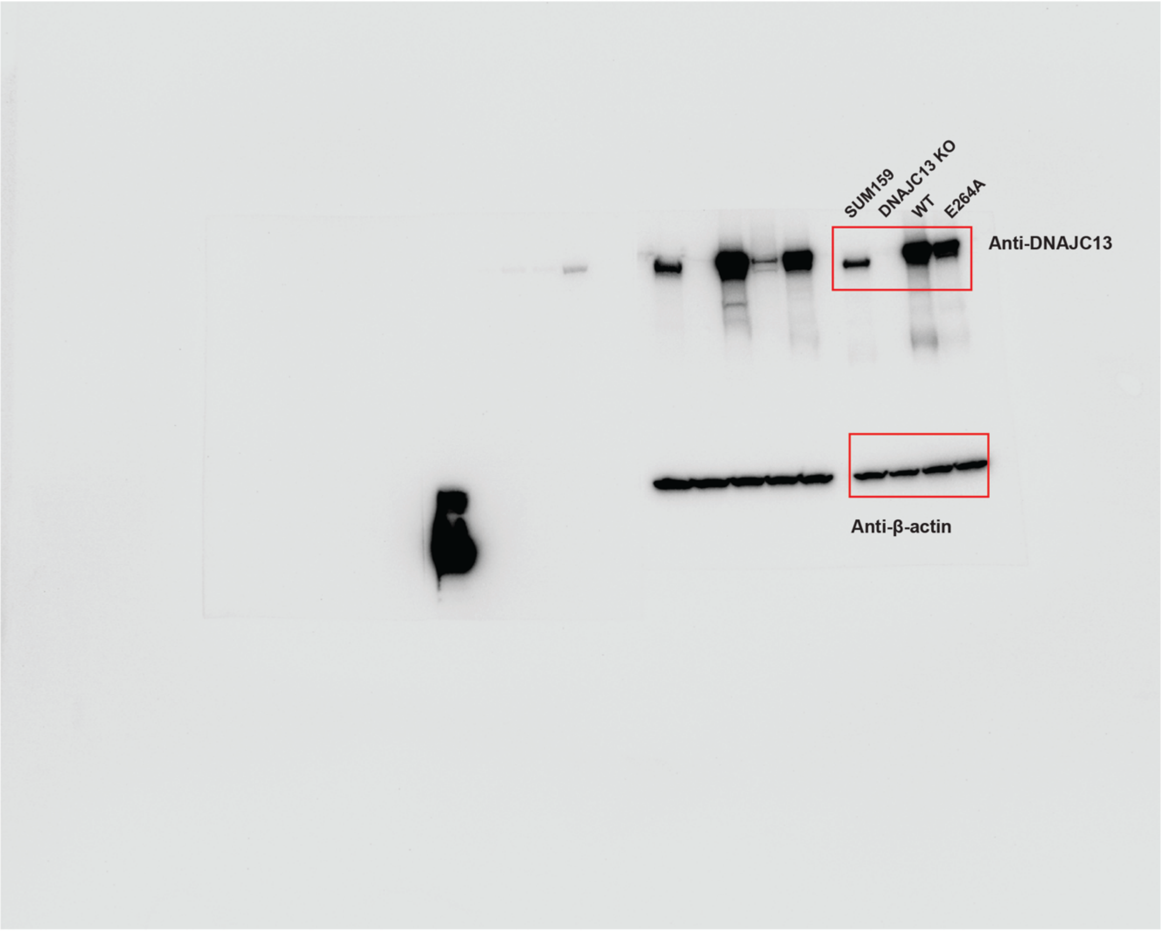

Raw images of western blots included in this study. Page 5/6

Raw data for Figure S3D

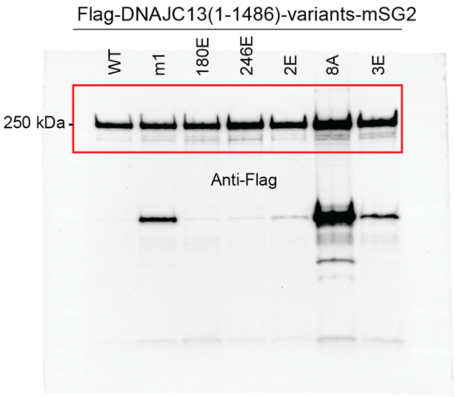


Raw data for Figure S4C

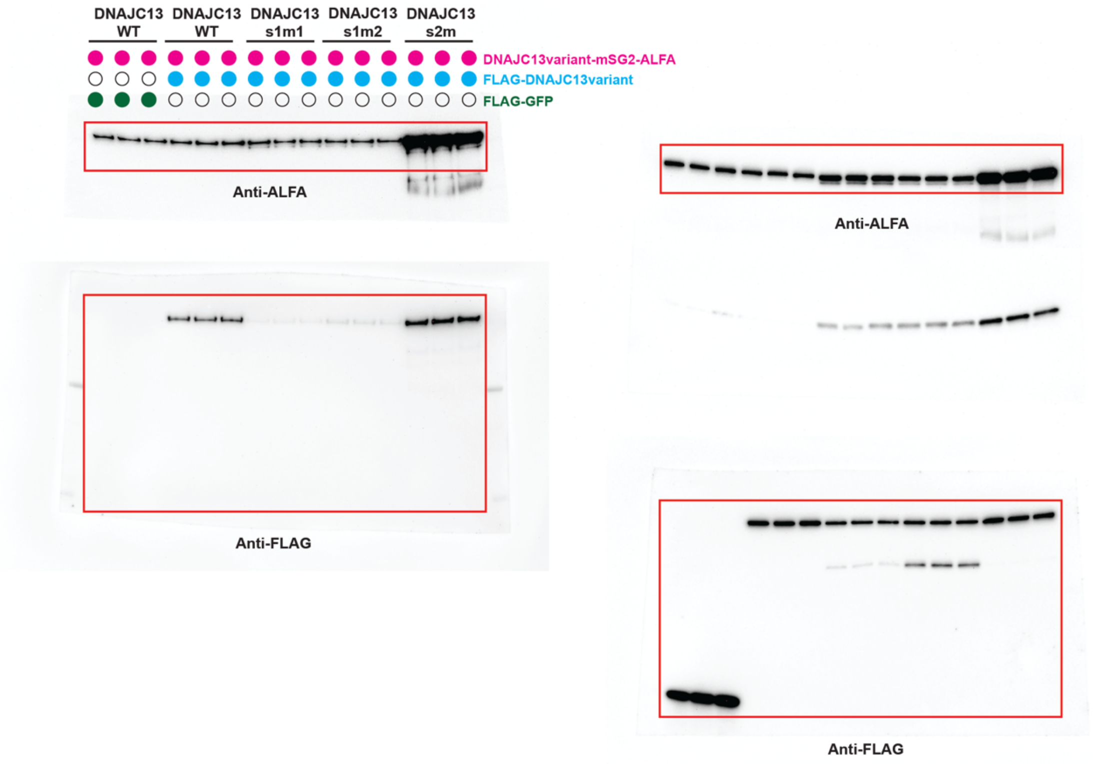

Raw images of western blots included in this study. Page 6/6

Raw data for Data S4

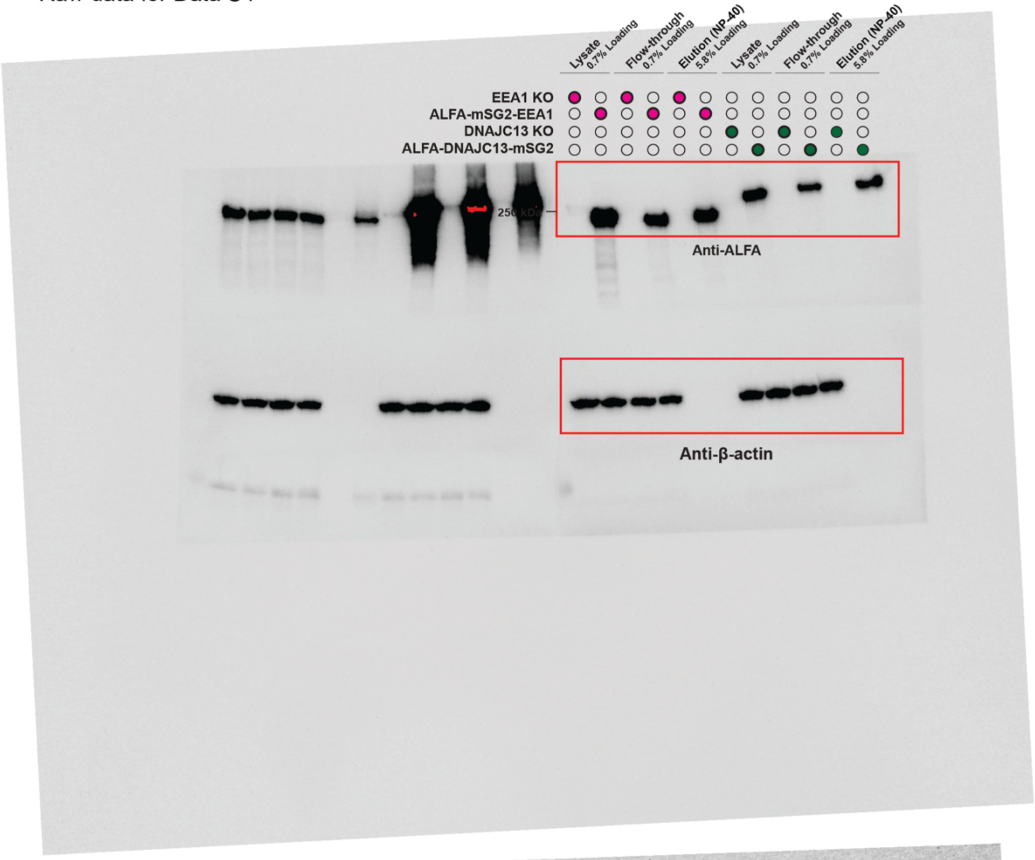


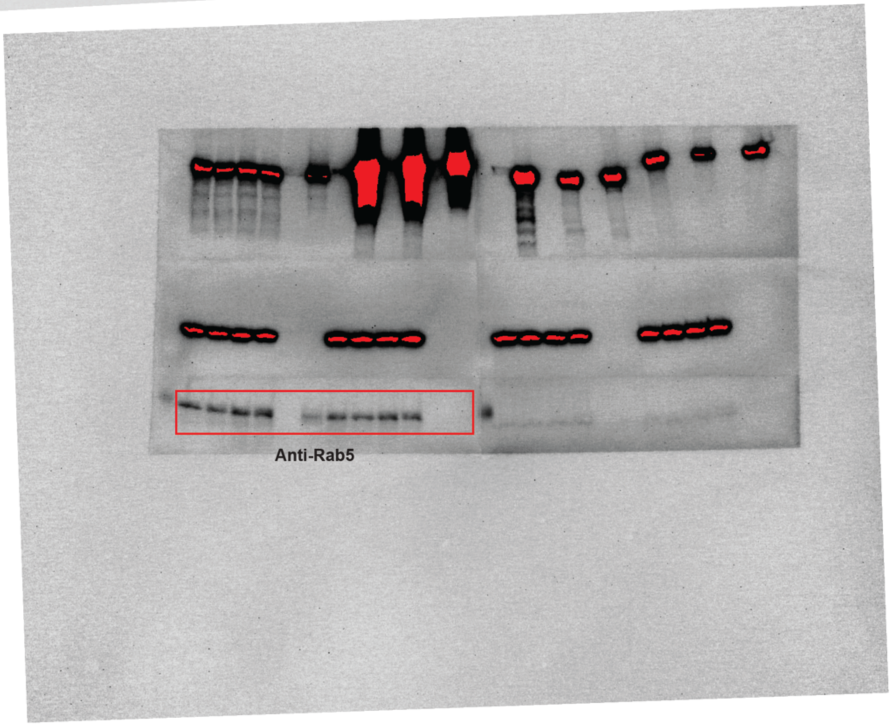
